# An ancestral signalling pathway is conserved in plant lineages forming intracellular symbioses

**DOI:** 10.1101/804591

**Authors:** Guru V. Radhakrishnan, Jean Keller, Melanie K. Rich, Tatiana Vernié, Duchesse L. Mbadinga Mbaginda, Nicolas Vigneron, Ludovic Cottret, Hélène San Clemente, Cyril Libourel, Jitender Cheema, Anna-Malin Linde, D. Magnus Eklund, Shifeng Cheng, Gane KS Wong, Ulf Lagercrantz, Fay-Wei Li, Giles E. D. Oldroyd, Pierre-Marc Delaux

## Abstract

Plants are the foundation of terrestrial ecosystems and their colonization of land was facilitated by mutualistic associations with arbuscular mycorrhizal fungi. Following that founding event, plant diversification has led to the emergence of a tremendous diversity of mutualistic symbioses with microorganisms, ranging from extracellular associations to the most intimate intracellular associations, where fungal or bacterial symbionts are hosted inside plant cells. Through analysis of 271 transcriptomes and 122 plant genomes, we demonstrate that the common symbiosis signalling pathway controlling the association with arbuscular mycorrhizal fungi and with nitrogen-fixing bacteria specifically co-evolved with intracellular endosymbioses, including ericoid and orchid mycorrhizae in angiosperms and ericoid-like associations of bryophytes. In contrast, species forming exclusively extracellular symbioses like ectomycorrhizae or associations with cyanobacteria have lost this signalling pathway. This work unifies intracellular symbioses, revealing conservation in their evolution across 450 million years of plant diversification.

## Introductory paragraph

Since they colonized land 450 million years ago, plants have been the foundation of most terrestrial ecosystems^1^. Such successful colonization occurred only once in the plant kingdom and was supported by the symbiotic association formed with arbuscular mycorrhizal fungi^2,3^. Following that founding event, plant diversification was accompanied by the emergence of alternative or additional symbionts^4^. Among alternative symbioses, the association between orchids and basidiomycetes and between Ericales and ascomycetes or basidiomycetes are two endosymbioses with specific intracellular structures in two plant lineages that lost the ability to form the Arbuscular Mycorrhizal Symbiosis (AMS)^5^. As such, orchid mycorrhiza and ericoid-mycorrhiza represent two clear symbiosis switches, whereby intracellular associations are sustained, but the nature of the symbionts are radically different. Similarly, within the liverworts, the Jungermanniales engage in ericoid-like endosymbioses but not AM symbiosis and represent another symbiont switch that occurred during plant evolution^6^. Other symbioses can occur simultaneously with AMS, for example the root nodule symbiosis, an association with nitrogen-fixing bacteria that evolved in the last common ancestor of Fabales, Fagales, Cucurbitales and Rosales^7^. Another example is ectomycorrhizae, an extracellular symbiosis found in several gymnosperm and angiosperm lineages: in some lineages both AMS and ectomycorrhizae have been retained; while other lineages have switched from AMS to ectomycorrhizae^8^. Finally, associations with cyanobacteria, which occur only in the intercellular spaces of the plant tissue, can be found in diverse species within the embryophytes, in hornworts, liverworts, ferns, gymnosperms and angiosperms^9^. Despite the improved nutrient acquisition afforded to plants by these different types of mutualistic symbioses, entire plant lineages have completely lost the symbiotic state, a phenomenon known as mutualism abandonment^4^.

Our understanding of the molecular mechanisms governing the establishment and function of these symbioses comes from forward and reverse genetics conducted in legumes and a few other angiosperms^10^ and restricted to AMS and the root-nodule symbiosis^7,11^. These detailed studies in selected plant species have allowed phylogenetic analyses to more precisely link the symbiotic genes with either AMS or root-nodule symbiosis. Indeed, the loss of AMS or the root-nodule symbiosis correlates with the loss of many genes known to be involved in these associations^7,11^. The gene losses are thought to be the result of relaxed selection following loss of the trait, resulting in co-elimination, that specifically targets genes only required for the lost trait^12^. Co-elimination can be tracked at the genome-wide level using comparative phylogenomic approaches on species with contrasting retention of the trait of interest^13,14^. Such approaches led for instance to the discovery of genes associated with small RNA biosynthesis and signalling^13^ or cilia function^15^. Applied to the AMS, comparative phylogenomics in angiosperms identified a set of more than 100 genes that were lost in a convergent manner in lineages that lost the AMS^16–18^. All classes of functions essential for AMS were detected among these genes, including the initial signalling pathway, essential for the host plant to activate its symbiotic program, and genes involved in the transfer of lipids from the host plant to the AM fungi. Since their identification by phylogenomics, novel candidates were validated for their involvement in the AM symbiosis through reverse genetic analyses in legumes^19–21^.

Targeted phylogenetic analyses have identified multiple symbiotic genes in the transcriptomes of bryophytes, but study on the overall molecular conservation of symbiotic mechanisms in land plants are lacking^22–24^. Similarly, knowledge on the plant molecular mechanisms behind the diverse array of mutualistic associations, either intracellular or extracellular, are poorly understood^10^. Here, we demonstrate through analysis of a comprehensive set of plant genomes and transcriptomes, the loss and conservation of symbiotic genes associated with the evolution of diverse mutualistic symbioses in plants.

## RESULTS AND DISCUSSION

### A database covering the diversity of plant lineages and symbiotic associations

Genomic and transcriptomic data are scattered between public repositories, specialized databases and personal websites. To facilitate large scale phylogenetic analysis, we compiled resources for species covering the broad diversity of plants and symbiotic status in a centralized database, SymDB (www.polebio.lrsv.ups-tlse.fr/symdb/). This sampling of available resources covers lineages forming most of the known mutualistic associations in plants (Supplementary Table 1), including the AM symbiosis, root nodule symbiosis, ectomycorrhizae, orchid mycorrhiza, cyanobacterial associations in hornworts and ferns, and ericoid-like symbioses in liverworts. SymDB also includes genomes of lineages that have abandoned mutualism in the angiosperms, gymnosperms, monilophytes, and bryophytes. To enrich this sampling, we generated two additional datasets: an *in depth* transcriptome of the liverwort *Blasia pusilla* that associates with cyanobacteria and since no genome of an AM-host was available for the bryophytes, we *de novo* sequenced the genome of the complex thalloid liverwort *Marchantia paleacea* which specifically associates with AM fungi^25^. The obtained assembly was of similar size and completeness to the *Marchantia polymorpha* TAK1 genome^26^ (Table 1). These new datasets augmented the SymDB database encompassing a total of 125 genomes and 271 transcriptomes that provided broad coverage of mutualistic symbioses in plants.

**Table 1:** Genome assembly statistics for Marchantia species sequenced as part of this study and comparison to the *Marchantia polymorpha* ssp. *ruderalis* TAK1 reference genome.

|  |  | *Marchantia polymorpha* ssp.<br>*ruderalis* TAK-1 | *Marchantia*<br>*paleacea* | *Marchantia*<br>*polymorpha*<br>ssp.<br>*polymorpha* | *Marchantia*<br>*polymorpha*<br>ssp.<br>*montivagans* |
| --- | --- | --- | --- | --- | --- |
| **Assembly size (Mb)** |  | 210.6 | 238.61 | 222.7 | 225.7 |
| **Scaffolds** |  | 2957 | 22669 | 2741 | 2710 |
| **N50 length (Kb)** |  | 1313.57 | 77.78 | 368.25 | 589.42 |
| **BUSCO score** | **Complete** | 821 | 817 | 855 | 855 |
|  | **Single Copy** | 793 | 790 | 832 | 829 |
|  | **Duplicated** | 28 | 27 | 23 | 26 |
|  | **Fragmented** | 48 | 53 | 38 | 42 |
|  | **Missing** | 571 | 570 | 547 | 543 |
| **G+C (%)** |  | 41.1 | 40.3 | 42.2 | 42.1 |
| **Reference** |  | Bowman *et al.*<sup>26</sup> | This study | This study | This study |

### Mutualism abandonment leads to gene loss, positive selection or pseudogenization of symbiosis genes

Previous studies have demonstrated that loss of AMS in six angiosperm lineages is associated with the convergent loss of many genes^16,17^. SymDB contains species from across the entire land plant lineage that have lost AMS and thus provided us with a platform to assess co-elimination of genes associated with the abandonment of mutualism throughout the plant kingdom. We generated phylogenetic trees for all the genes identified previously as being lost in angiosperms with loss of AMS (Supplementary Fig. 1-32 and Supplementary Table 2). Among these gene phylogenies, those missing from most lineages that have abandoned mutualism were selected. Six genes, *SYMRK*, *CCaMK*, *CYCLOPS*, the GRAS transcription factor RAD1 and two half ABCG transporters STR and STR2 were consistently lost in non-mutualistic lineages in angiosperms, gymnosperms, ferns and Bryophytes (Fig. 1; Supplementary Fig. 1,3,13–15). Very few exceptions to this trend were found (Fig. 1; Supplementary Fig. 13–14), for instance the presence of *CCaMK* in the aquatic angiosperm *Nelumbo nucifera*, that was previously reported in Bravo *et al*^16^. However, further analysis of this locus revealed a deletion in the kinase domain leading to a likely non-functional pseudogene (Supplementary Fig. 33). The same deletion was present in two different ecotypes and three independent genome assemblies (Supplementary Fig. 33). The second significant exception was in mosses, where *CCaMK* and *CYCLOPS* were present despite the documented loss of AMS in this lineage (Fig. 1; Supplementary Fig. 13–14). Previously, it was proposed that the selection acting on both genes was relaxed in the branch following the divergence of the only mycorrhizae-forming moss (*Takakia*) and other moss species^24^. Using two independent approaches (RELAX and PAML) we confirmed this initial result and identified sites under positive selection (Supplementary Fig. 34–35; Supplementary Table 3), suggesting the neofunctionalization of these two genes. From our analysis we also found conservation of STR2 (in a single fern species) and RAD1 (in two liverworts) that are thought to be non-mutualistic. The presence of these genes may reflect additional cases of species-specific neofunctionalization or may be the result of misassignment of the symbiotic state^27^.

**Figure 1.**
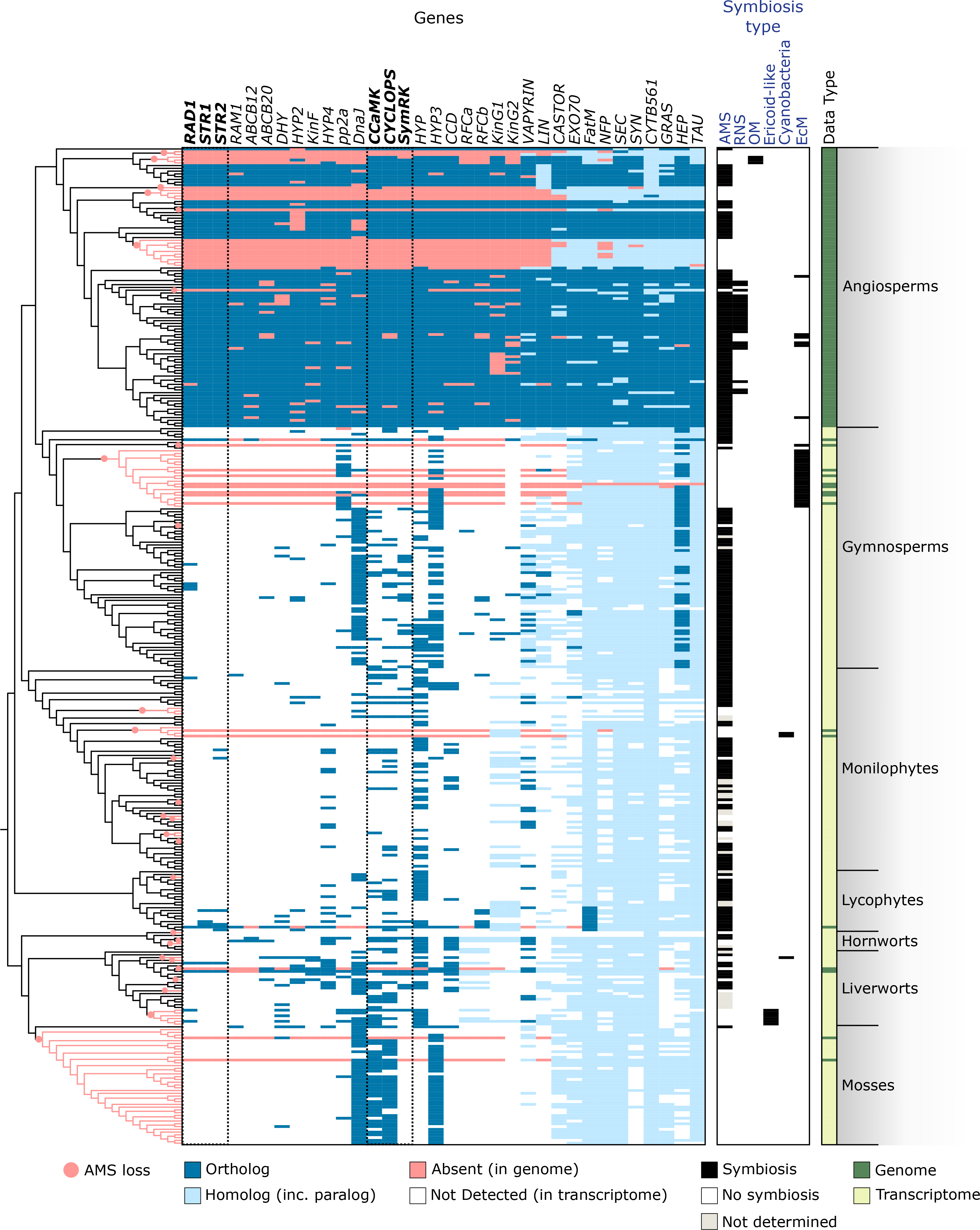
Conservation of the symbiotic genes in land plants. The tree on the left depicts the theoretical plant phylogeny. The heatmap indicates the phylogenetic pattern for each of the 34 investigated genes. The type of symbiosis formed by each investigated species is indicated by blue boxes. AMS: Arbuscular Mycorrhiza Symbiosis; RNS: Root Nodule Symbiosis; OM: Orchid Mycorrhiza; Ericoid-like; Cyanobacteria: association with cyanobacteria; EcM: EctoMycorrhizae.

Among the species that have abandoned mutualism, *M. polymorpha* is a particularly intriguing case, since the non-mutualistic *M. polymorpha* and the mutualistic *M. paleacea* belong to recently diverged lineages^28^. *M. polymorpha* represents the most recent loss of mutualism of which we are aware and hence this species may allow us to witness the process of co-elimination of symbiosis genes with the loss of mutualism. For this reason we chose to sequence the genome of the symbiotic *M. paleacea* to allow a detailed comparison between symbiotic and non-symbiotic liverwort species. Using microsynteny we identified potential remnants of symbiotic genes in *M. polymorpha* ssp. *ruderalis* TAK1, pseudogenes for *SYMRK*, *CCaMK*, *CYCLOPS* and *RAD1* existed in genomic blocks syntenic with *M. paleacea* while *STR* and *STR2* were completely absent (Fig. 2). These pseudogenes have accumulated point mutations, deletions and insertions (Fig. 2) and their presence supports a recent abandonment of mutualism in *M. polymorpha*. To better position the timing of this abandonment, we collected 35 *M. polymorpha* ssp. *ruderalis* accessions in Europe (Supplementary Table 4) and sequenced *CYCLOPS* and *CCaMK*. All accessions harboured pseudogenes at these two loci, confirming the fixation of these null alleles in the subspecies *ruderalis* (Fig. 2 and Supplementary Figure 36). Besides *M. polymorpha* ssp. *ruderalis*, two other *M. polymorpha* subspecies have been reported, ssp. *polymorpha* and ssp. *montivagans*, that are sister to *M. paleacea*. We phenotyped these two subspecies in controlled conditions, and confirmed that the loss of the AM symbiosis occurred before the radiation of the three *M. polymorpha* subspecies approximately 5 million years ago^28^ (Fig. 2). We sequenced high-quality genomes of *M. polymorpha* ssp. *montivagans* and *polymorpha* and searched for the presence of the six aforementioned genes. As for *M. polymorpha* ssp. *ruderalis* all six genes were pseudogenized or missing in these two novel assemblies (Fig. 2). As expected for genes under relaxed selection, the signatures of pseudogenization were different between the three subspecies (Fig. 2).

**Figure 2.**
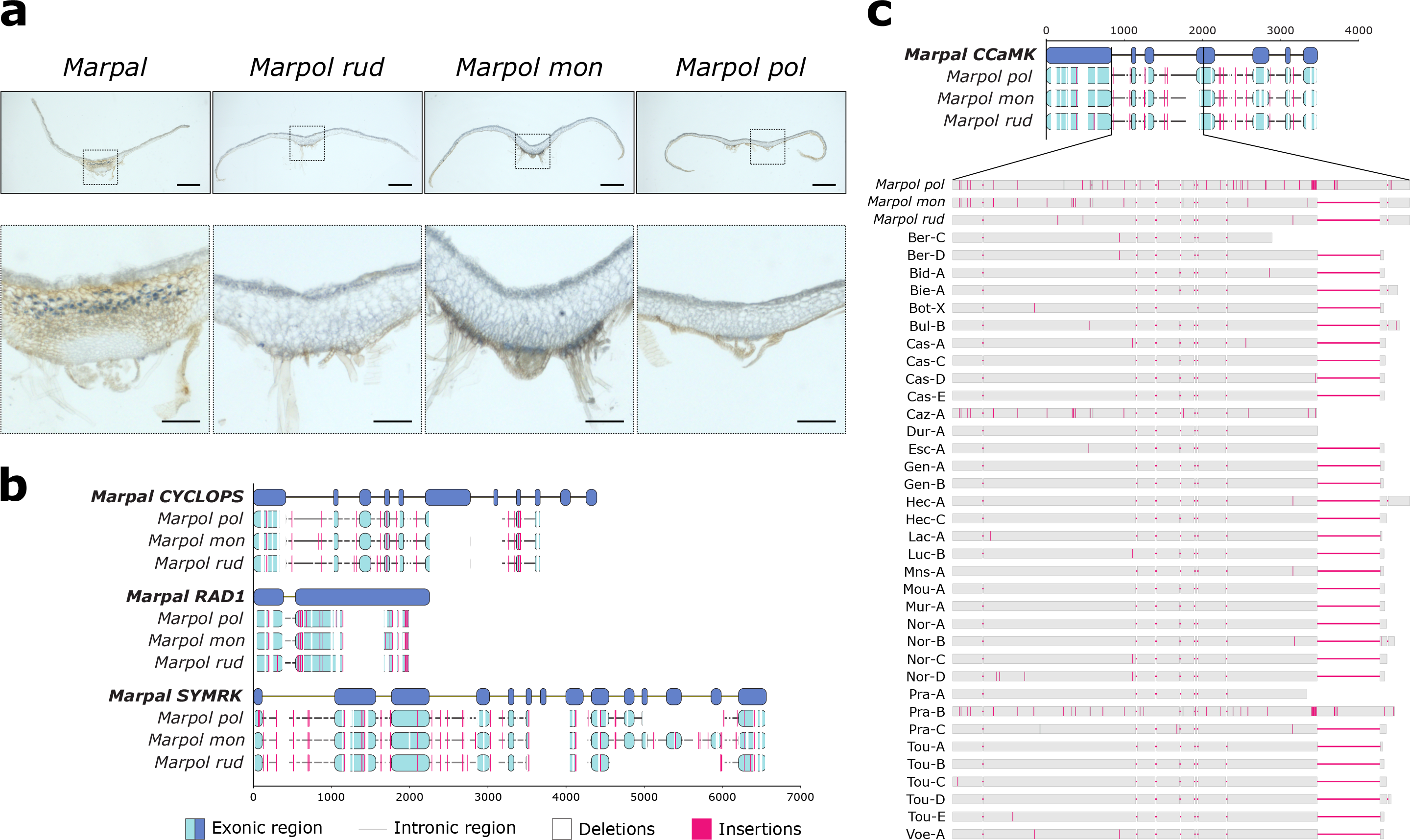
Loss of symbiotic genes following mutualism abandonment in Marchantia. **a**, Ink stained transversal sections of *Marchantia paleacea* (*Marpal*) and *Marchantia polymorpha* subspecies *ruderalis* (*Marpol rud*), *montivagans* (*Marpol mon*) and *polymorpha* (*Marpol pol*). Arbuscules are present in the midrib of *Marchantia paleacea* and absent from the three *Marchantia polymorpha* subspecies. top images, bar= 1 mm; bottom images, bar= 0.25 mm. **b**, *M. paleacea* gene models aligned with the corresponding pseudogenized loci from the three *M. polymorpha* subspecies. **c,** Multiple sequence alignment diversity in a ~1kb region of *CCaMK*. Pseudogenization pattern in 35 *Marchantia polymorpha* accessions compared to the three *Marchantia polymorpha* subspecies. Red vertical lines indicate mismatches and white boxes/red horizontal lines indicate gaps.

We conclude that mutualism abandonment leads to the consistent loss, pseudogenization or relaxed selection of at least six symbiosis-specific genes in all surveyed land plant lineages.

### Genes specific to AMS in land plants

We have shown consistent loss of six genes with mutualism abandonment, but with the broader array of genomes present we are now able to test whether these genes are lost with mutualism abandonment or specifically with the loss of AMS. Three genes, *RAD1*, *STR* and *STR2*, show a phylogenetic pattern consistent with gene loss specifically associated with the loss of AMS (Fig. 3). Our dataset covers at least 29 convergent losses of AMS, thus representing many independent replications of AMS loss in vascular plants and in bryophytes. We therefore conclude that *RAD1, STR* and *STR2* were specific to AMS in the most recent common ancestor of all land plants. The fact that all three genes are absent from non-AM host lineages indicates a particularly efficient co-elimination of these genes following the loss of AMS, suggesting possible selection against these genes^4,17,27^. We suggest that a potential driver for the loss of AMS is the adaptation to nutrient-rich ecological niches, which are known to inhibit the formation of AMS^29^ and thus render the symbiosis redundant. Alternatively, selection against these genes may be driven by the hijacking of the AMS-related pathway by pathogens that would result in positive selection acting against this pathway in the presence of a high pathogen pressure. *RAD1* has been demonstrated to act as a susceptibility factor to the oomycete pathogen *Phytophthora palmivora*^30^, providing support for this hypothesis. *RAD1* encodes a transcription factor in the GRAS family and *rad1* mutants display reduced colonization by AM fungi, defective arbuscules, the interface for nutrient exchange formed by both partners inside the plant cells and reduced expression of *STR* and *STR2*^19,31^. These two half-ABC transporters are present on the peri-arbuscular membrane and essential for the transfer of lipids from the host plant to AM fungi^20,21,32,33^. The specialization of *RAD1*, *STR* and *STR2* to AMS in all plant lineages analysed supports an ancient ancestral origin in land plants for symbiotic lipid transfer to AMS.

**Figure 3.**
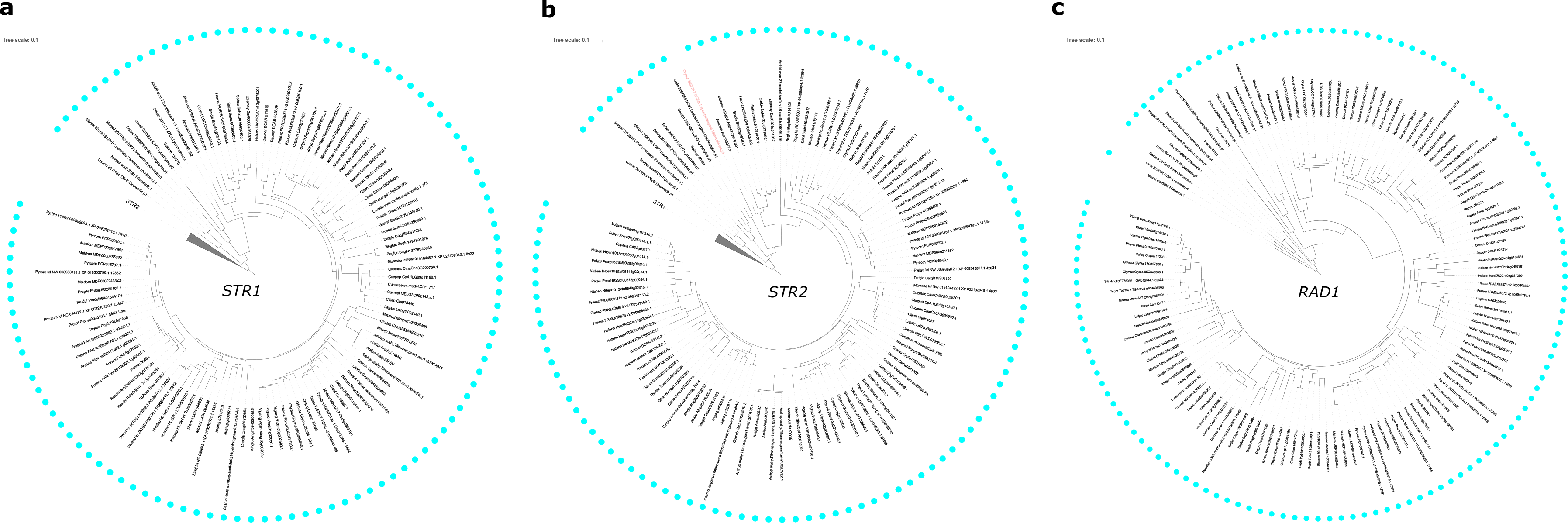
Maximum-likelihood trees of genes specific to the arbuscular mycorrhizal symbiosis in land plants. **a**, *STR1* (model: TVMe+R5); **b**, *STR2* (SYM+R6); **c**, *RAD1* (TVMe+R5). *STR1* and *STR2* trees were rooted using their closest paralogs *STR2* and *STR1* respectively. The *RAD1* tree was rooted on the bryophyte clade. Species names were coloured as follow, black: species with intracellular symbiosis; light red: species without intracellular infection; light grey: species with undetermined symbiotic status. Cyan dots indicate species forming the arbuscular mycorrhizal symbiosis.

### The symbiotic signalling pathway is conserved in species with intracellular symbioses

In contrast to *RAD1*, *STR* and *STR2*, the symbiosis signalling genes *CCaMK*, *CYCLOPS* and *SYMRK* are not absent from all species that have lost AMS. To understand this mixed phylogenetic pattern, we investigated their conservation across species with diverse symbioses. *CCaMK*, *CYCLOPS* and *SYMRK* were absent from seven genomes and fourteen transcriptomes of Pinaceae, that form ectomycorrhizae, but not AMS. None of these genes were detected in the genome of the fern *Azolla filiculoides* or in the transcriptome of the liverwort *B. pusilla*, that have independently evolved associations with nitrogen-fixing cyanobacteria, but lost AMS^9^ (Fig. 4, Supplementary Fig. 13–15). During ectomycorrhizae, the symbiotic fungi colonize the intercellular space between epidermal cells and the first layer of cortical cells^5^. Similarly, in both *A. filiculides* and *B. pusilla*, nitrogen-fixing cyanobacteria are hosted in specific glands, but outside plant cells^34,35^. Therefore, all the lineages in our sampling that host fungal or bacterial symbionts exclusively outside their cells did not retain *SYMRK, CCaMK* and *CYCLOPS* suggesting their dispensability for extracellular symbiosis. Confirming this, knock-down analysis of *CCaMK* in poplar, which forms both AMS and ectomycorrhizae, resulted in a very subtle decrease in ectomycorrhizae, while AMS was completely aborted^36^.

**Figure 4.**
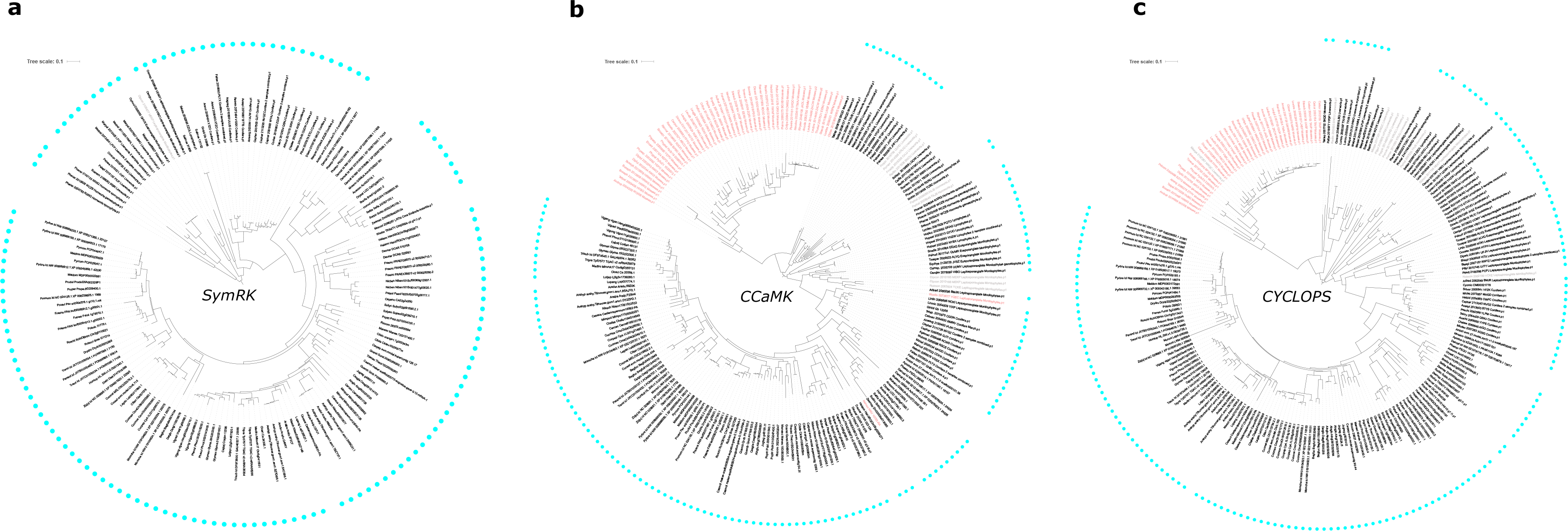
Maximum-likelihood trees of genes specific to intracellular symbiosis in land plants. **a**, *SymRK* (model: GTR+F+R5); **b**, *CCaMK* (SYM+R6); **c**, *CYCLOPS* (GTR+F+R5). Trees were rooted on the bryophyte clade. Species name were coloured as follow, black: species with intracellular symbiosis; light red: species without intracellular infection; light grey: species with undetermined symbiotic status. Cyan dots indicate species forming the arbuscular mycorrhizal symbiosis.

All other lineages that switched from AMS to other types of mutualistic symbioses retained the three signalling genes *SYMRK, CCaMK and CYCLOPS* (Fig. 4, Supplementary Fig. 13–15). These lineages are scattered throughout the land plant phylogeny, thus excluding the hypothesis of a lineage-specific retention of these genes. The three genes were found in the genomes of three Orchidaceae, *Apostasia shenzhenica*, *Dendrobium catenatum* and *Phalaenopsis equestris* and in the transcriptome of *Bletilla striata*^37^, that form Orchid Mycorrhizae with Basidiomycetes that develop intracellular pelotons^5^. All three genes were also detected in the transcriptomes of liverworts from the Jungermaniales order *Scapania nemorosa* (3/3), *Calypogeia fissa* (1/3), *Odontoschisma prostratum* (1/3), *Bazzania trilobata* (2/3) and *Schistochila sp.* (1/3), which have switched from AMS to diverse Ericoid-like associations with Basidiomycetes and Ascomycetes that form intracellular coils^5^. Furthermore, these genes are also conserved in the legume genus *Lupinus*, which associates with nitrogen-fixing rhizobia forming intracellular symbiosomes in root nodules, but have lost AMS^38^.

A unifying feature of these species that have preserved the symbiosis signalling pathway, but lost AMS, is their ability to engage in alternative intracellular mutualistic symbioses and we therefore suggest that these genes may be conserved specifically with intracellular symbioses throughout the plant kingdom. To test this hypothesis, we added to our initial analysis the only known intracellular mutualistic symbiosis not covered in our sampling: ericoid mycorrhizae, which evolved before the radiation of the angiosperm family Ericaceae. In ericoid-mycorrhiza, ascomycetes colonize epidermal root cells and develop intracellular hyphal complexes^5^. We collected available transcriptomic data from six species, assembled and annotated them, and specifically searched for the presence of *SYMRK, CCaMK*, *CYCLOPS*, as well as *STR*, *STR2* and *RAD1*. Congruent with the loss of AMS, neither *RAD1*, *STR* nor STR2 were detected (Supplementary Fig. 37), but *SYMRK, CCaMK* and *CYCLOPS* were all recovered from the transcriptome of *Rhododendron fortunei* roots, but not in the other five transcriptomes derived from leaves or stems (Fig. 4, Supplementary Fig. 37).

The receptor-like kinase SYMRK, the Calcium and calmodulin dependent protein kinase CCaMK and the transcription factor CYCLOPS are known components of the common symbiosis signalling pathway and contribute successive steps in the signalling processes triggered by AM fungi and nitrogen-fixing bacteria^2,39^. Genetic analysis of *SYMRK*, *CCaMK* and *CYCLOPS* have been conducted in multiple angiosperms, including dicots from the Fabaceae, Casuarinaceae, Fagaceae, Rosaceae or Solanaceae families as well as monocots such as rice^40–43^. In all these species, defects in any of these three genes resulted in aborted, or strongly attenuated, intracellular infection by AM fungi. In addition, knock-out or knock-down in root-nodule symbiosis-forming species resulted in impaired intracellular infection by nitrogen-fixing bacteria^2,39^. Conversely, *CCaMK* knock-down in the Fabaceae *Sesbania rostrata* did not impact extracellular infection of cortical cells by nitrogen-fixing rhizobia^44^. Together with this genetic evidence, our results demonstrate that *SYMRK*, *CCaMK* and *CYCLOPS* specifically occur in species accommodating intracellular symbionts, defining a universal signalling pathway for intracellular mutualistic symbioses in plants.

### Conservation of CCaMK and CYCLOPS biochemical properties in land plants

We propose that the symbiosis signalling pathway has been co-opted for all intracellular endosymbioses in land plants and this would imply conservation of the biochemical properties of the corresponding proteins over the 450 million years of land plant evolution. To test this hypothesis, we cloned *CCaMK* from three dicots forming AMS or both AMS and root-nodule symbiosis, from two monocots forming only the AMS and from the liverwort *M. paleacea* that forms AMS in the absence of roots. Two assays were used to assess the conservation of the biochemical properties of these CCaMK orthologs. First, truncated versions that only contain the kinase domain of CCaMK (CCaMK-K) were cloned under control of a constitutive promoter. If functional, these constructs are expected to induce the expression of root-nodule symbiosis reporter genes such as *ENOD11*^45^ when overexpressed in the Fabale *Medicago truncatula* roots in absence of symbiotic bacteria, as does the *M. truncatula CCaMK-K* construct^45^. All the constructs were introduced in a *M. truncatula pENOD11:GUS* background and GUS activity monitored in the absence of symbiotic bacteria. CCaMK-K from every tested species resulted in the spontaneous activation of the *pENOD11:GUS* reporter (Fig 5). As a second test of the conservation of CCaMK, trans-complementation assays of a *M. truncatula ccamk* (*dmi3*) mutant were complemented with *CCaMK* orthologs from the above mentioned species. In the presence of symbiotic bacteria, all of the CCaMK orthologs were able to restore nodule formation and intracellular infection in the *ccamk* mutant (Fig. 5 and Supplementary Table 5).

**Figure 5.**
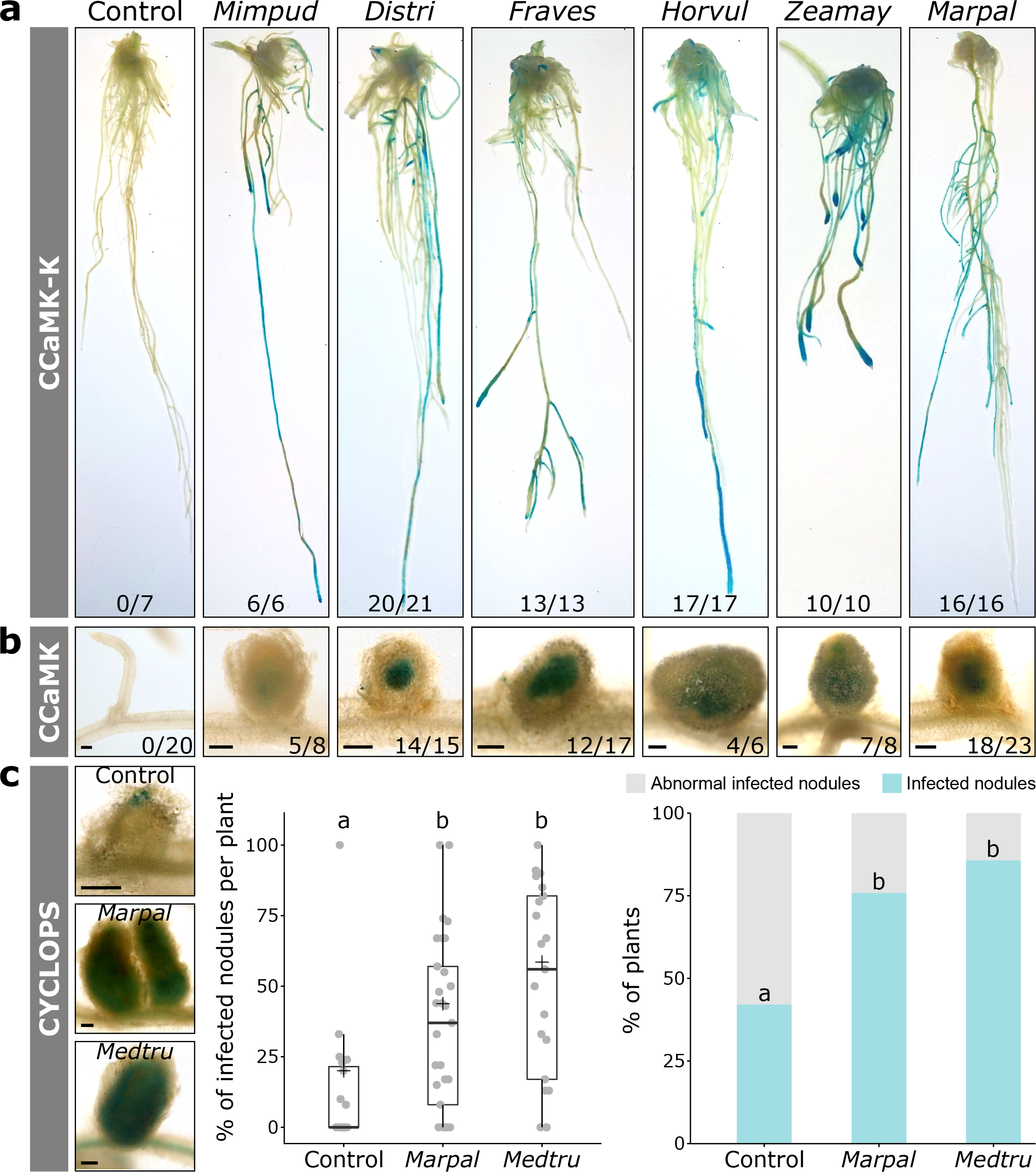
Conservation of CCaMK and CYCLOPS biochemical properties in land plants. **a,** *Medicago truncatula pENOD11:GUS* roots transformed with *pUb:CCaMK-K* from *Mimosa pudica* (*Mimpud*), *Discaria trinervis* (*Distri*), *Fragaria vesca* (*Fraves*), *Hordeum vulgarum* (*Horvul*), *Zea mays* (*Zeamay*) and *M. paleacea* (*Marpal*) show strong activation of the *ENOD11:GUS* reporter (in blue). Control roots transformed with an empty vector show little or no GUS activity. Numbers of plants showing a strong *ENOD11:GUS* activation out of the total transformed plants are indicated. **B,** *M. truncatula ccamk* mutant roots transformed with *pUb:CCaMK* from *M. pudica*, *D. trinervis*, *F. vesca*, *H. vulgarum*, *Z. mays* and *M. paleacea* show infected nodules 26 days post inoculation with *Sinorhizobium meliloti LacZ*. Bacteria in the nodules are stained in blue. A representative infected nodule is shown for each *CCaMK* ortholog. Number of plants showing infected nodules out of the total transformed plants are indicated. Scale bar 200μm. **c,** *M. truncatula cyclops* mutant roots transformed with *pUb:CYCLOPS* from *M. truncatula*, *M. paleacea* and an empty vector (control) show nodules with variable infection level. Whereas with the control plants most of the nodules are unifected or with arrested infection (as illustrated), with MedtruCYCLOPS and MarpalCYCLOPS, fully infected nodules are observed (as illustrated). The boxplot shows differences in the percentage of fully infected nodules per plant (n__control_=19, n__*Marpa*l_=29, n__*Medtru*_=21). “+” indicates mean value. Different letters indicate different statistical groups after a FDR correction at a 0.95 threshold (Kruskal-Wallis rank sum test; 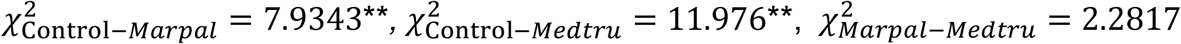). The barplot shows percentage of plants with fully infected nodules. Different letters indicate different statistical groups (Chi-Square test of independence; 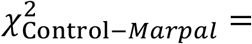 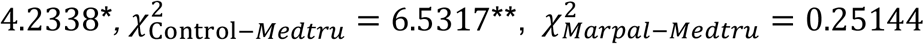).

In legumes, CCaMK phosphorylates CYCLOPS. Phosphorylated CYCLOPS then binds to the promoter and activates the transcription of downstream genes^46^. To determine whether the CCaMK-CYCLOPS module itself is biochemically conserved across land plants, we conducted trans-complementation assays of a *M. truncatula cyclops* (*ipd3*) mutant with *M. paleacea CYCLOPS*. Nodules can be formed in the *M. truncatula cyclops* mutant due to the presence of a functional paralog^47^. In our assay, *M. truncatula cyclops* mutants transformed with the empty vector could develop root nodules, but were mostly uninfected. By contrast, transformation with either *M. truncatula CYCLOPS* or *M. paleacea CYCLOPS* resulted in the formation of many fully infected nodules in most of the transformed *cyclops* roots (Fig. 5).

Altogether, these assays indicate that CYCLOPS and CCaMK orthologs that evolved in different symbiotic (AMS, root-nodule symbiosis, both) and developmental (gametophytes in *M. paleacea*, root sporophytes in angiosperms) contexts have conserved biochemical properties.

### Infectosome-related genes are conserved in angiosperms with intracellular symbioses

For a given gene with dual biological functions, co-elimination is not predicted to occur following the loss of a single trait because of the selection pressure exerted by the other, still present, trait^12^. For instance, DELLA proteins that are involved in AMS and are essential players of gibberellic-acid signalling are retained in all embryophytes^48^. To become sensitive to co-elimination, a given gene must become specific to a single trait. This may occur via the successive losses of the traits or via gene duplication leading to subfunctionalization between the two paralogs^12^. Angiosperm genomes experienced multiple rounds of whole-genome duplications^49^ and we hypothesized that, besides the common symbiotic signalling pathway, other genes might be specialized to intracellular symbioses following subfunctionalization. We screened our phylogenies for genes that are retained in angiosperm species forming intracellular symbioses but lost in those that have experienced mutualism abandonment. Six genes followed that pattern: *KinF*, *HYP*, *VAPYRIN, LIN/LIN-like, CASTOR* and *SYN* (Fig. 6, Supplementary Fig. 9, 16, 21, 22, 23 and 28). Among them, CASTOR is another component of the common symbiotic signalling pathway, while, VAPYRIN and LIN/LIN-like are directly involved in the formation of a structure required for the intracellular accommodation of rhizobial bacteria, the so called infectosome^50–52^, and also functioning in intracellular accommodation of arbuscular mycorrhizal fungi^53–55^. Similarly, *SYN* has been characterized in *M. truncatula* for its role in the formation of the intracellular structures during both AMS and root-nodule symbiosis^56^.

**Figure 6.**
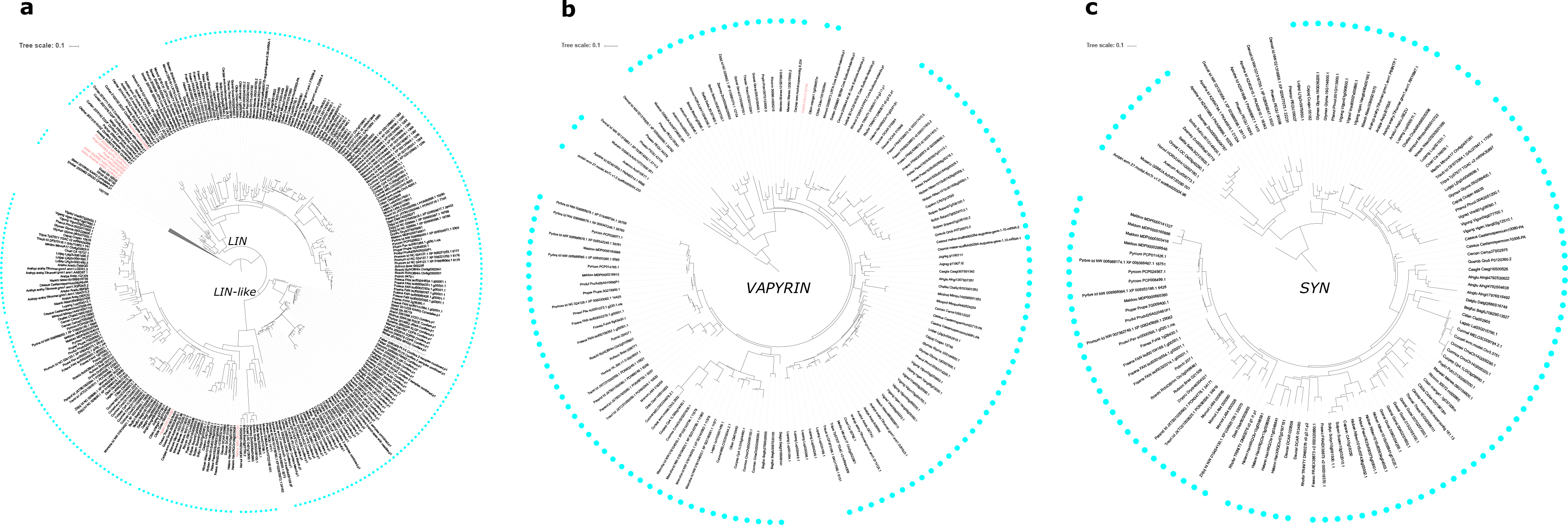
Maximum-likelihood trees of genes of the infectosome. **a**, *LIN* and its paralog *LIN-like* (model: GTR+F+R7); **b**, *VAPYRIN* (GTR+F+R6); **c**, *SYN* (TIM3+F+R5). Due to the high duplication of each families, only the angiosperms clade is displayed for *VAPYRIN* and *SYN*; whereas Gymnosperms were conserved for *LIN* and *LIN*-like due to their divergence following the seed plants whole genome duplication event. Full trees are available as Supplementary Figures 21, 22 and 28. *LIN/LIN-like* tree was rooted on non-seed plants; whereas *VAPYRIN* and *SYN* trees were rooted on *Amborella trichopoda*. Species names were coloured as follow, black: species with intracellular symbiosis; light red: species without intracellular infection; light grey: species with undetermined symbiotic status. Cyan dots outside indicate species forming AMS.

These results demonstrate that, besides the common symbiotic signalling pathway, genes directly involved in the intracellular accommodation of symbionts are exclusively maintained in species that form intracellular mutualistic symbioses in angiosperms, irrespective of the type of symbiont or the plant lineage. Pro-orthologs of these genes are found in species outside the angiosperms, in both symbiotic and non-symbiotic species. Given the overall cellular and molecular conservation observed in AMS processes in land plants, we hypothesize that these genes most likely have an endosymbiotic function in species outside of the angiosperms, but the lack of gene erosion of these genes in non-angiosperms suggests that infectosome-related genes have an additional function that ensures their retention. In angiosperms we see loss of these genes concomitant with the loss of intracellular symbioses, suggesting either that their essential function is now redundant in angiosperms or is supported by gene paralogs resulting from the whole genome duplications that predate modern angiosperms.

## CONCLUDING REMARKS

Through comprehensive phylogenomics of previously unexplored plant lineages, we demonstrate that three genes are evolutionary linked to the AM symbiosis in all land plants, including two directly involved in the transfer of lipids from the host plant to AM fungi. We propose that the symbiotic transfer of lipids has been essential for the conservation of AMS in land plants. Surprisingly, we demonstrate that genes associated with symbiosis signalling are invariantly conserved in all land plant species possessing intracellular symbionts, implying the repeated recruitment of this signalling pathway, independent of the nature of the intracellular symbiont. Furthermore, we see evidence for conservation of genes associated with the formation of the cellular structure necessary for intracellular accommodation of symbionts, but this correlation is restricted to angiosperms that have duplicated many of these components. Our results provide compelling evidence for the early emergence of genes associated with accommodation of intracellular symbionts, with the onset of AMS in the earliest land plants and then recruitment and retention of these processes during symbiont switches that have occurred on many independent occasions in the 450 million years of land plant evolution (Fig. 7). Our work also suggests that mutualistic interactions involving extracellular symbionts do not utilise the same molecular machinery as intracellular symbioses, suggesting an alternative evolutionary trajectory for the emergence of ectomycorrhizal and cyanobacterial associations.

**Figure 7.**
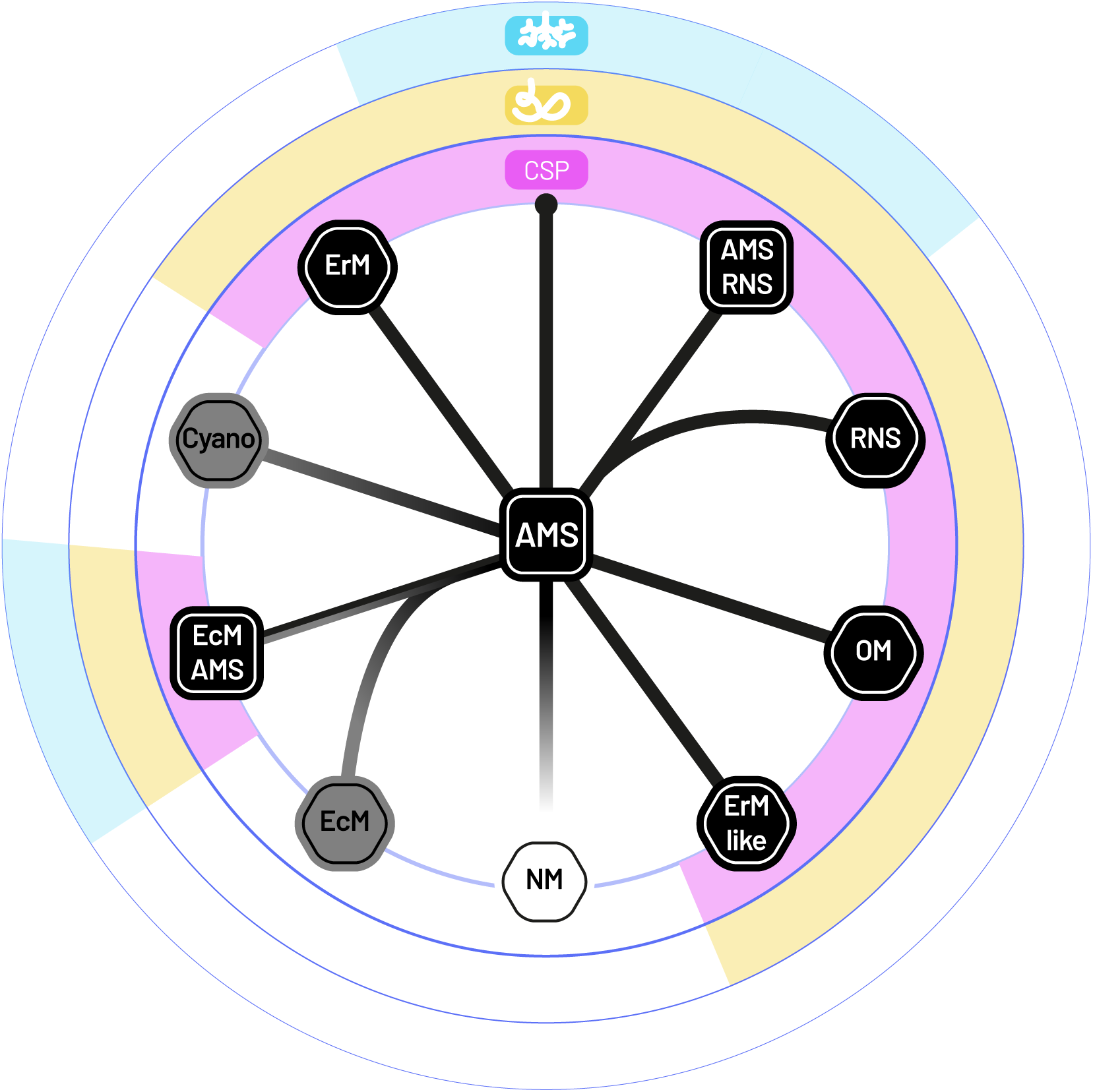
Model for the conservation of symbiotic genes across symbiosis types. *RAD1*, *STR* and *STR2* (Cyan) are exclusively conserved in species forming the Arbuscular mycorrhizal symbiosis (AMS). The Common Symbiosis Pathway genes *SYMRK*, *CCaMK* and *CYCLOPS* (CSP, magenta) in all land plants and the infectosome-related genes (*i.e. VAPYRIN*, *SYN* or *LIN/Lin-like*, in yellow) in angiosperms are specific to species forming intracellular symbiosis (black background). Mutualism abandonment (NM, white) or loss of intracellular symbiosis (grey) result in the loss of all these genes. Cyano: cyanobacteria association; EcM: EctoMycorrhizae; OM: Orchid Mycorrhiza; RNS: Root-Nodule Symbiosis; ErM: Ericoid Mycorrhiza; ErM-like: Ericoid-like Mycorrhiza.

## Methods

### Genome and transcriptome sequencing

#### Marchantia paleacea

##### Plant material

*M. paleacea* thalli previously collected in Mexico (Humphreys) were kindly provided by Dr Katie J. Field and Professor David J. Beerling (The University of Sheffield). Gemma from these thalli were collected using a micropipette tip and placed into a micro-centrifuge tube. 5% sodium hypochlorite solution was used to sterilize the gemmae for about 30 seconds followed by rinsing with sterile water 5 times to remove residual sodium hypochlorite solution. The sterilized gemmae were grown on Gamborg’s half-strength B5 medium on sterile tissue culture plates under a 16/8-hour day-night cycle at 22° C under fluorescent illumination with a light intensity of 100 μmol/μm^2^s.

##### DNA and RNA extraction

Genomic DNA from 8 week old *M. paleacea* thalli were extracted as described previously^57^. RNA extraction was also carried out from 8 week old *M. paleacea* thalli using the RNeasy mini plant kit following the manufacturer’s protocols. Fragmentation of RNA and cDNA synthesis were done using kits from New England Biolabs (Ipswich, MA) according to the manufacturer’s protocols with minor modifications. Briefly, 1μg of total RNA was used to purify mRNA on Oligo dT coupled to paramagnetic beads (NEBNext Poly(A) mRNA Magnetic Isolation Module). Purified mRNA was fragmented and eluted from the beads in one step by incubation in 2x first strand buffer at 94°C for 7 min, followed by first strand cDNA synthesis using random primed reverse transcription (NEBNext RNA First Strand Synthesis Module), followed by random primed second strand synthesis using an enzyme mixture of DNA PolI, RnaseH and E.coli DNA Ligase (NEBNext Second Strand Synthesis Module).

##### Genome sequencing

For the *M. paleacea* genome sequencing, short-insert paired-end and long-insert mate-pair libraries were produced. For the paired-end library, the DNA fragmentation was done using the Covaris (Covaris Inc., Woburn, MA). The NEBNext Ultra DNA Library Prep Kit (New England Biolabs, Ipswich, MA) was used for the library preparation, and the bead size selection for 300-400 bp based on the manufacturer’s protocol. The library had an average insert size of 336 bp. For the mate-pair library, preparation was done using the Nextera Mate Pair DNA library prep kit (Illumina, San Diego, CA) following the manufacturer’s protocol. After enzymatic fragmentation, we used gel size selection for 3-5 kb fragments. The average size that was recovered from the gel was 4311 bp. Sequencing was carried out on an Illumina HiSeq2500 on 2×100bp Rapid Run mode. The library preparation and sequencing were carried out by GENEWIZ (South Plainfield, NJ).

##### Genome assembly

Adapter and quality trimming were performed on the paired-end library using Trimmomatic v0.33 using the following parameters (ILLUMINACLIP:TruSeq2-PE.fa:2:30:10 LEADING:3 TRAILING:3 SLIDINGWINDOW:4:15 MINLEN:12). The trimmed paired-end reads were carried forward for assembling the contigs using multiple assemblers. Scaffolding of the contigs was done using the scaffolder of SOAPdenovo2^58^ with the mate-pair library after processing the reads through the NextClip^59^ pipeline to only retain predicted genuine long insert mate-pairs. Assembly completeness was measured using the BUSCO^60^ plants dataset.

##### Transcriptome sequencing

cDNA was purified and concentrated on MinElute Columns (Qiagen) and used to construct an Illumina library using the Ovation Rapid DR Multiplex System 1-96 (NuGEN, Redwood City, CA). The library was amplified using MyTaq (Bioline, London, UK) and standard Illumina TruSeq amplification primers. PCR primer and small fragments were removed by Agencourt XP bead purification. The PCR components were removed using an additional purification on Qiagen MinElute Columns. Normalisation was done using Trimmer Kit (Evrogen, Moscow, Russia). The normalized library was re-amplified using MyTaq (Bioline, London, UK) and standard Illumina TruSeq amplification primers. The normalized library was finally size selected on a LMP-Agarose gel, removing fragments smaller than 350Bp and those larger than 600Bp. Sequencing was done on an Illumina MiSeq on 2 × 300bp mode. The RNA extractions, cDNA synthesis library preparation and sequencing were carried out by LGC Genomics GmbH (Berlin, Germany).

#### Blasia pusilla

*Blasia pusilla* was originally collected from Windham County, Connecticut, USA, and maintained in Duke University greenhouse. RNA was extracted from plants with symbiotic cyanobacterial colonies using Sigma Spectrum Plant Total RNA kit. Library preparation and sequencing were done by BGI-Shenzhen. A Ribo-Zero rRNA Removal Kit was used to prepare the transcriptome library. In total, 3 libraries were constructed, which were sequenced on the Illumina Platform Hiseq2000, 150bp paired-ends, with insert size 200bp.

#### Marchantia polymorpha subspecies

##### Plant material and DNA extraction

Sterilized gemmae from one individual each of *M. polymorpha* ssp. *montivagans* (sample id MpmSA2) and *M. polymorpha* ssp. *polymorpha* (sample id MppBR5) were grown as for *M. paleacea* and isolated for DNA extraction. DNA was extracted with a modified CTAB protocol^57^.

##### Genome sequencing

DNA were sequenced with Single-molecule real-time (SMRT) sequencing technology developed by Pacific BioSciences on a PacBio Sequel System with Sequel chemistry and sequence depth of 60X^61^..

##### Genome assembly

The reads were assembled using HGAP 4^62^. Assembly statistics were assessed using QUAST^63^ version 4.5.4, BUSCO^60^ version 3.0.2 and CEGMA^64^ version 2.5.

#### OneKP transcriptome annotation

A total of 264 transcriptomes from non-angiosperm species were collected from the 1KP project (https://db.cngb.org/onekp/). For each transcriptome, ORFs were predicted using TransDecoder v5.5.0 (http://transdecoder.github.io) with default parameters. Then, predicted ORFs were subjected to BLASTp v2.7.1+^65^ and PFAM searches against the Uniprot (accessed 11/2018) and PFAM v32^66^ database. Results of BLAST and PFAM searches were used to improve final annotation and coding-sequence prediction.

### Phylogeny and sequence analysis

#### Phylogenetic analysis of candidate genes

Homologs of each candidate genes (Supplementary Table 2) were retrieved from the SymDB database with the tBLASTn algorithm v2.9.0+^65^ with e-value no greater than 1e-10 (e-value threshold was adapted for highly duplicated gene families) and using the protein sequence from *Medicago truncatula* as query. To remove short and partial transcript that are abundant in the transcriptomes from the 1KP project, shorter transcript than 60% of the *M. truncatula* query were removed prior alignment. Sequences were aligned using MAFFT v7.407^67^ with default parameter and the resulting alignments trimmed using trimAl v1.2rev57^68^ to remove all positions with more than 20% of gaps. Cleaned alignments were subjected to Maximum Likelihood (ML) approach using the IQ-TREE software v1.6.7^69^. Prior to ML analysis, best-fitted evolutionary model was tested for each alignment using ModelFinder^70^ as implemented in IQ-TREE. Branch support was tested with 10,000 replicates of SH-alrt^71^. From these initial phylogenies, clade containing the candidate genes were extracted, removing distant homologs, and a new phylogenetic analysis performed as described above. Trees were visualized and annotated using the iTOL v4.4.2^72^. Following gene phylogenies, for genes missing from sequenced genomes, the tBLASTn search was repeated on the genome assembly. This lead to the identification of a number of non-annotated genes that are summarized in Supplementary Table 6, and included in Figure 1 and Supplementary Figure 1.

#### Synteny analysis

For the genes for which orthologs/homologs were not found in in the genome and transcriptome of *M. polymorpha* ssp. *ruderalis*, more detailed searches in the genomes of *M. polymorpha* ssp. ruderalis, *M. polymorpha* ssp. polymorpha and *M. polymorpha* ssp. monitvagans were conducted. To do this, the coding sequences of the respective *M. paleacea* genes were used as queries for a BLASTn search against the genome assemblies of all three *M. polymorpha* subspecies. The genomic regions were then aligned with the corresponding *M. paleacea* genomic regions for comparisons using Mauve 2.3.1 and visualized in Geneious. Following this, gene syntenic analysis was performed by mapping the respective transcriptome onto the *M. polymorpha* ssp. *ruderalis* and *M. paleacea* genomic regions upstream and downstream of each gene using Geneious. The identities of the preceding and subsequent genes were confirmed by aligning the *M. paleacea* and *M. polymorpha* ssp. *ruderalis* sequences using MAFFT^67^. Their functional identity was confirmed using BLAST through searches against the SWISSPROT database and using Pfam.

#### Analysis of the Nelumbo CCaMK locus

Reference coding sequence of *Medicago truncatula CCaMK* was used as query to search for homolog sequence in three different *Nelumbo nucifera* genome assemblies^73,74^. Search was performed using the BLASTn+, with an e-value threshold set at 1e−10 and default parameters. For each hit in the BLAST results, 5 kb of upstream and downstream sequences were extracted using custom Python script. Extracted sequences were then aligned using MAFFT^67^ with the genomic reference and the coding sequences of *M. truncatula* and *Oryza sativa*.

#### Signature of selection on CCaMK and CYCLOPS in mosses

With the exception of *Takakia lepidozioides*, mosses are not mycorrhizal^75^. We investigated the selection acting on *CCaMK* and *CYCLOPS* on the branch before the radiation of mosses but after the divergence of *T. lepidozioides* (foreground branch). We adopted branch models^76^ to identify differential signatures of selective pressure acting on the sequences between the foreground branch and the rest of the tree (i.e. background branches) and branch-site models^77^ to identify differential signatures of selective pressures acting on specific sites between foreground and background branches. These models are implemented in the codeml program^78^ using ete-evol program^79^. In addition to the branch model, we also used the RELAX program^80^ to look for relaxation (K<1) or intensification (K>1) of the strength of selection acting on the non-mycorrhizal mosses (mosses clade without *T. lepidozioides*) compared to the rest of the tree. These methods calculate different synonymous and nonsynonymous substitution rates 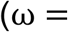 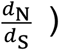 using the phylogenetic tree topology for both foreground and background branches. Because (i) the programs used do not accept gap in codon sequences and (ii) there is a negative correlation between the number of sequences and the number of ungapped positions, we used different number of sequences for each analysis. Protein sequences from CCaMK and CYCLOPS orthologs were aligned using MUSCLE v3.8.3^81^. Short sequences were excluded to maximize sequence number while limiting gapped positions compared to *CCaMK/CYCLOPS* sequences of *T. lepidozioides* using a custom R script^82^. We opted for 124 sequences for *CCaMK* and 121 sequences for *CYCLOPS* corresponding to 1092 and 552 nucleotide positions, respectively (Supplementary Table 3). We used likelihood-ratio test (LRT) to assess significance between models ran with codeml. For the branch model, we compare likelihoods from the “b_free” and “M0” models to determine if the ratios are different between background and foreground branches^76^ (*p-val*>0.05 : ω_foreground_ and ω_background_ are not different, *p-val*<0.05 : ω_foreground_ and ω_background_ are different). We also compared likelihoods from the “b_free” and “b_neut” models to determine if the ratio on the foreground branch is not neutral^76^ (*p-val*>0.05: no signature of selection on foreground (neutral), *p-val*<0.05 and ω_foreground_<1: signature of negative/relaxed selection on foreground, *p-val*<0.05 and ω_foreground_>1: signature of positive selection on foreground). For the branch-site model, we compare likelihoods from the “bsA” and “M1” models to determine if the selection pressure is relaxed on sites on the foreground branch^77^ (*p-val*<0.05: signature of relaxed selection on sites on foreground). We also compared likelihoods from the “bsA” and “bsA1” models to determine if there are sites under positive selection on the foreground branch^77^ (, *p-val*<0.05: signature of positive selection on specific sites on foreground).

### MpoCCaMK and MpoCYCLOPS pseudogenes in wild populations

#### Plant material and DNA extraction

Samples from a total of 35 accessions of *Marchantia polymorpha* (Supplementary Table 4) were collected. Approximately 0.1 g of cleaned young thalli was reduced to a fine powder in liquid nitrogen and cleaned as follow. First, 1 ml of cleaning STE buffer (Sucrose 0.25 M, Tris-HCl 0.03 M at pH=7.5 and EDTA 0.05M at pH=7.5) was added to the powder and the mixture was thoroughly vortexed. After 5 min of centrifugation at 14, 000 rpm for 5 min, the supernatant was removed. Washes with STE buffer were repeated until the red coloration was completely gone. Subsequent DNA extraction was performed using a CTAB method^57^.

#### PCR and sequencing

Putative *MpoCCaMK* and *MpoCYCLOPS* genomic sequences (1 kb) were amplified by PCR using GoTaq polymerase (Promega) according to manufacturer’s protocol. Primers used were (*MpoCCAMK* fwd GGTCATCTTGTACATTCTCCT, rev CTTGCTACTGAAGATACTTGCA; *MpoCYCLOPS* fwd CAGCACCCTCAAAACTTAGAGT, *MpoCYCLOPS* rev CCCAGCTGTTACCTTAAGAAG). After amplification, PCR products were sent for Sanger sequencing at Eurofins (Germany) using the PCR forward and reverse primers of the two genes. Sanger sequences were assembled using the Tracy software v0.5.5 and aligned using MUSCLE^81^ v3.8.3 with genomic *CCaMK* and *CYCLOPS* sequences from the three *Marchantia polymorpha* subspecies.

### Mycorrhization assays

#### Material and growth conditions

Thalli of *Marchantia paleacea*, *Marchantia polymorpha* ssp. ruderalis (TAK1), *Marchantia polymorpha* ssp. *polymorpha* (MppBR5) and *Marchantia polymorpha* ssp. *montivagans* (MpmSA2) were grown on a substrate containing 50% Oil‐Dri UK (Damolin, Etrechy, France) and 50% sand (0.7–1.3 mm) in 9×9×8 cm pots (5 thalli per pot, 5 pots per species/subspecies). Each pot was inoculated with 1000 sterile spores of *Rhizophagus irregularis* DAOM 197198 (Agronutrition Labège, France), grown at a 16h/8h photoperiod at 22°C/20°C and watered once a week with ½ Long Ashton medium containing 7.5 μM of phosphate^83^.

#### Microscopic phenotyping

After 8 weeks of colonization, thalli were harvested Sections of the thalli were made by hand or using a Leica vt1000s vibratome (100μm sections) on thalli included in 6% agarose. Sections were cleared in 10% KOH overnight and ink colored one hour in 5% Sheaffer ink, 5% acetic acid. Sections were observed under a microscope and pictures of representative sections were taken with a Zeiss Axiozoom V16 microscope.

#### Molecular phenotyping

After 8 weeks of colonization, portions of 5 thalli per condition were harvested and frozen in liquid nitrogen. DNA was extracted from frozen thalli material as described in Edwards^84^. PCR was performed using GoTaq polymerase (Promega) following manufacturer instructions on the *M. polymorpha Elongation factor-1 alpha* (Mapoly0024s0116, fwd CGACCACTGGTCACCTTATC, rev AACCTCAGTGGTCAGACCG) and on the *Rhizophagus irregularis Elongation factor-1 alpha* (Rhiir2_1_GeneCatalog_transcripts_20160502.nt, fwd TGTTGCTTTCGTCCCAATATC, rev CTCAACACACATCGGTTTGG)

### Medicago truncatula root assays

#### Plasmid construction

The Golden Gate modular cloning system was used to prepare the plasmids as described in Schiessl *et al*^85^. Levels 0 and 1 used in this study are listed in Supplemental Table 6 and held for distribution in the ENSA project core collection (https://www.ensa.ac.uk/). We used L1 plasmid piCH47811^86^, L1 construct L1M-R1-pAtUBI-DSred-t35S^23^ and L2 acceptor backbone EC50507. Sequences were domesticated (listed in Supplemental Table 7) synthesized and cloned into pMS (GeneArt, Thermo Fisher Scientific, Waltham, USA).

#### Activation of pENOD11:GUS by CCaMK orthologs

Constructs containing CCaMK-K were transformed in *A. rhizogenes* A4TC24 by electroporation. Transformed strains were grown at 28°C in Luria-Bertani medium supplemented with rifampicin and kanamycin (20 μg/mL). *M. truncatula pENOD11:GUS* L416 roots^87^, were transformed with the different CCaMK-K as described by Boisson-Dernier *et al.*^88^, and grown on Fahraeus medium for 2 months, selected with the DsRed marker present in all the constructs and GUS stained as in Vernié *et al*^89^.

#### Trans-complementation assays

The *MedtruCCaMK* null allele *dmi3-1*^90^ and *MedtruCYCLOPS* null allele *ipd3-1*^91^ were used in this study. Roots were transformed as described above with the various *CCaMK* and *CYCLOPS* constructs respectively (Supplementary Table 7). For nodulation assays, *M. truncatula* plants with DsRed positive roots were transferred in pots containing Zeolite substrate (50% fraction 1.0-2,5mm, 50% fraction 0,5-1.0-mm, Symbiom) and watered with liquid Fahraeus medium. Wild-type *S. meliloti* RCR2011 pXLGD4 (GMI6526) was grown at 28°C in tryptone yeast medium supplemented with 6 mM calcium chloride and 10 μg/mL tetracycline, rinsed with water and diluted at OD_600_=0.02. Each pot was inoculated with 10 ml of bacterial suspension. Two independent biological replicates were conducted for *ccamk* and *cyclops* complementation assays. For *ccamk*, one biological assay was harvested at 26DPI and scored for the ratio of infected nodules after X-Gal staining, and one at 42DPI scored for the total nodule number. For *cyclops*, one biological assay was harvested at 32DPI and one at 49 DPI, both were scored for infected nodules after X-Gal staining and the ratio of fully infected nodules out of the total number of nodules was calculated and used on the box plot (both biological replicates combined).

## Data availability

All assemblies and gene annotations generated in this project can be found in SymDB (www.polebio.lrsv.ups-tlse.fr/symdb/). Raw sequencing data can be found under NCBI Bioproject PRJNA576233 (*Blasia pusilla*), PRJNA362997 and PRJNA362995 (*Marchantia paleacea* genome and transcriptome respectively) and PRJNA576577 *(Marchantia polymorpha* ssp. *montivagans* and ssp. *polymorpha*).

## Code availability

## Acknowledgements

This work was supported by the ANR grant EVOLSYM (ANR-17-CE20-0006-01) to P-M.D, by the research project Engineering Nitrogen Symbiosis for Africa (ENSA), which is funded through a grant to the University of Cambridge by the Bill & Melinda Gates Foundation (OPP11772165), by the 10KP initiative (BGI-Shenzhen), by the National Science Foundation (DEB1831428) to F.-W.L and by the Swedish Research Council VR to U.L. (2011‐5609 and 2014‐522) and to D.M.E (2016-05180). G.V.R is additionally supported by a BBSRC Discovery Fellowship (BB/S011005/1). Part of this work was conducted at the LRSV laboratory, which belongs to the TULIP Laboratoire d’Excellence (LABEX) (ANR-10-LABX-41). We are grateful to the genotoul bioinformatics platform Toulouse Midi-Pyrenees for providing computing and storage resources. We thank Fabrice Roux for helping with the collection of *M. polymorpha* accessions, Aisling Cooke for assistance with *M. paleacea* DNA extraction, Peter Szoevenyi for advices on *M. polymorpha* DNA extraction, David Barker and members of the ENSA project for helpful comments and discussion. Figure 7 was prepared by Jeremy Calli (www.jeremy-calli.fr).

## Author contributions

P-M.D., G.V.R., M.K.R., J.K., G.E.D.O. and T.V. conceived the experiments; J.K, H.S.C and L.C. developed symDB; G.V.R., M.K.R., J.K., T.V., D.L.M.M., N.V., C.L., J.C., and P-M.D. conducted the experiments; A-M.L., D.M.E., U.L. generated the *M. polymorpha* subspecies genomes; F-W.L., S.C. and G.K.S.W. generated the *Blasia pusilla* transcriptome; P-M.D., G.V.R., M.K.R., J.K. and T.V. analysed the data; J.K. compiled the Supplementary material; G.V.R., M.K.R., G.E.D.O. and P-M.D. wrote the manuscript.

