## Supplementary Data for "An ancestral signalling pathway is conserved in plant lineages forming intracellular symbioses"

**Supplementary Figure 1: Detailed view of Figure 1 with species name added on terminal branches.**

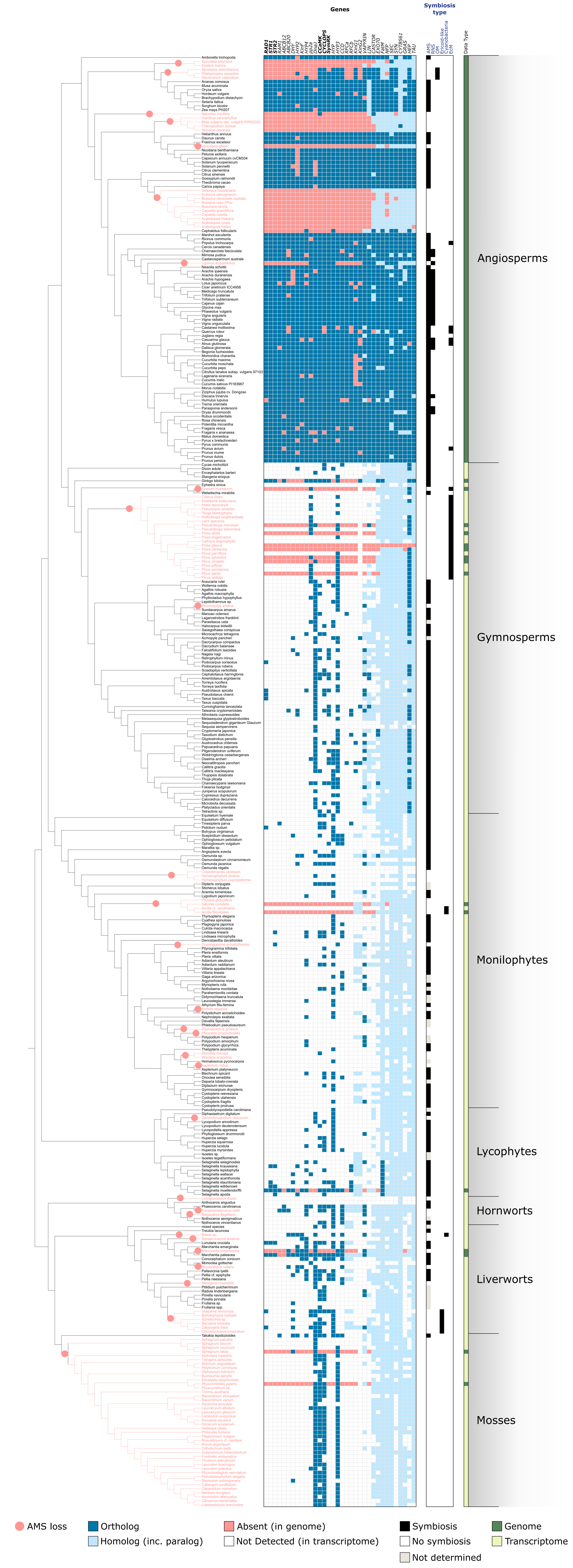

Supplementary Figure 2: Maximum likelihood tree of *RAD1*.

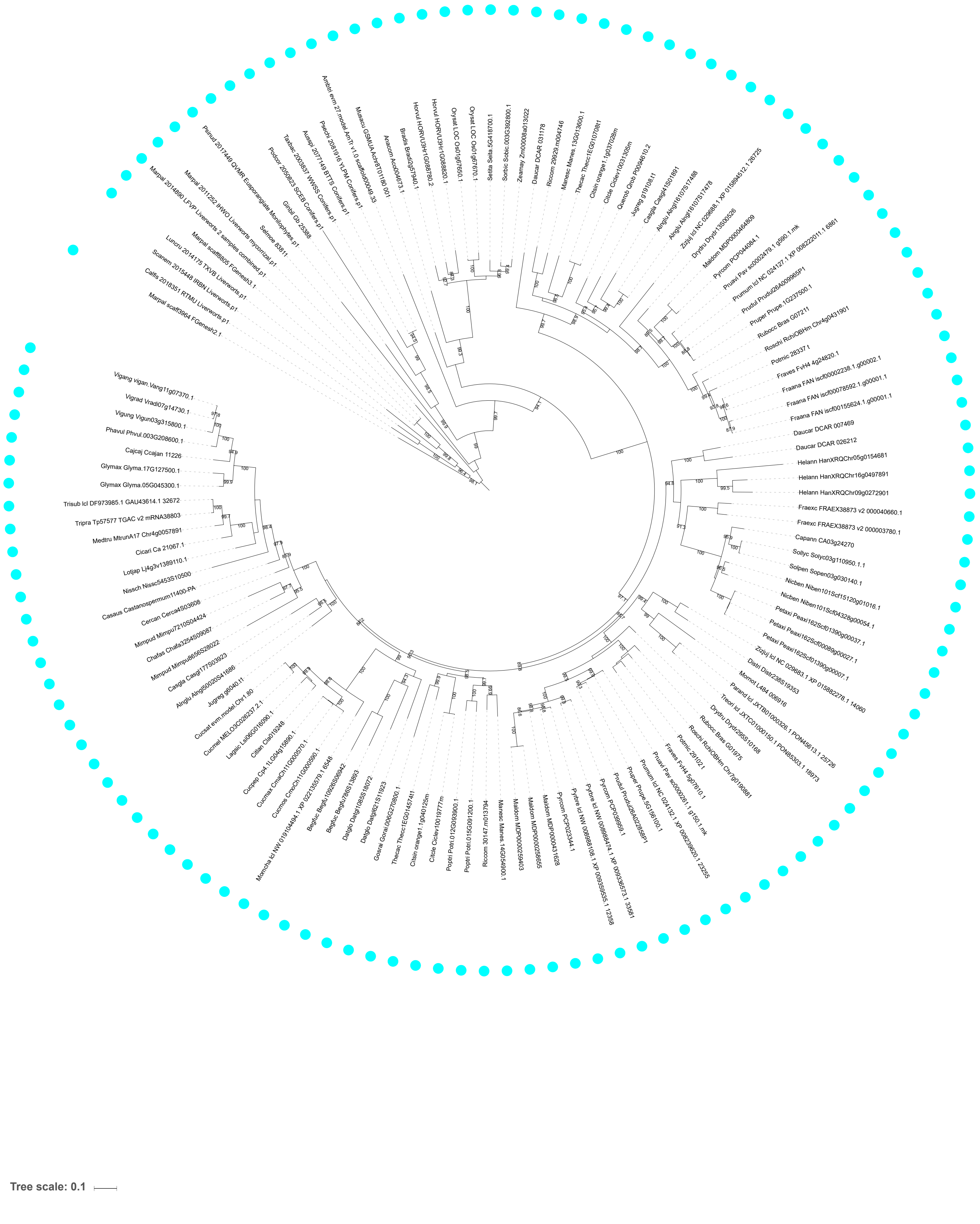

Supplementary Figure 3: Maximum likelihood tree of STR1/STR2.

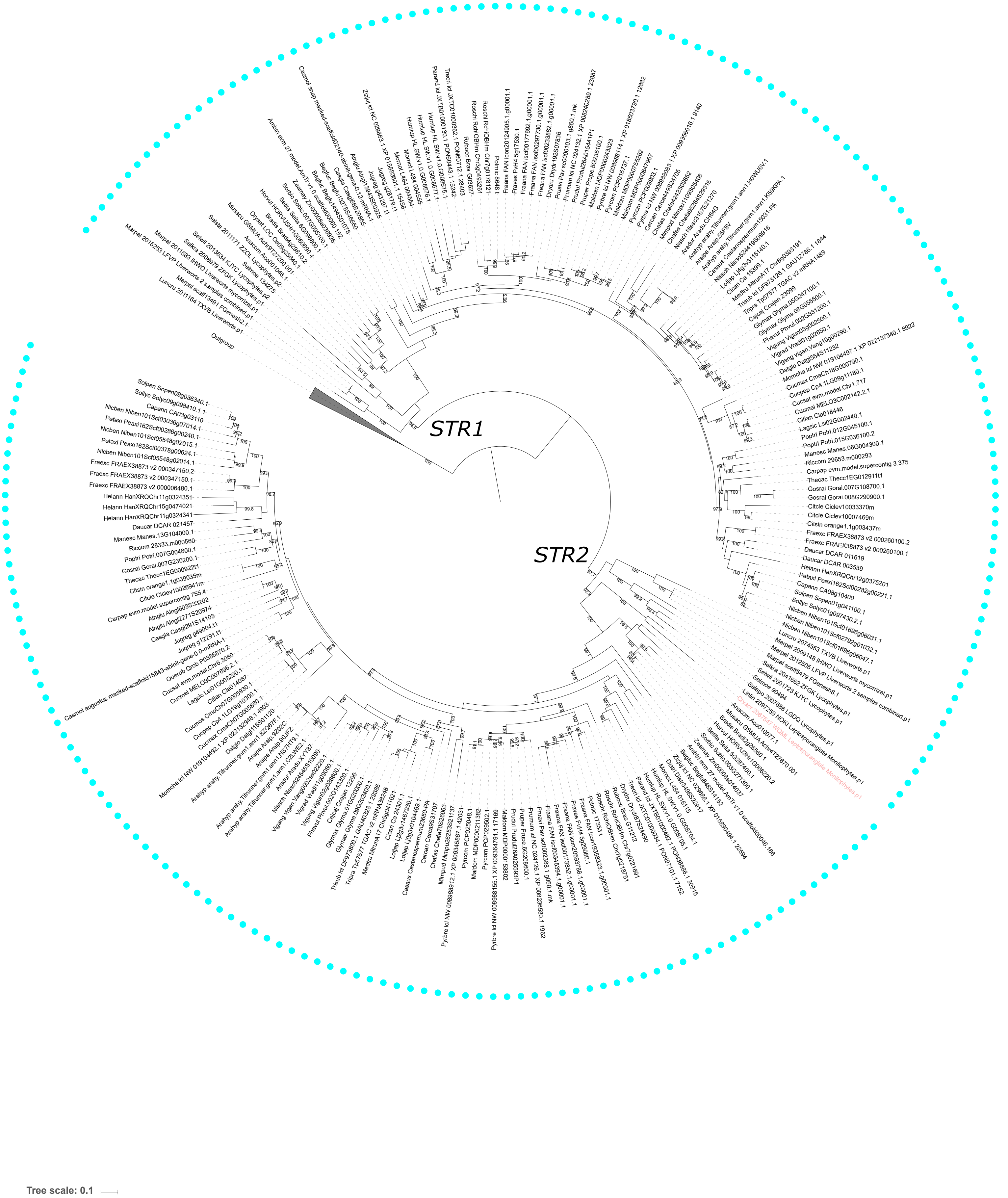

Supplementary Figure 4: Maximum likelihood tree of *RAM1*.

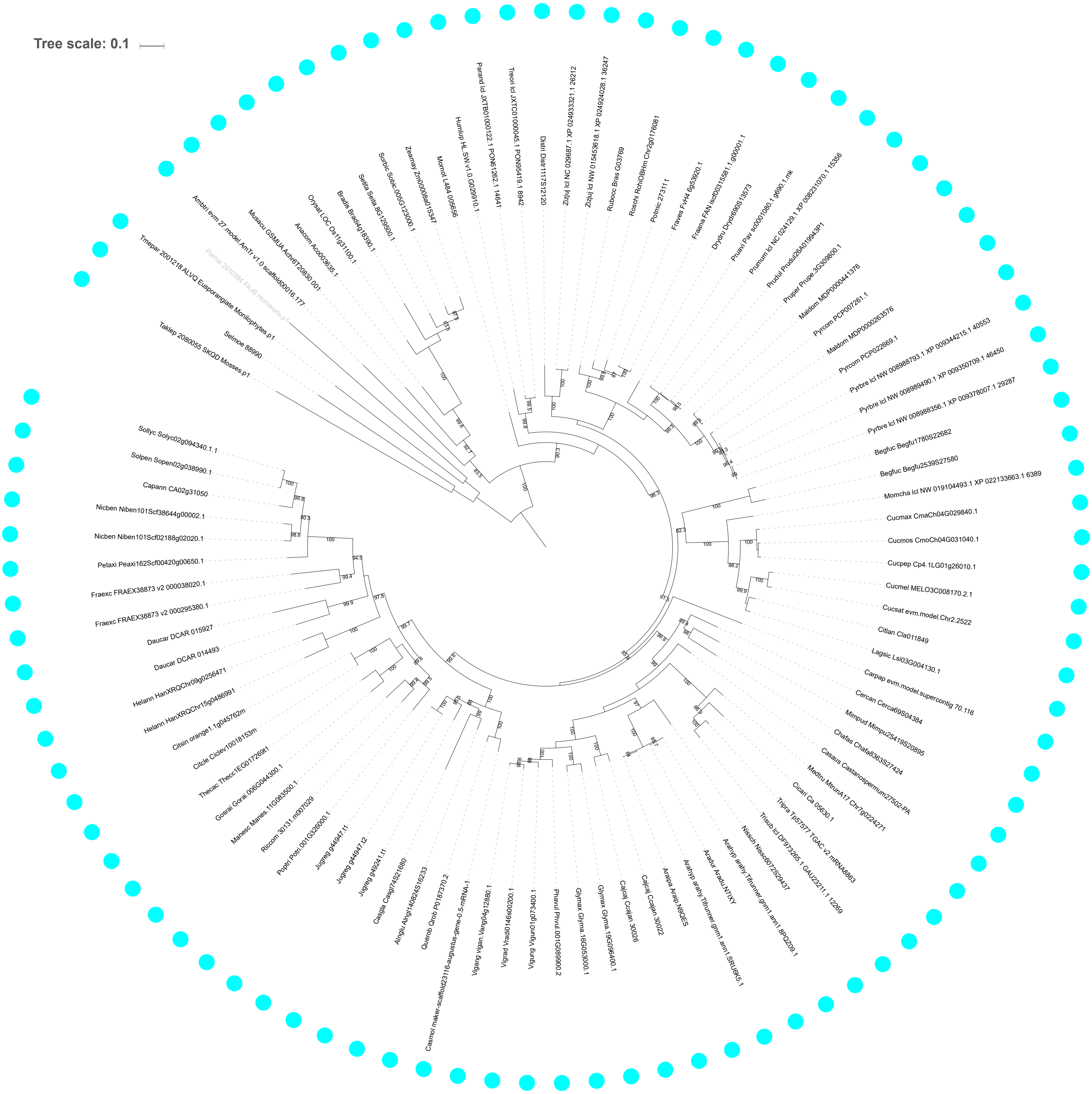

Supplementary Figure 5: Maximum likelihood tree of *ABCB12*.

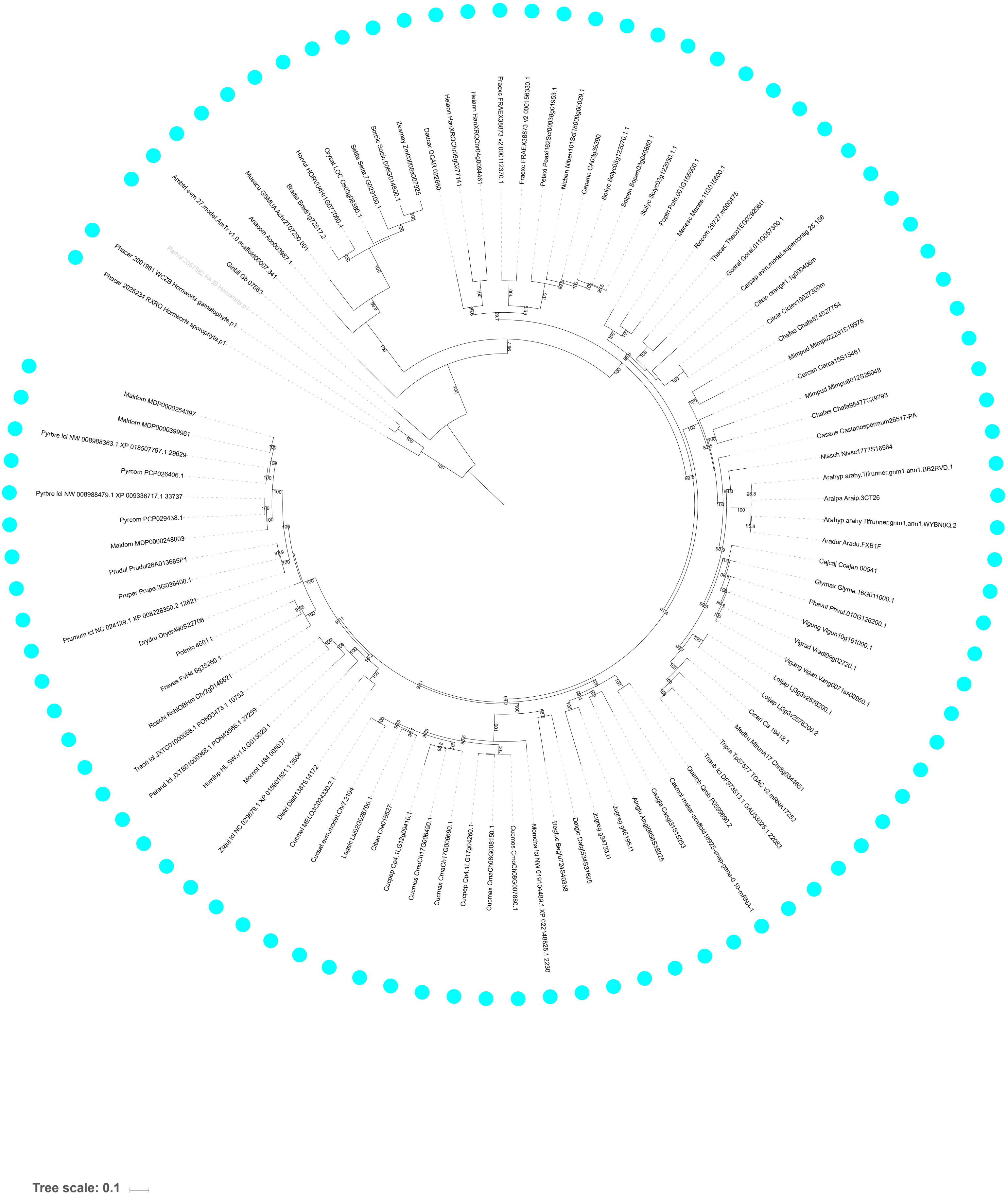

Supplementary Figure 6: Maximum likelihood tree of *ABCB20*.

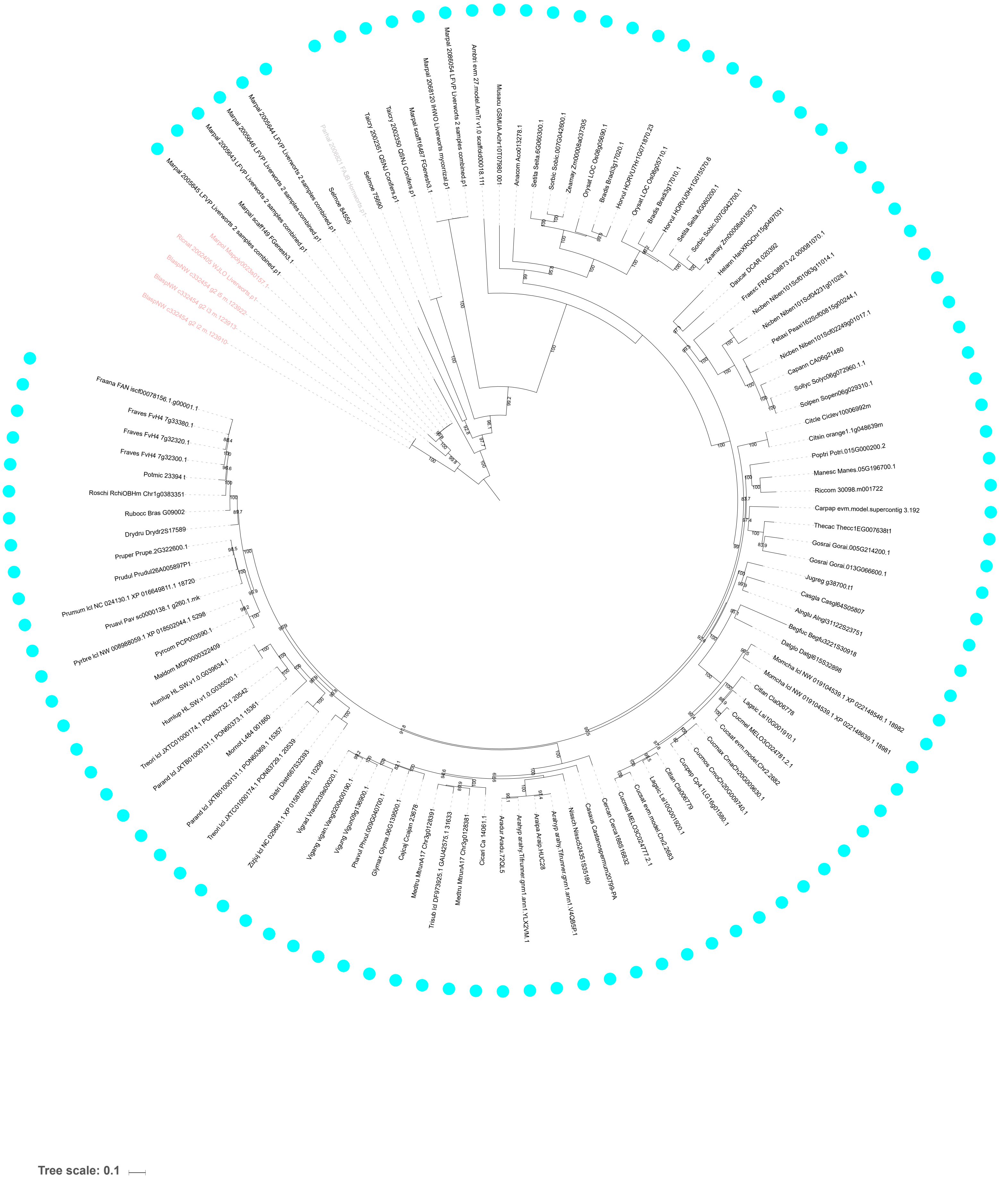

Supplementary Figure 7: Maximum likelihood tree of *DHY*.

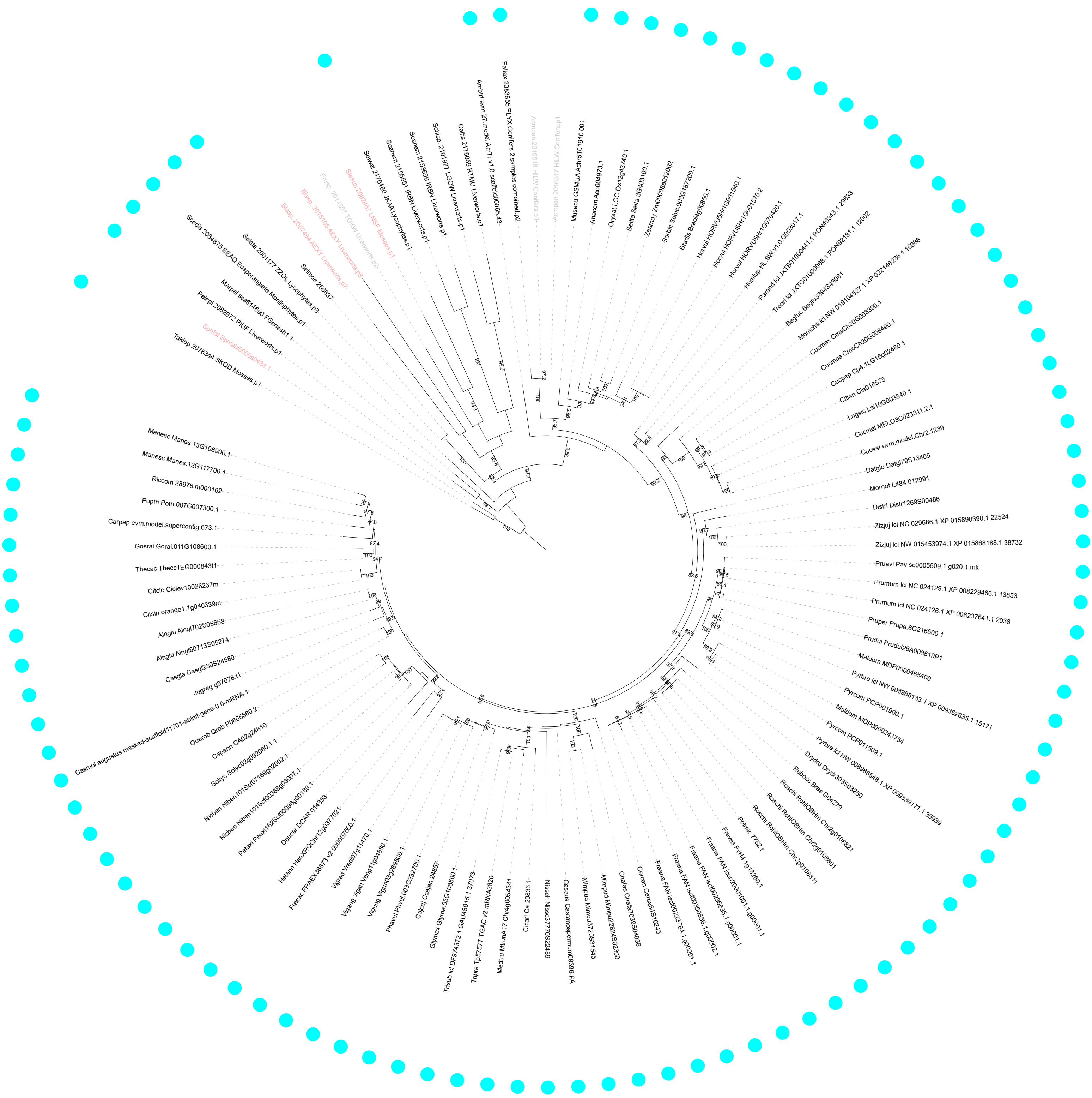

Supplementary Figure 8: Maximum likelihood tree of *HYP2*.

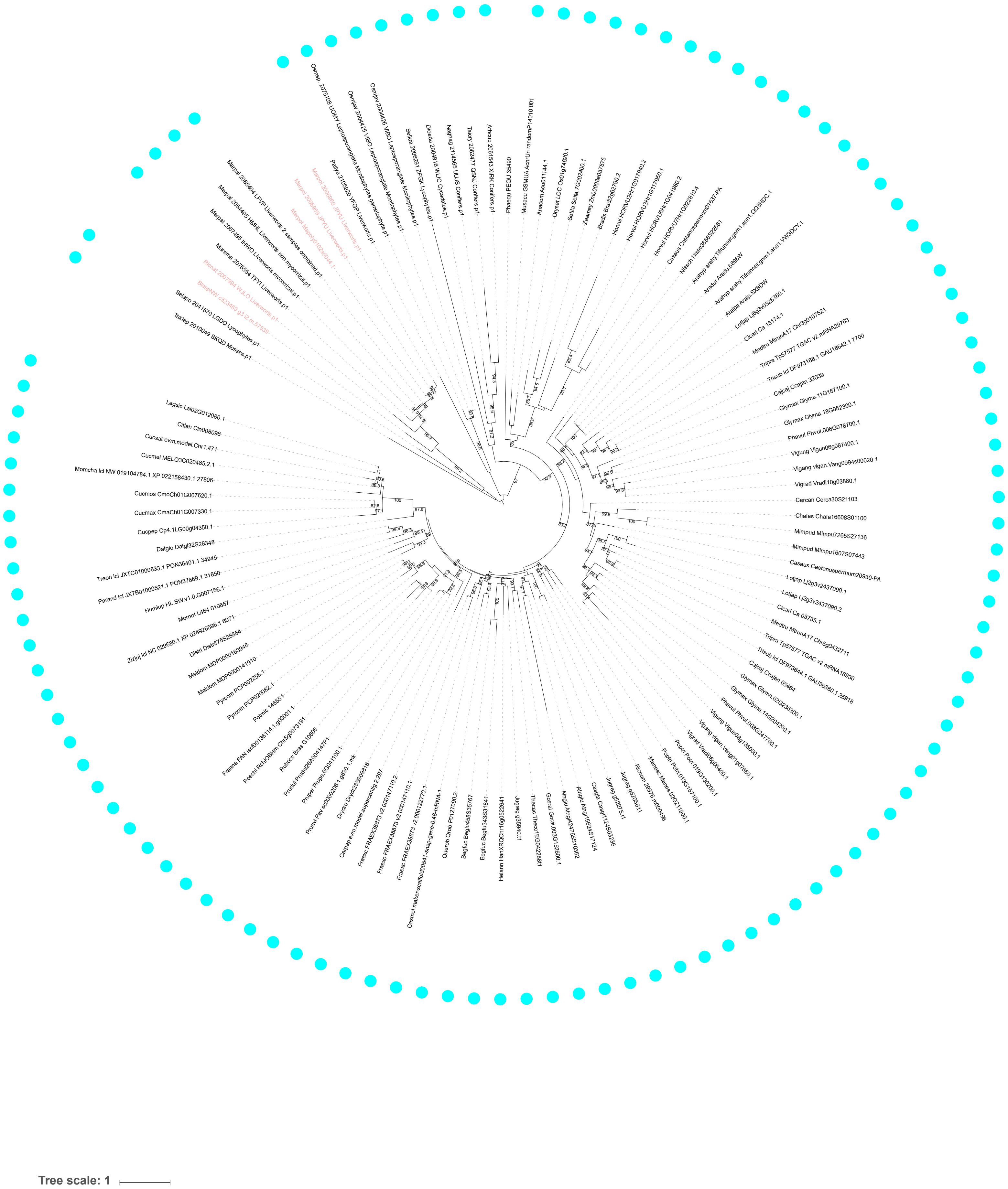

Supplementary Figure 9: Maximum likelihood tree of *KinF*.

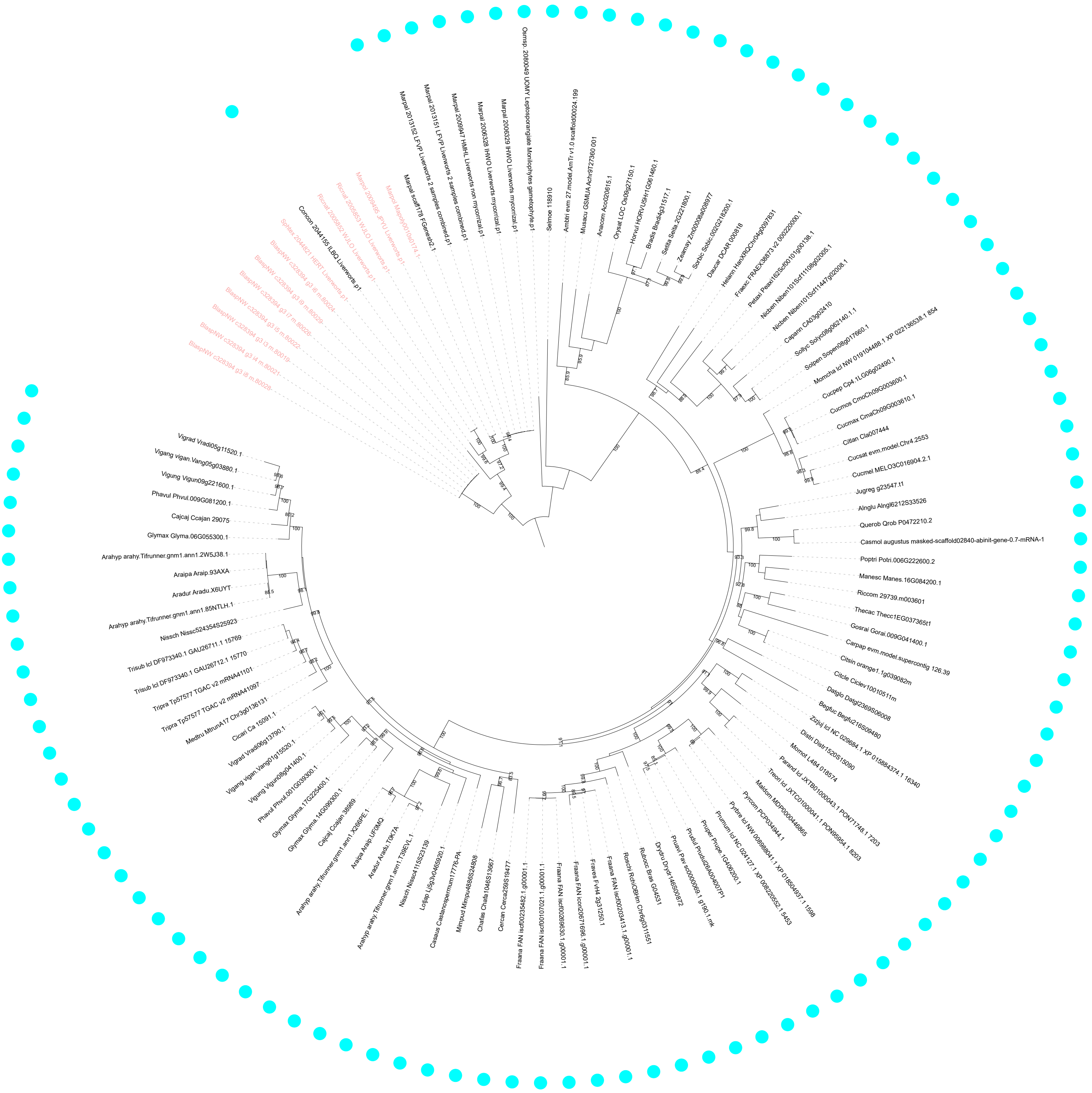

Tree scale: 0.1

Supplementary Figure 10: Maximum likelihood tree of *HYP4*.

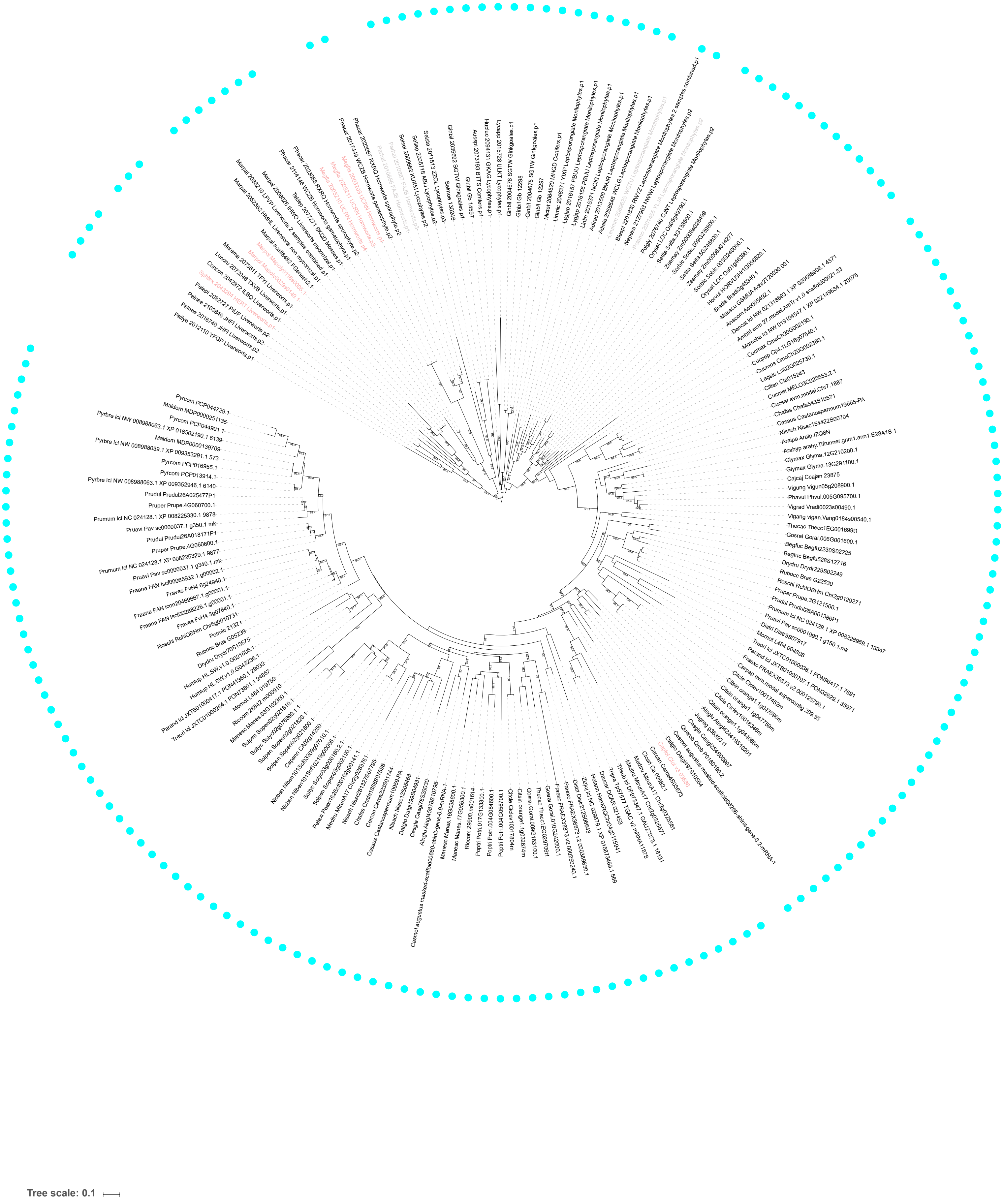

Supplementary Figure 11: Maximum likelihood tree of *pp2a*.

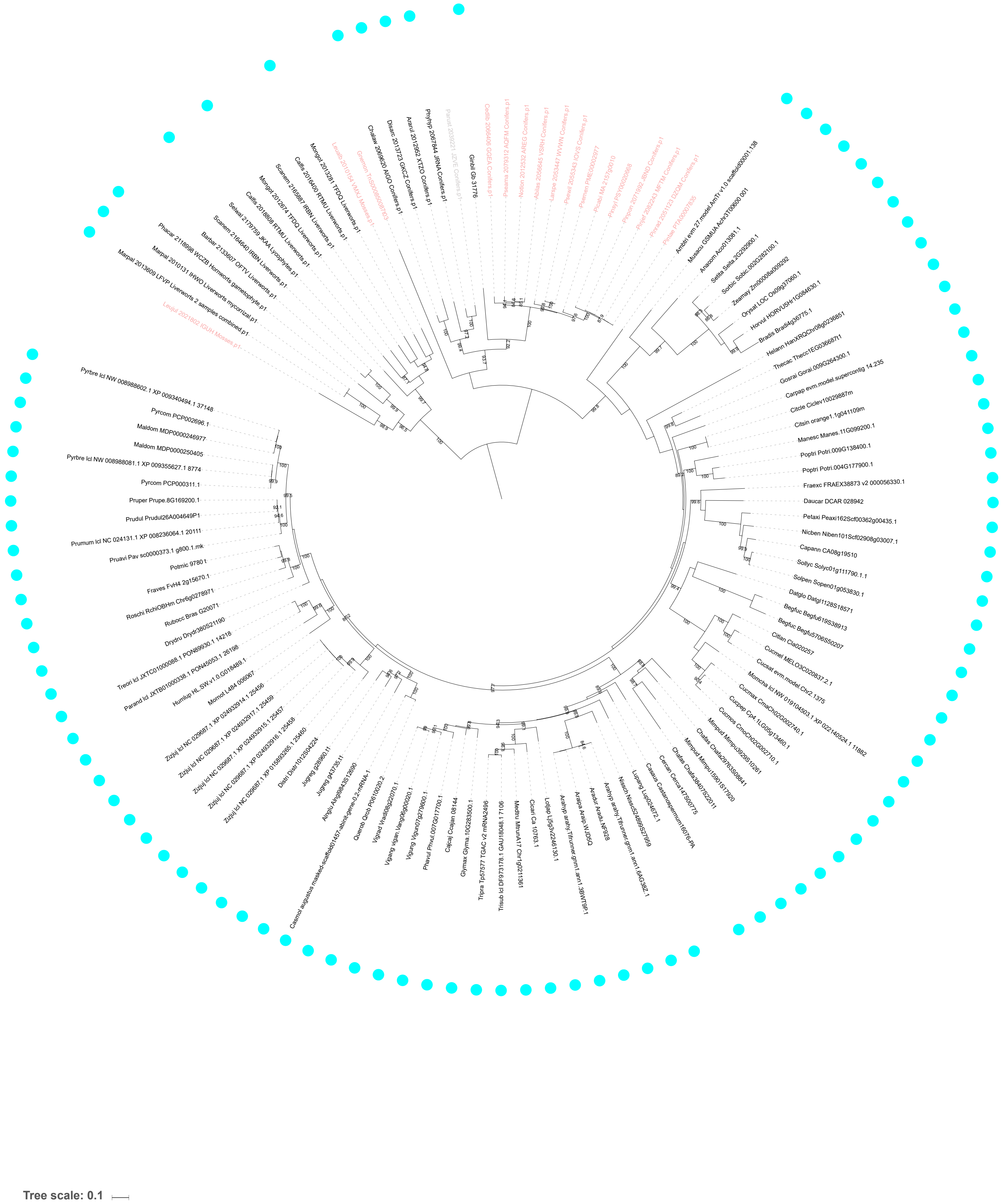

Supplementary Figure 12: Maximum likelihood tree of *DnaJ*.

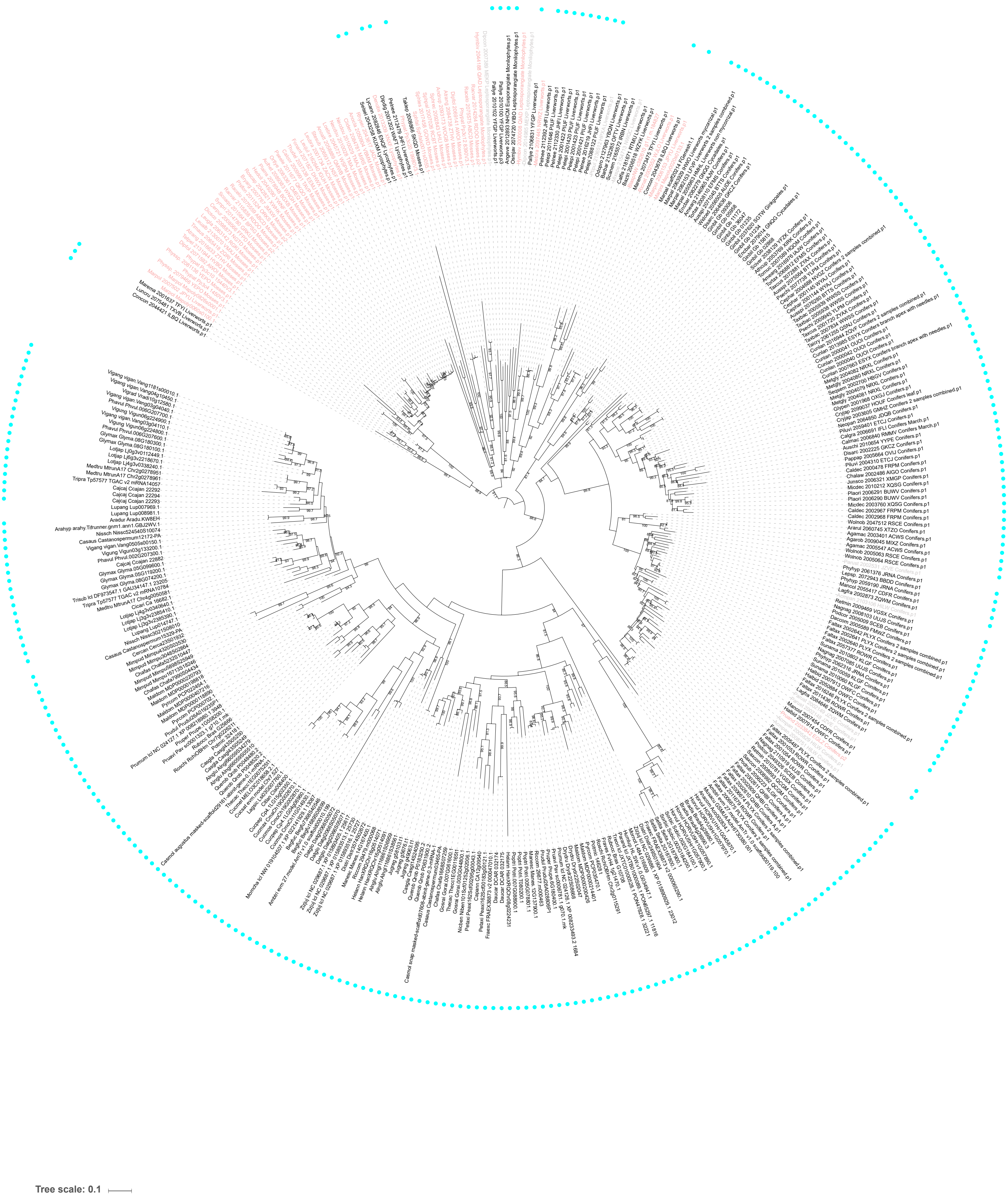

Supplementary Figure 13: Maximum likelihood tree of *CCaMK*.

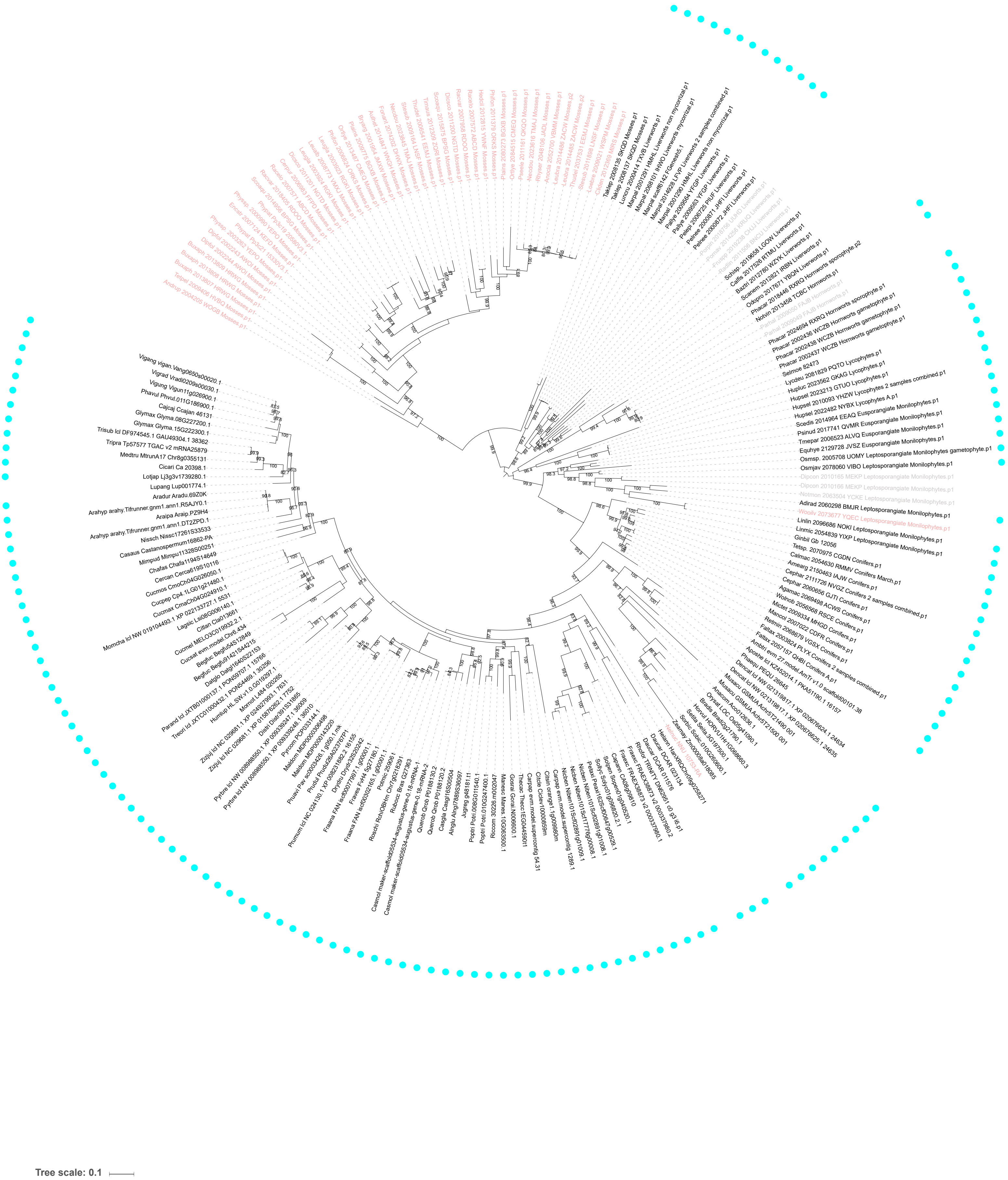

Supplementary Figure 14: Maximum likelihood tree of *CYCLOPS*.

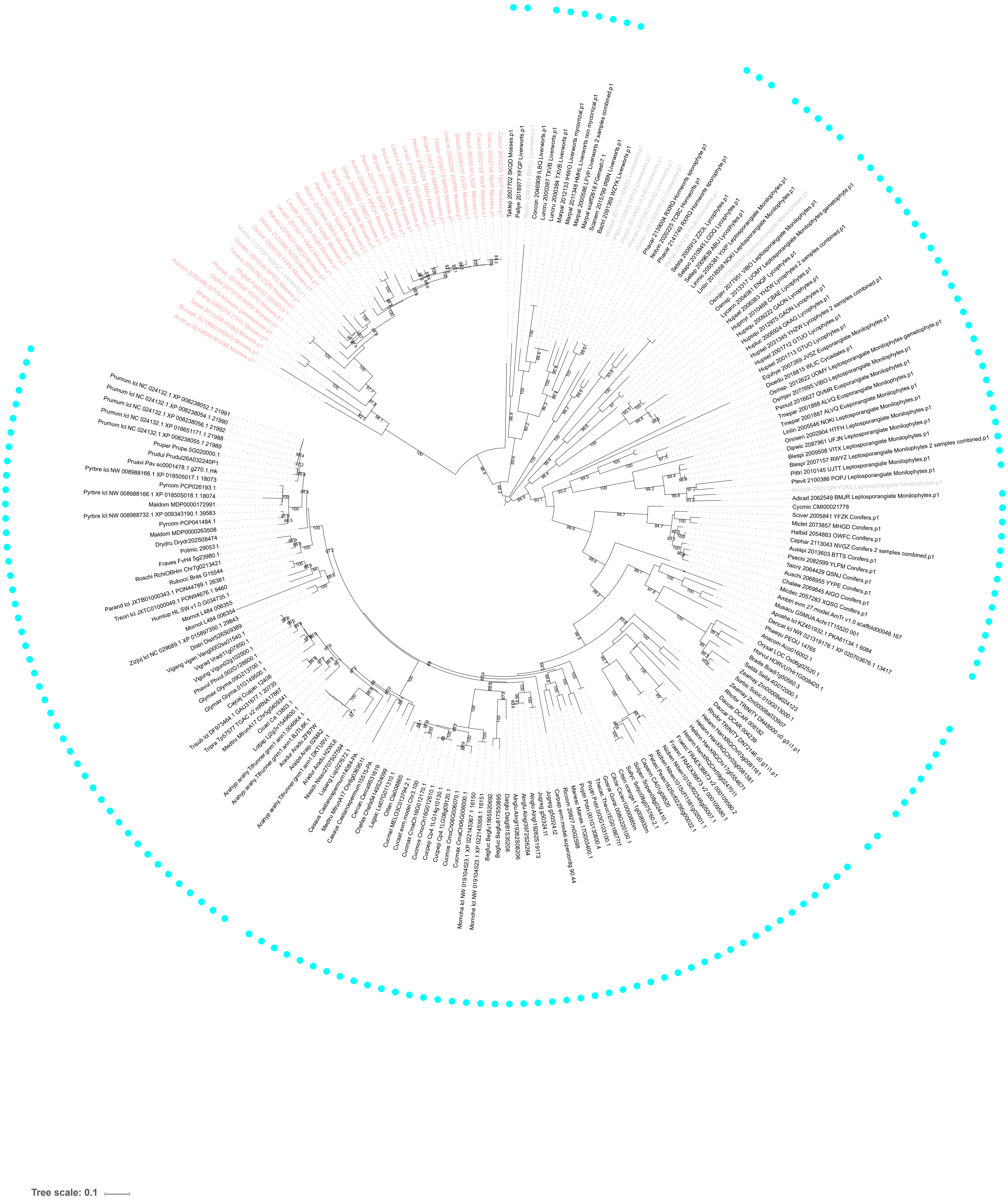

Supplementary Figure 15: Maximum likelihood tree of *SymRK*.

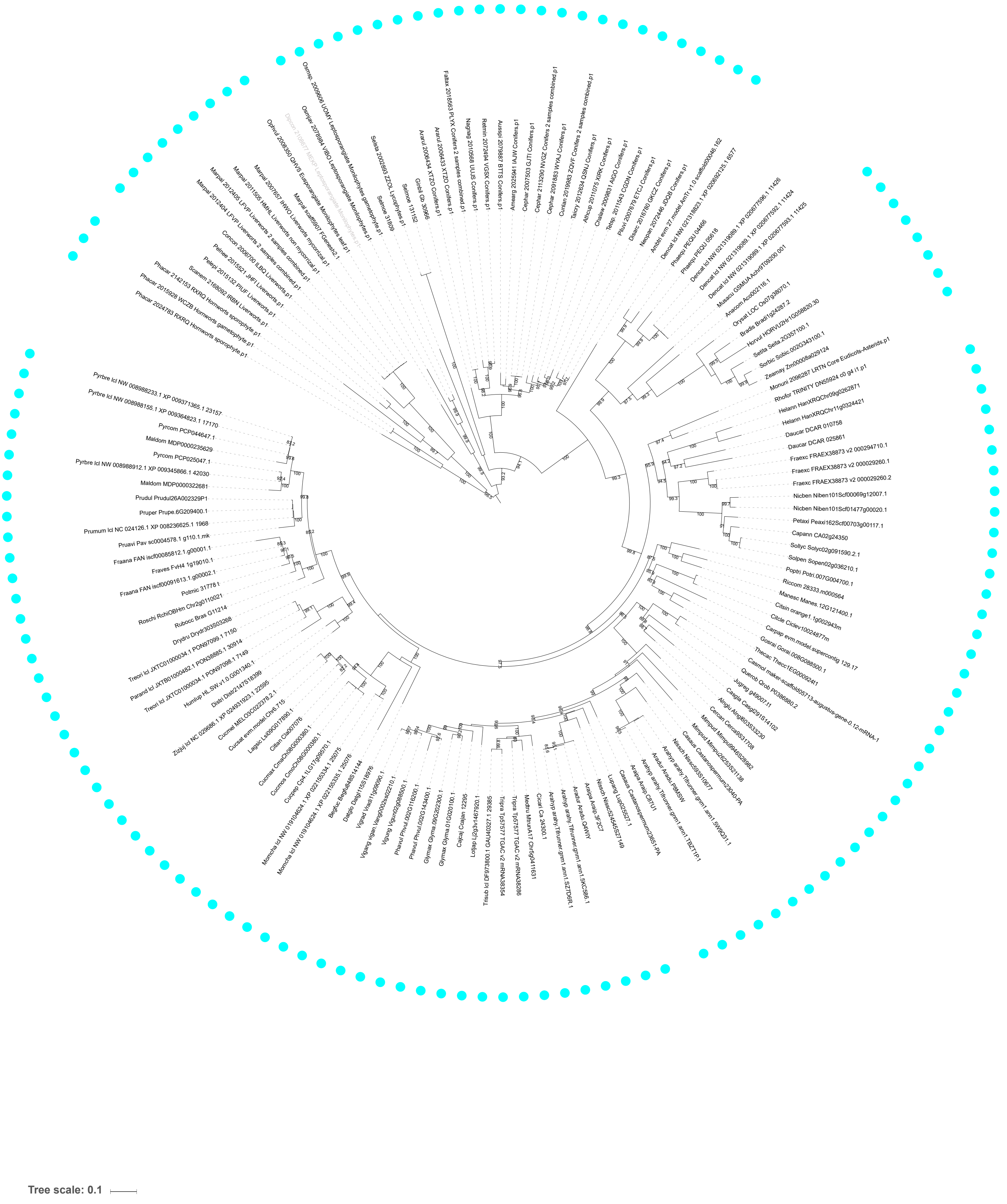

Supplementary Figure 16: Maximum likelihood tree of *HYP*.

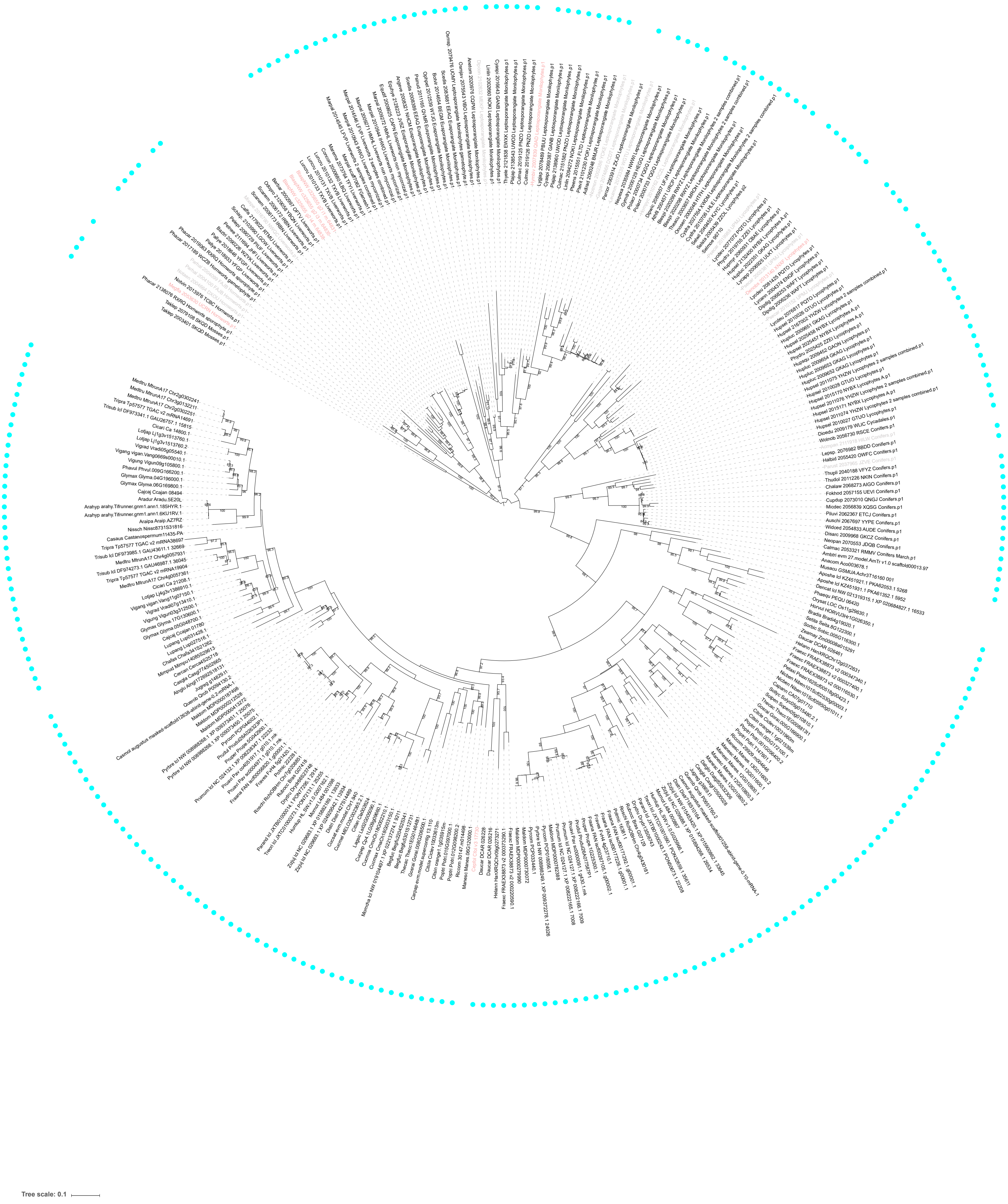

Supplementary Figure 17: Maximum likelihood tree of *HYP3*.

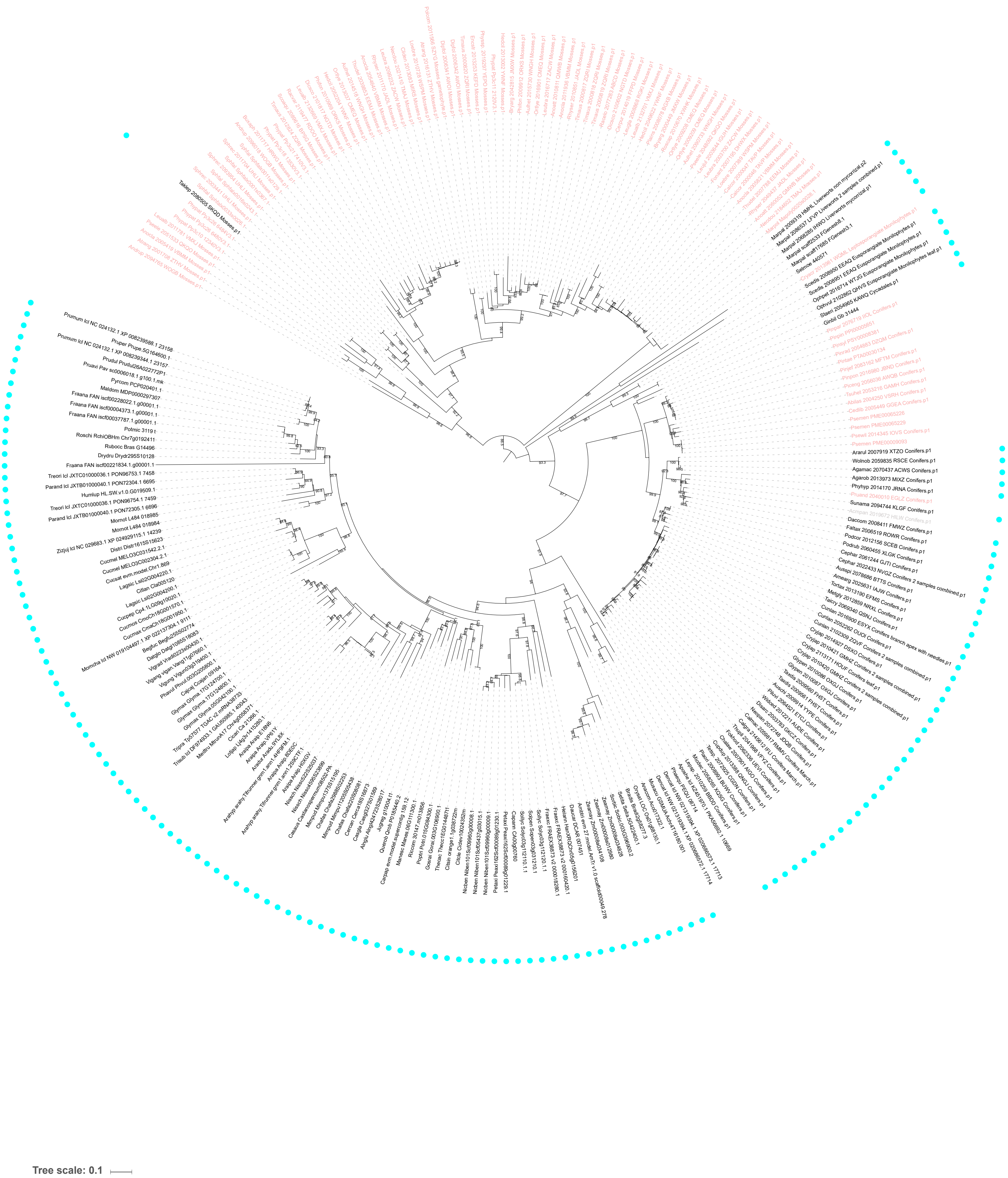

Supplementary Figure 18: Maximum likelihood tree of *CCD*.

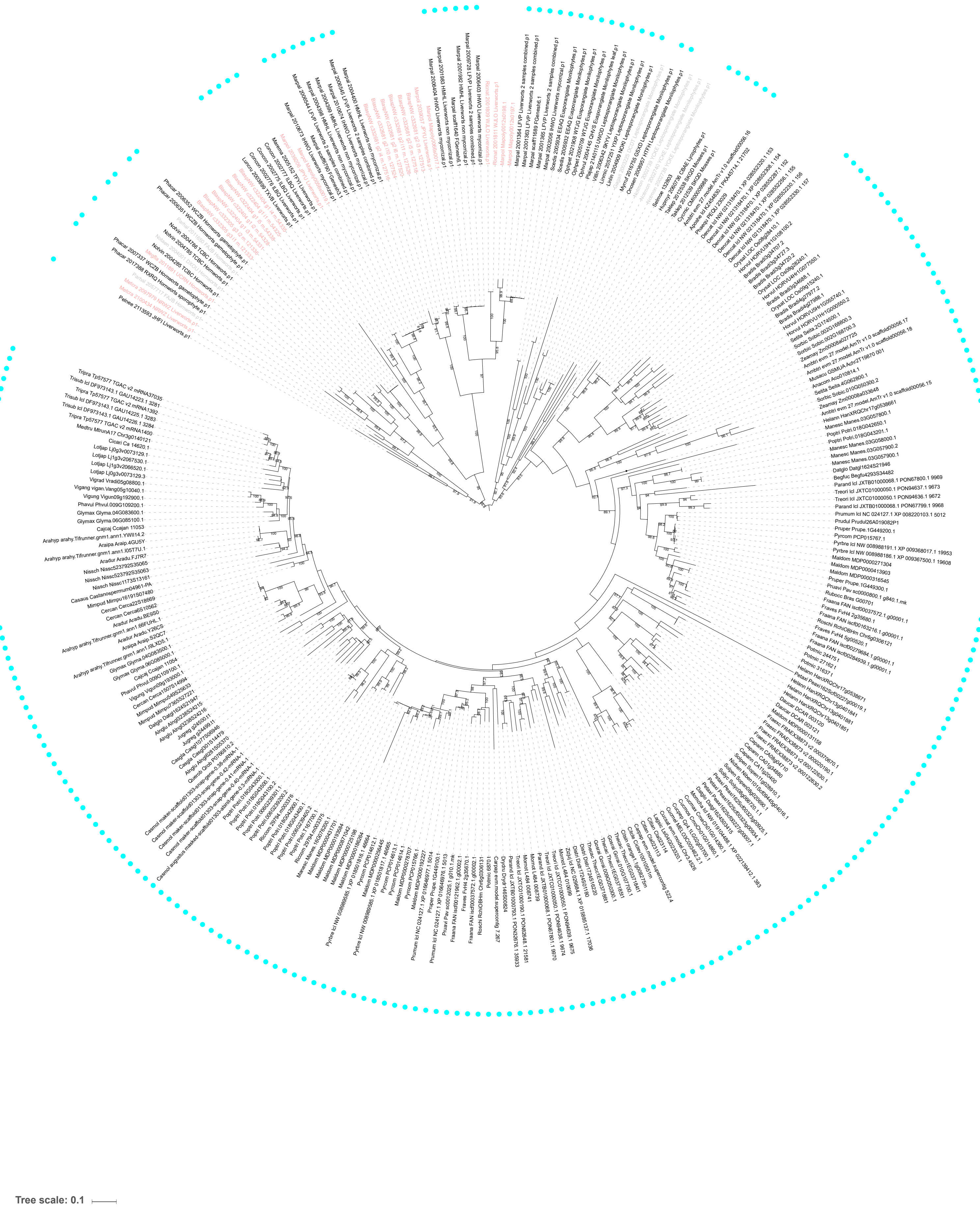

Supplementary Figure 19: Maximum likelihood tree of *RFCa/RFCb*.

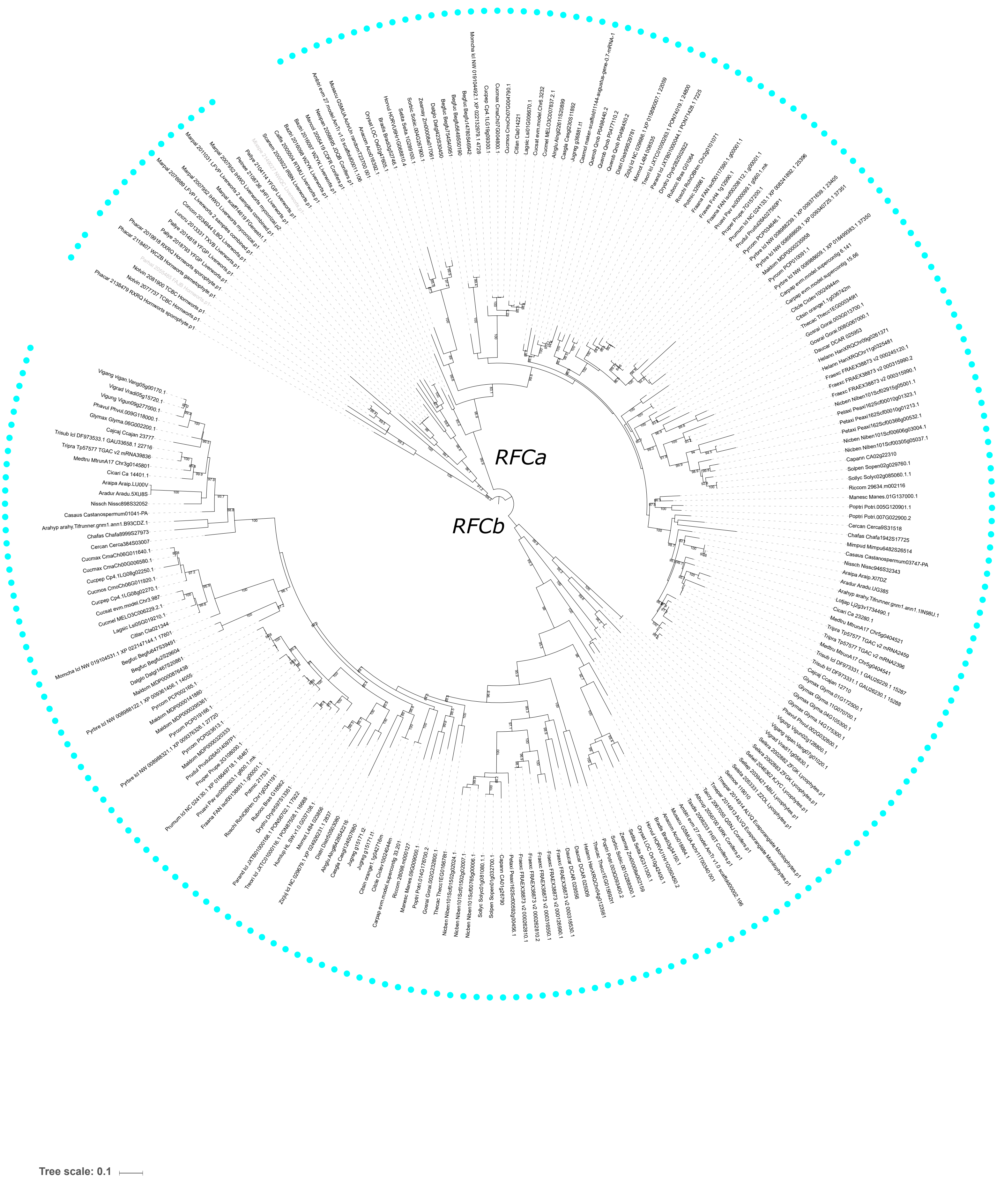

Supplementary Figure 20: Maximum likelihood tree of *KinG1/KinG2*.

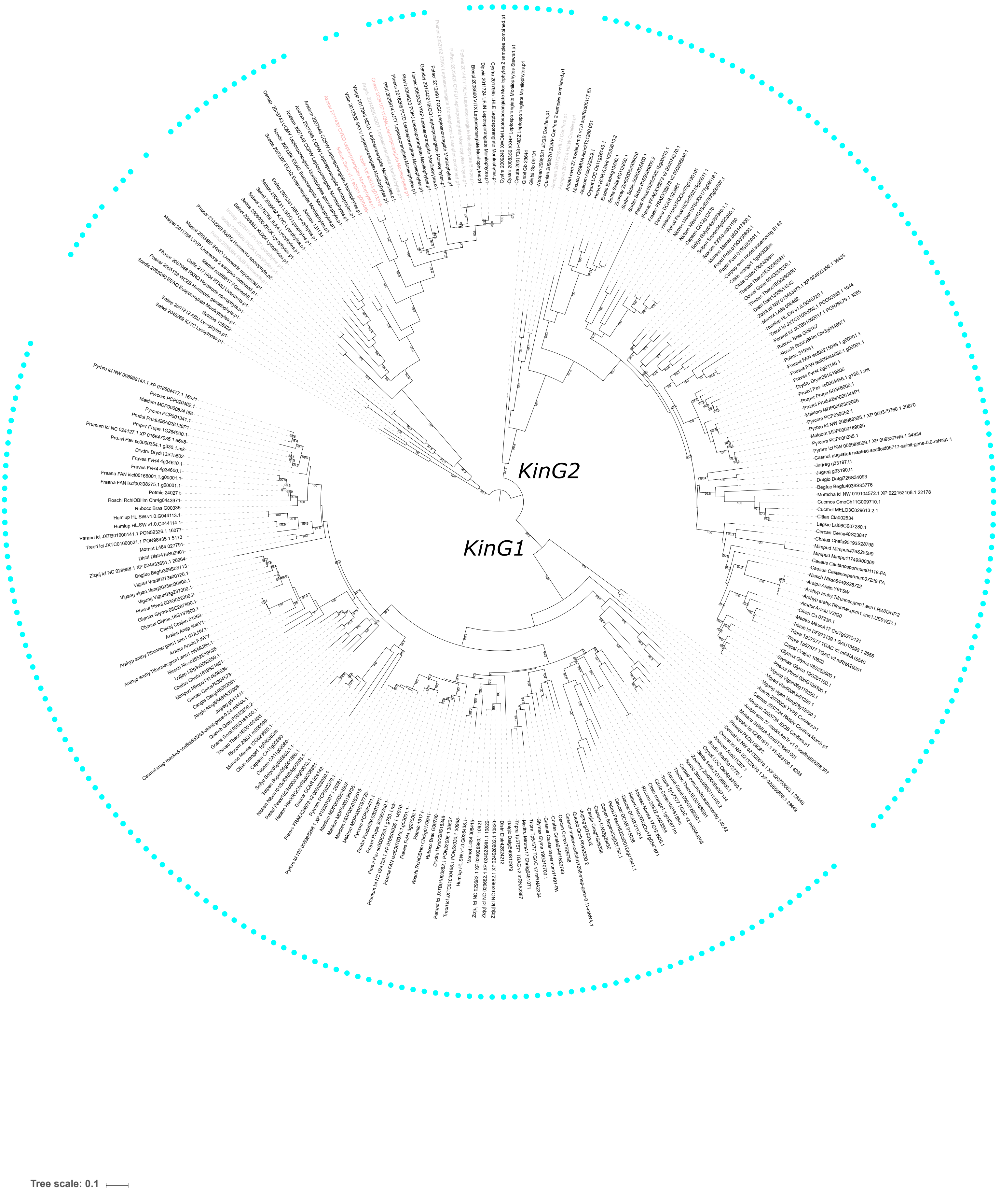

Supplementary Figure 21: Maximum likelihood tree of *VAPYRIN*/*VAPYRIN-like*.

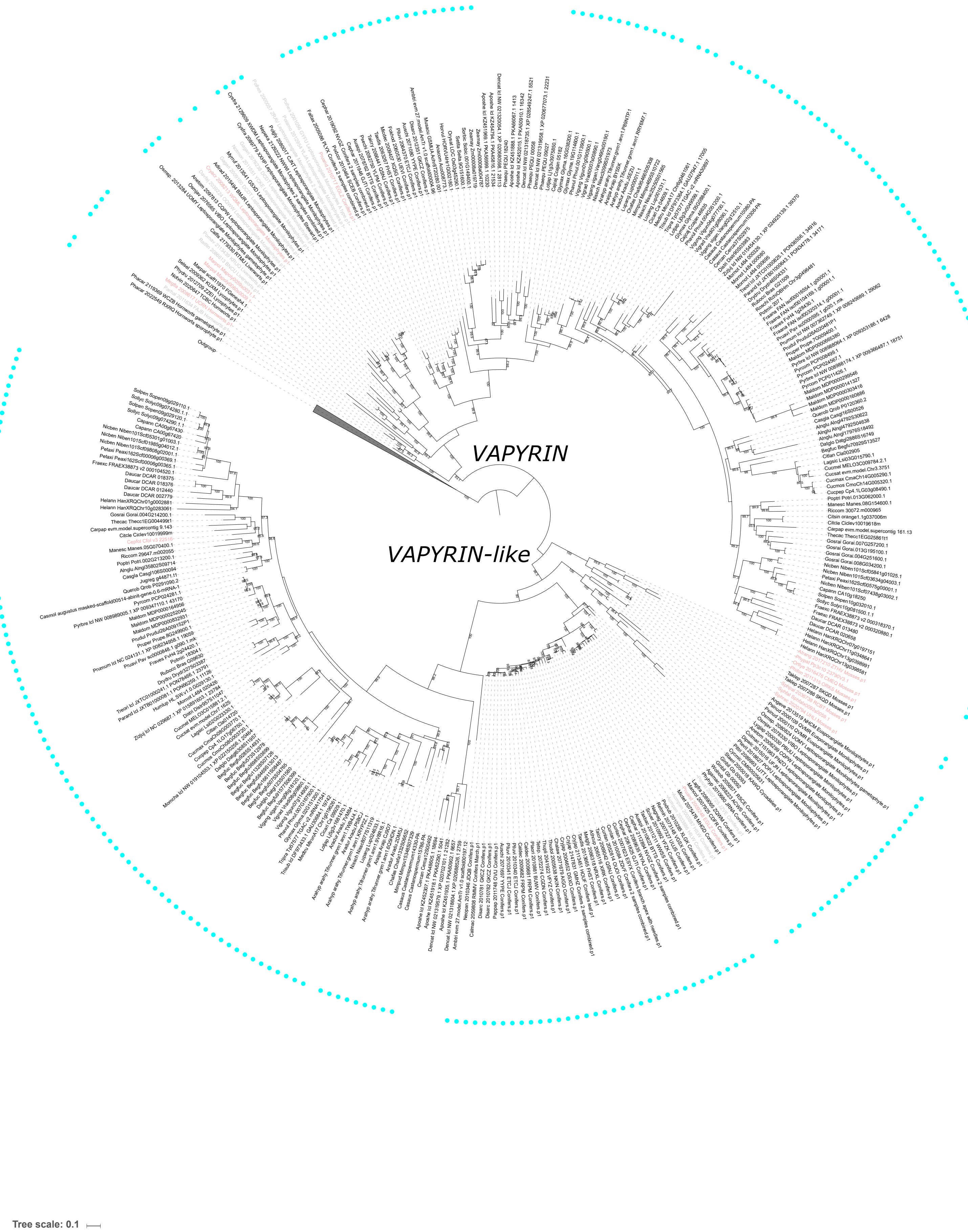

Supplementary Figure 22: Maximum likelihood tree of *LIN/LIN-like*.

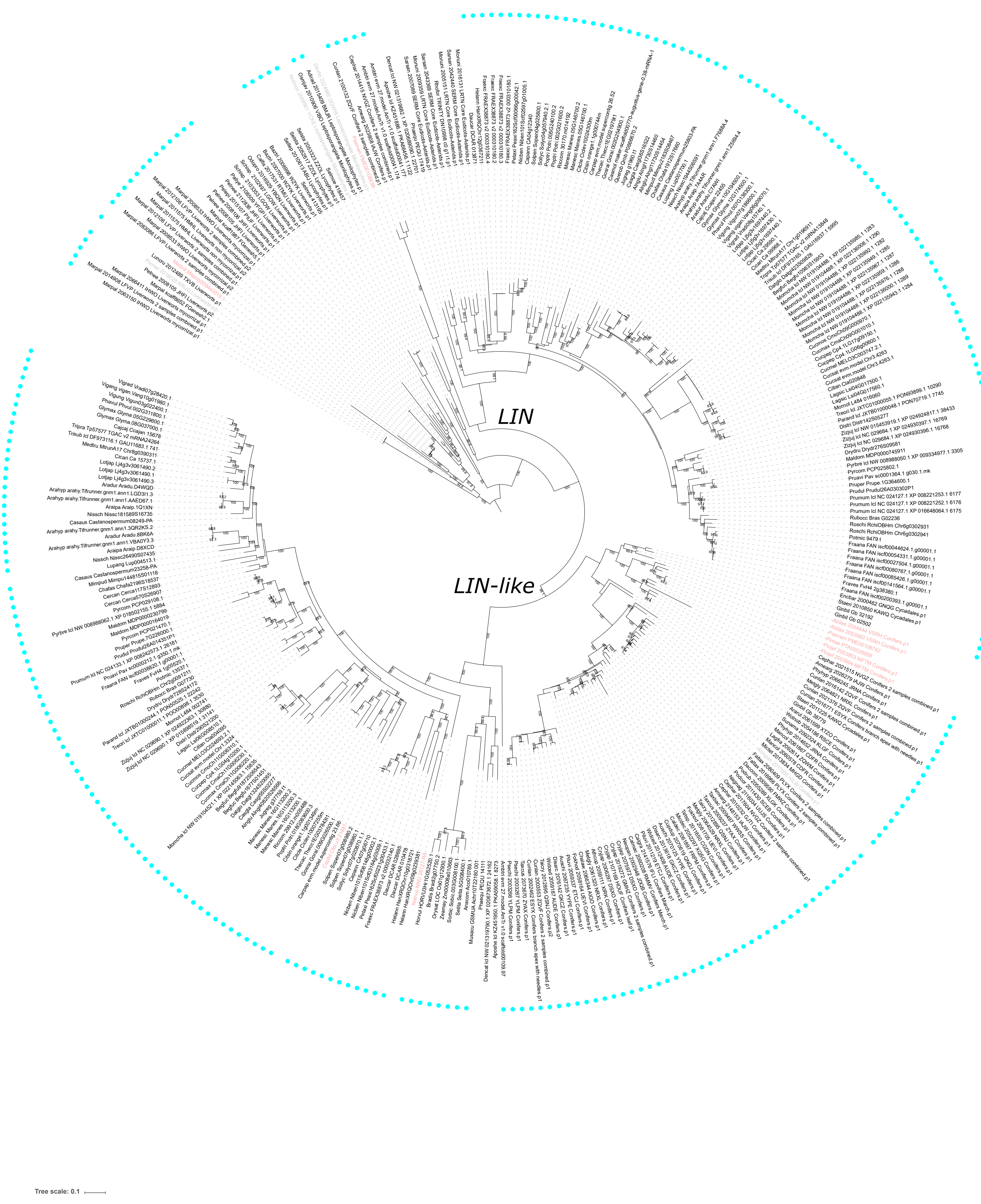

Supplementary Figure 23: Maximum likelihood tree of *CASTOR*.

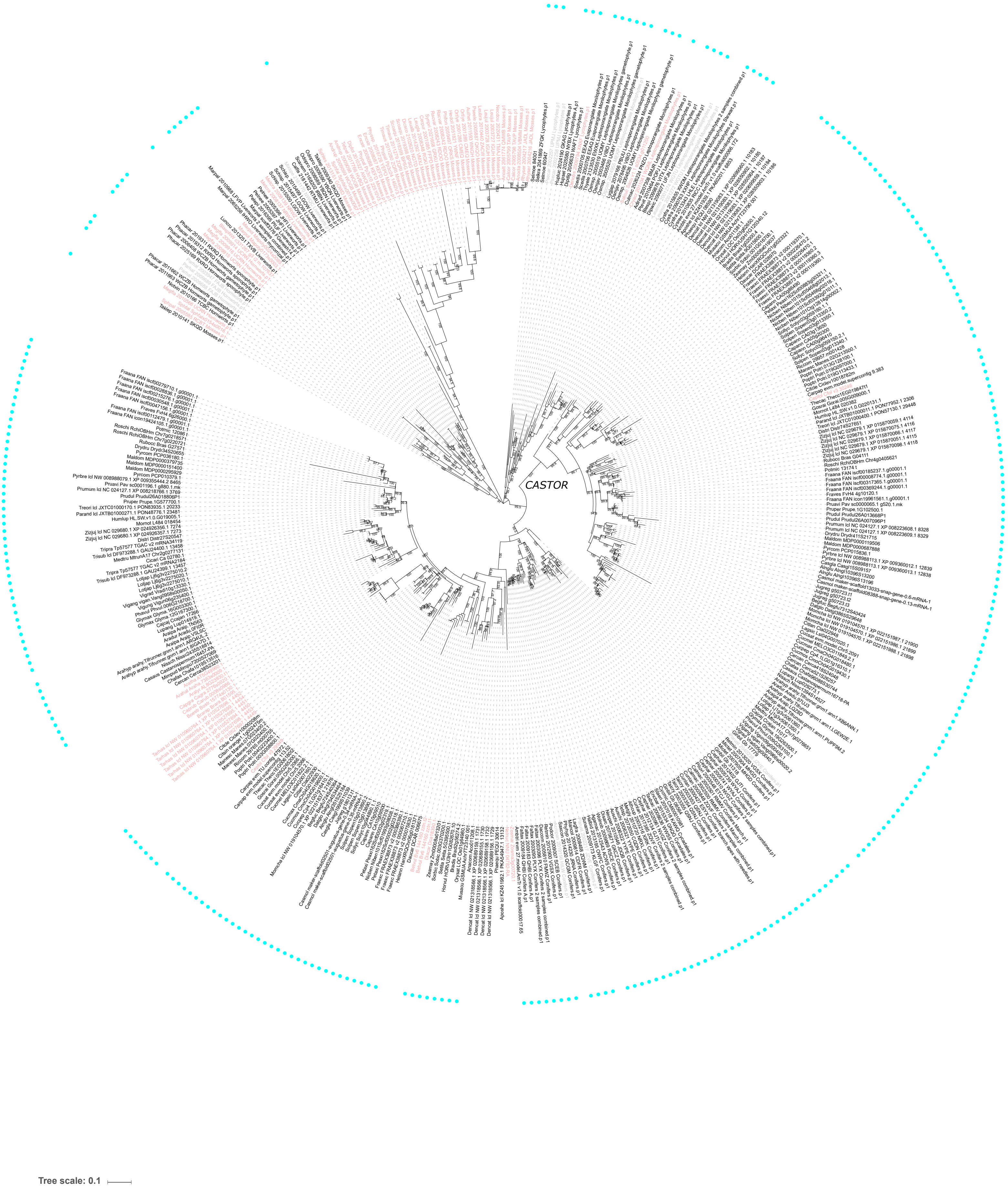

Supplementary Figure 24: Maximum likelihood tree of *EXO70*.

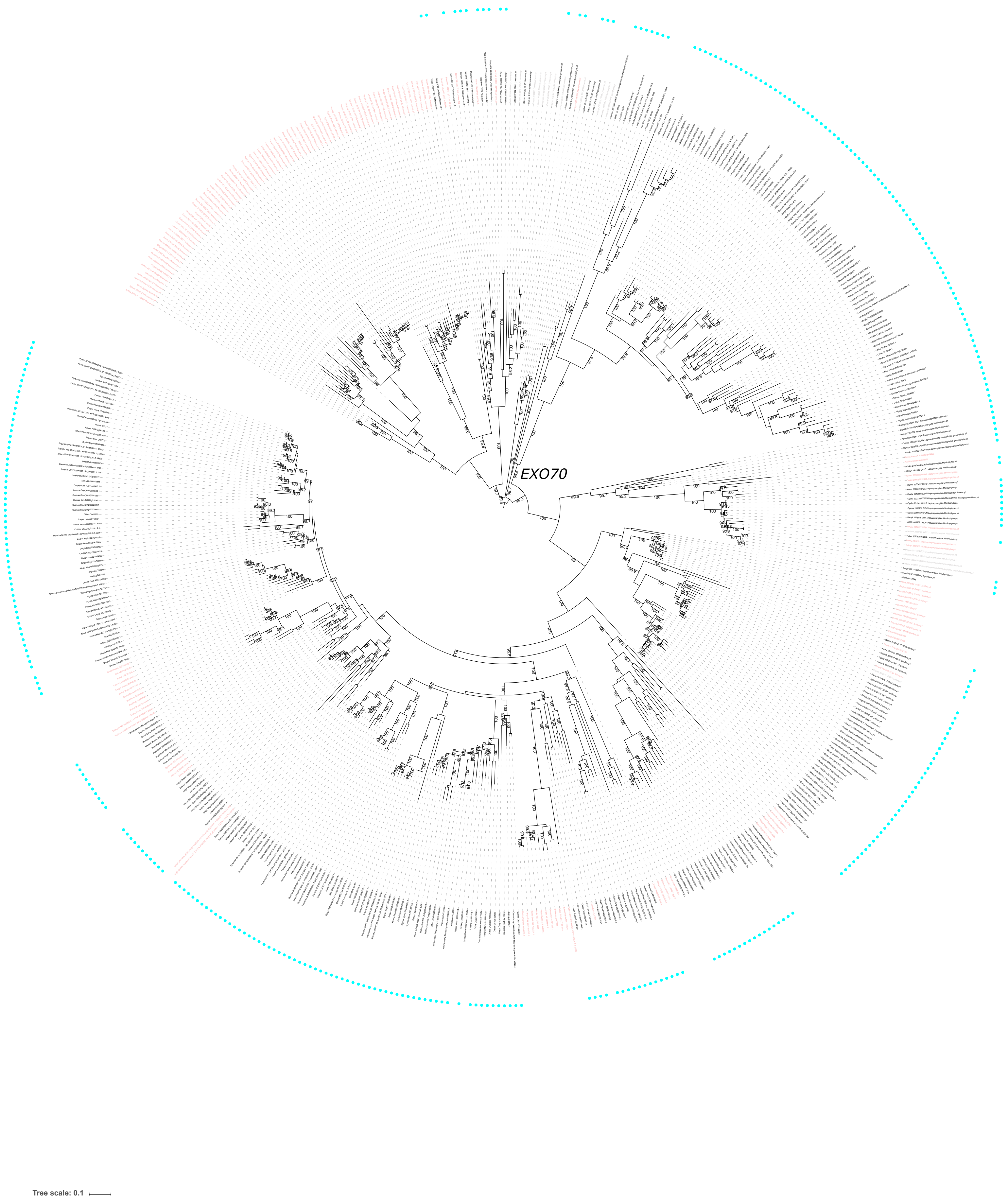

Supplementary Figure 25: Maximum likelihood tree of *FatM*.

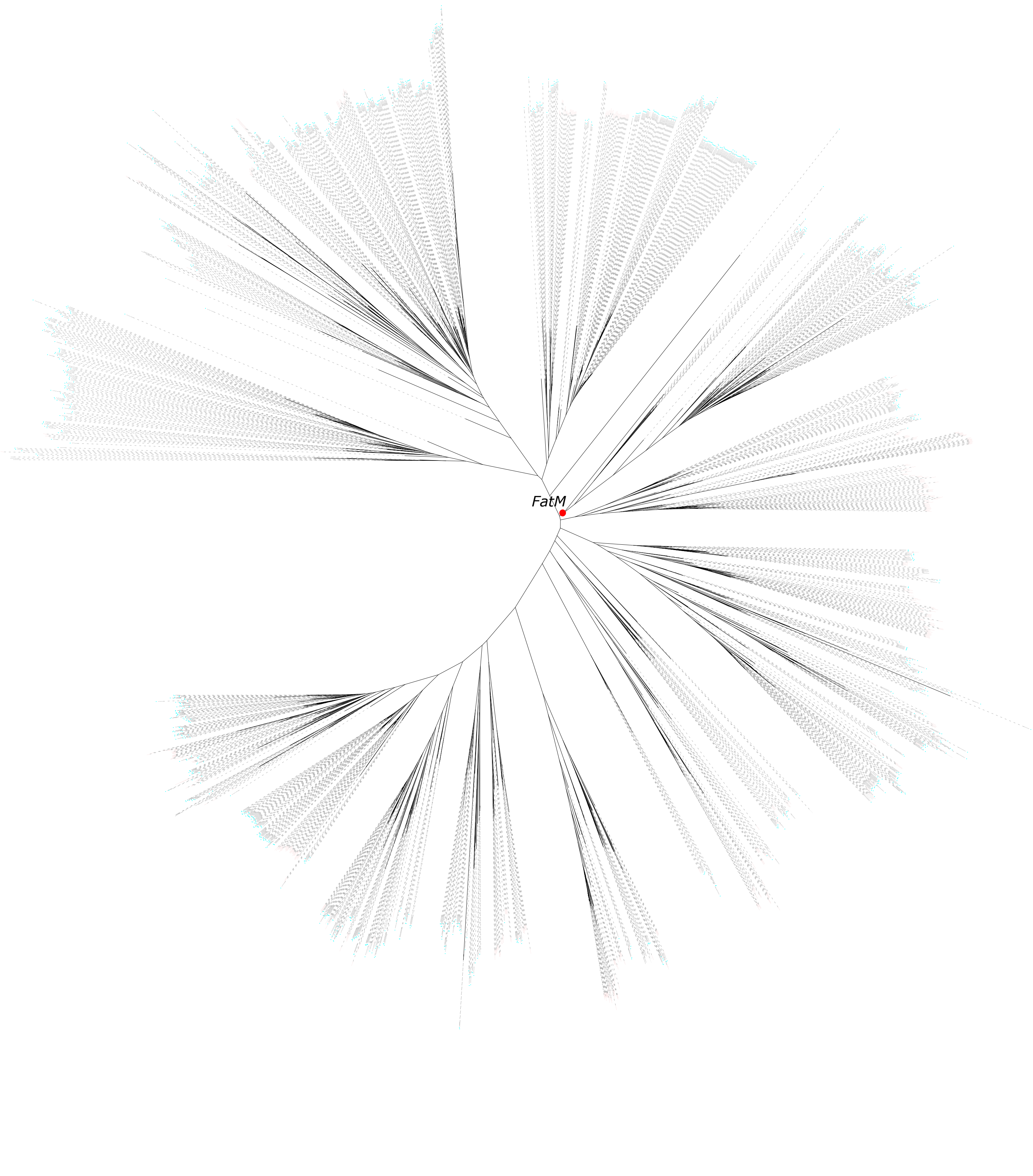

Supplementary Figure 26: Maximum likelihood tree of *NFP*.

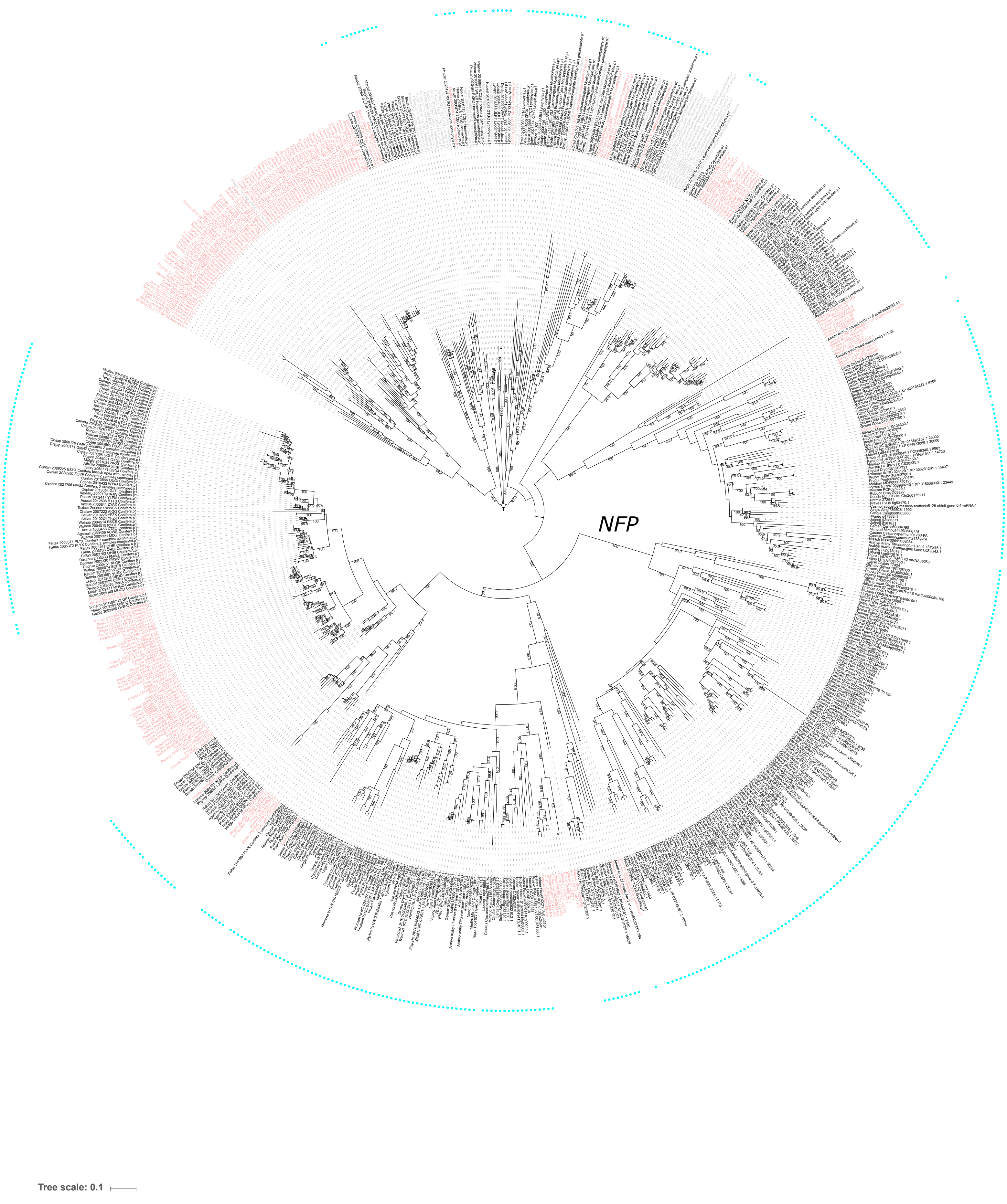

Supplementary Figure 27: Maximum likelihood tree of *SEC*.

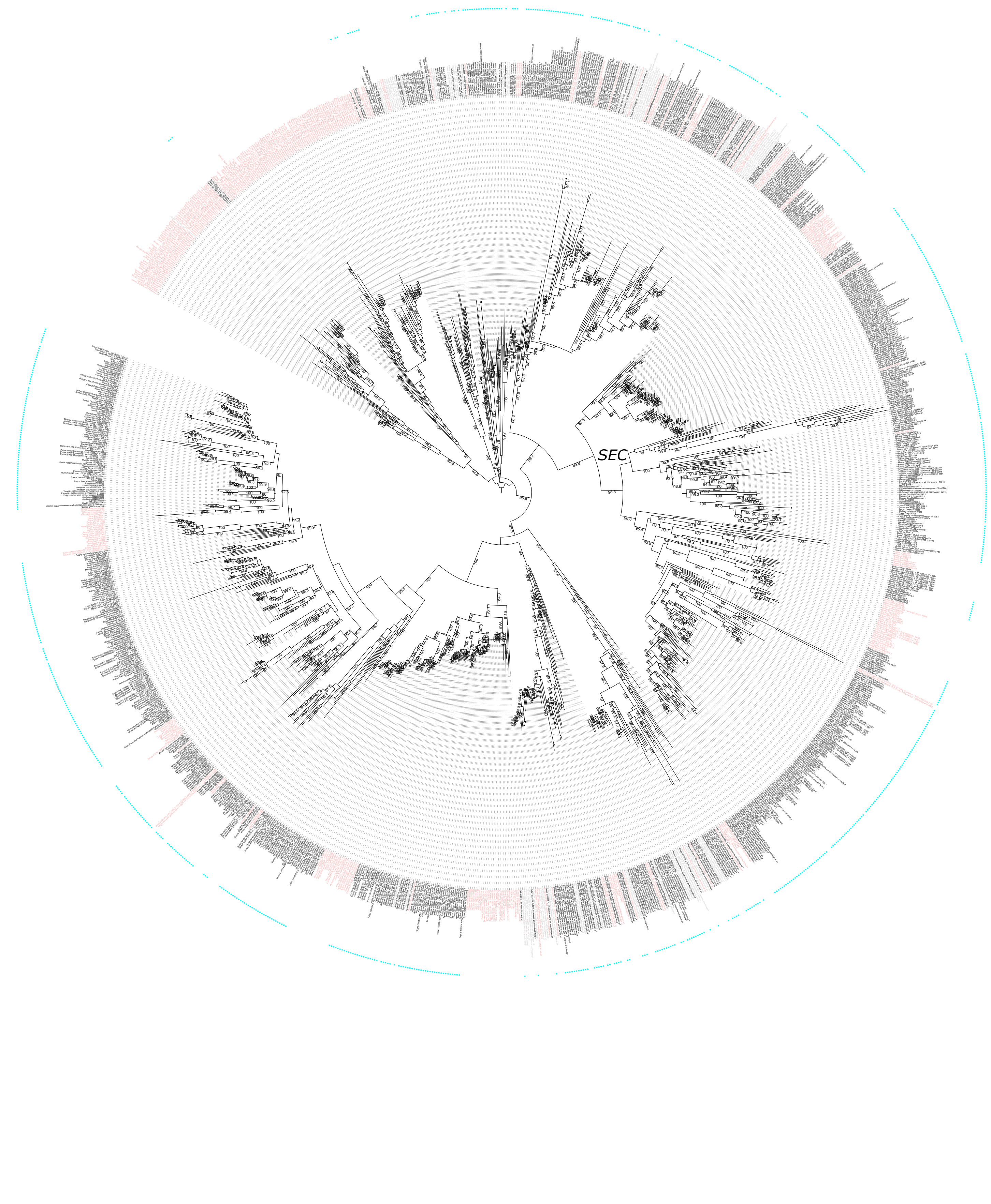

Supplementary Figure 28: Maximum likelihood tree of *SYN*.

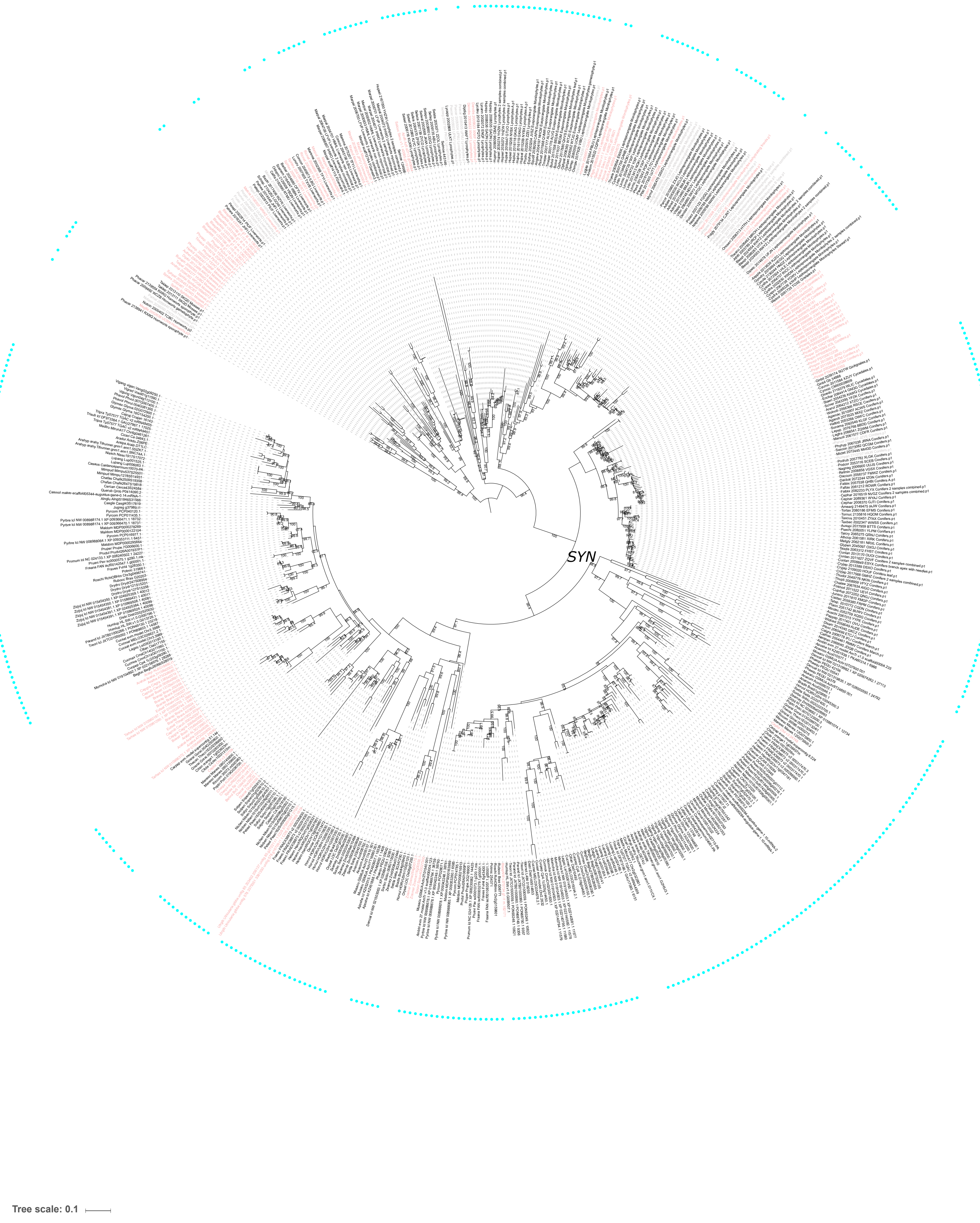

Supplementary Figure 29: Maximum likelihood tree of *CYTB561*.

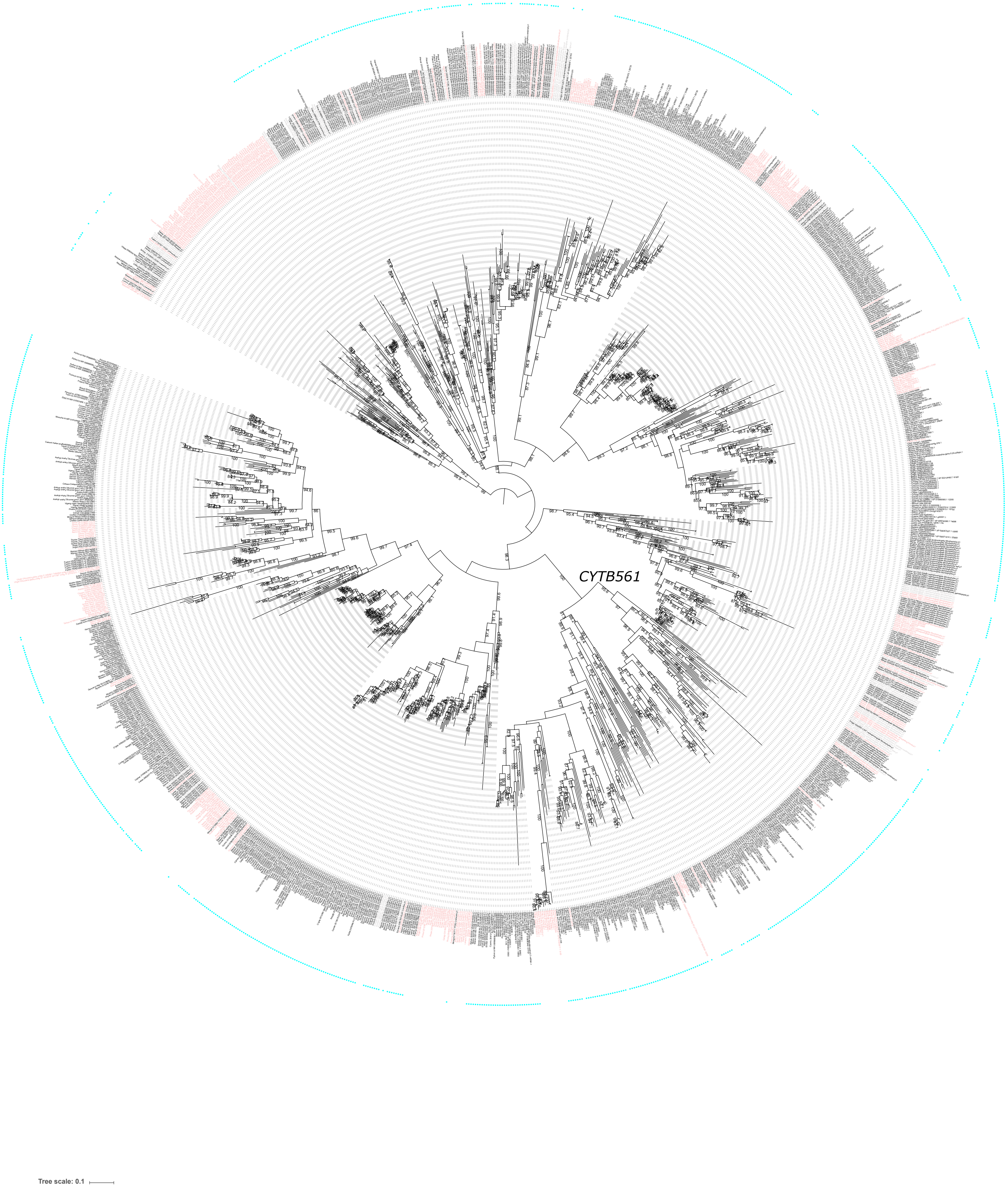

Supplementary Figure 30: Maximum likelihood tree of *GRAS*.

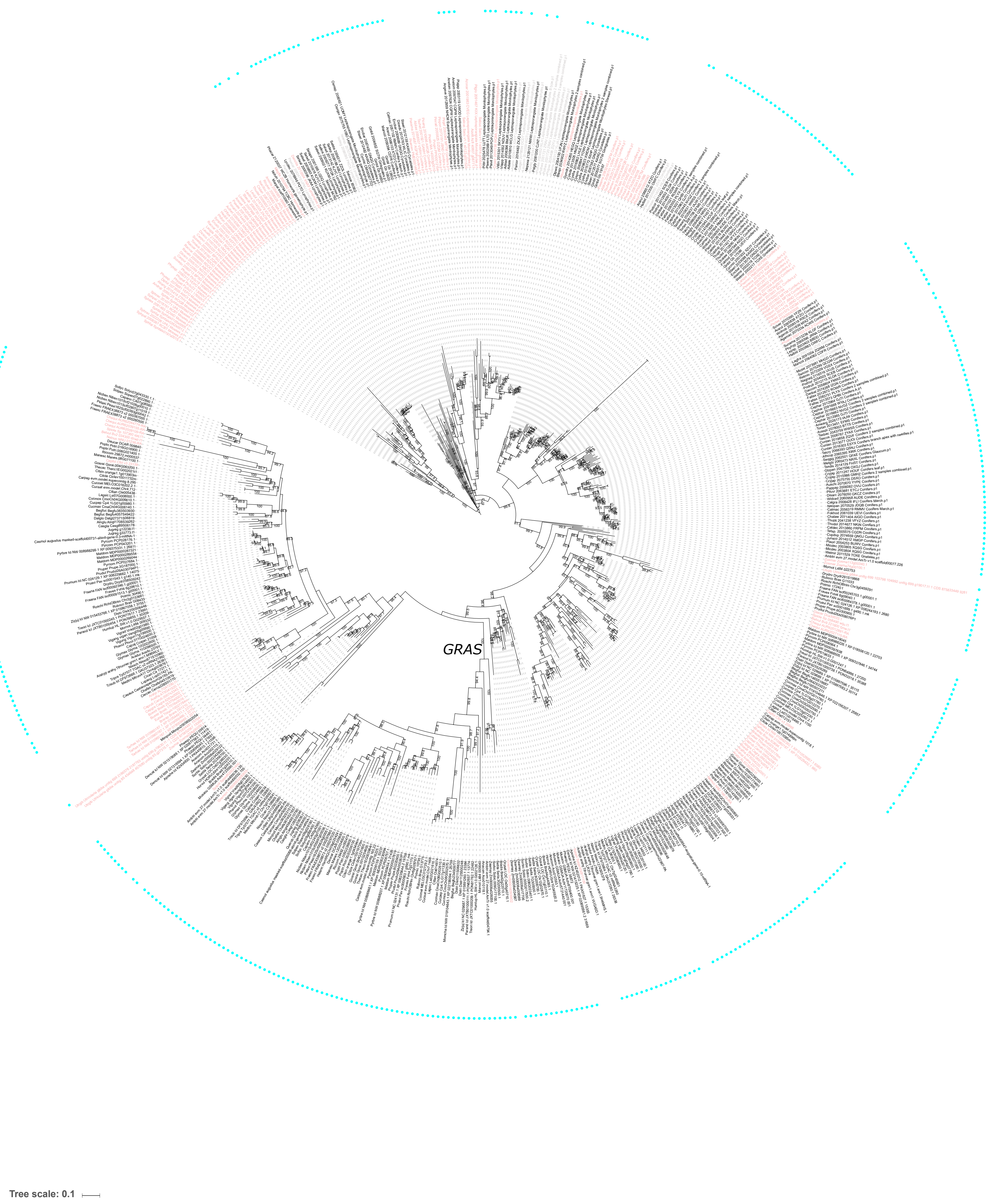

Supplementary Figure 31: Maximum likelihood tree of *HEP*.

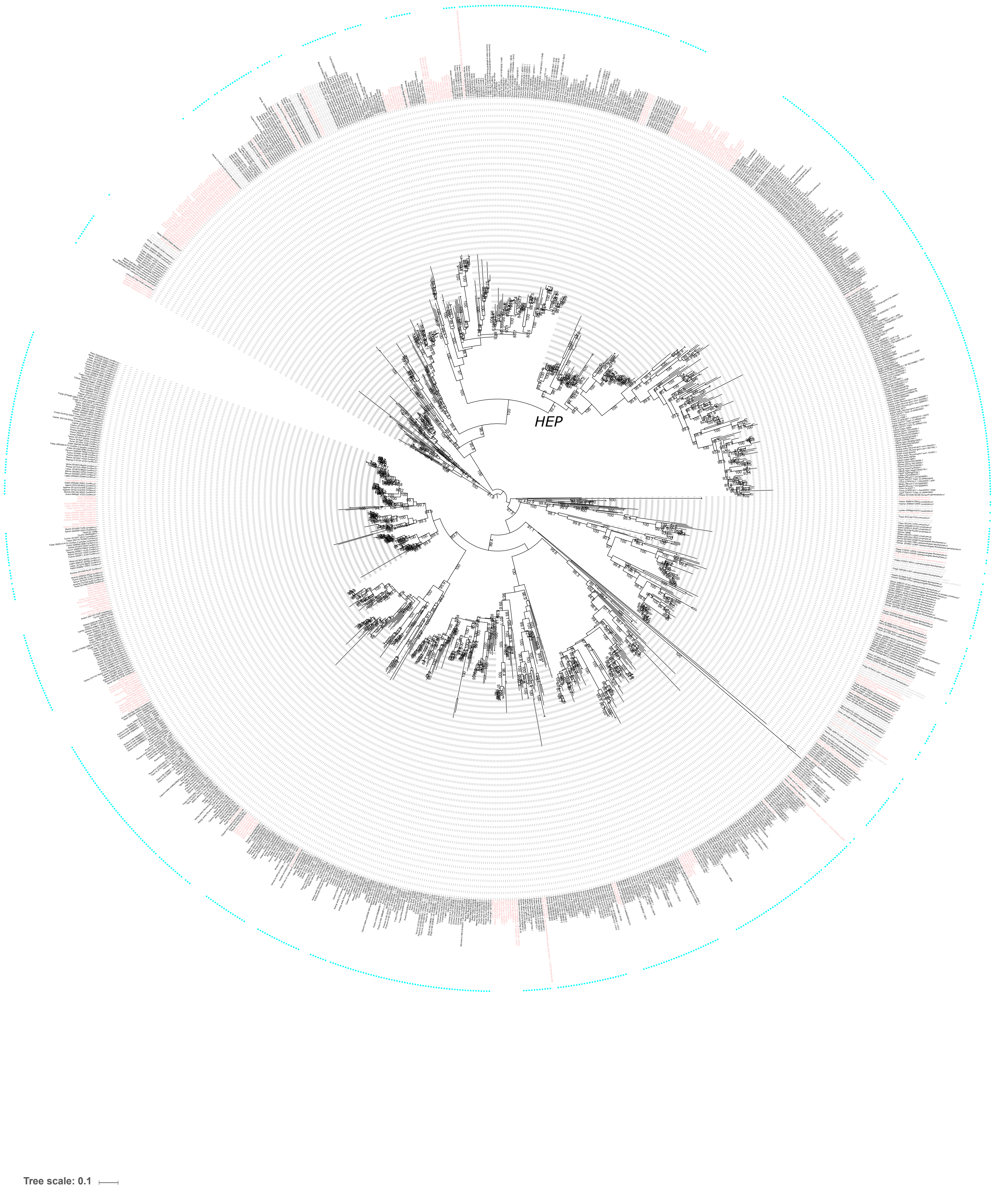

Supplementary Figure 32: Maximum likelihood tree of *TAU*.

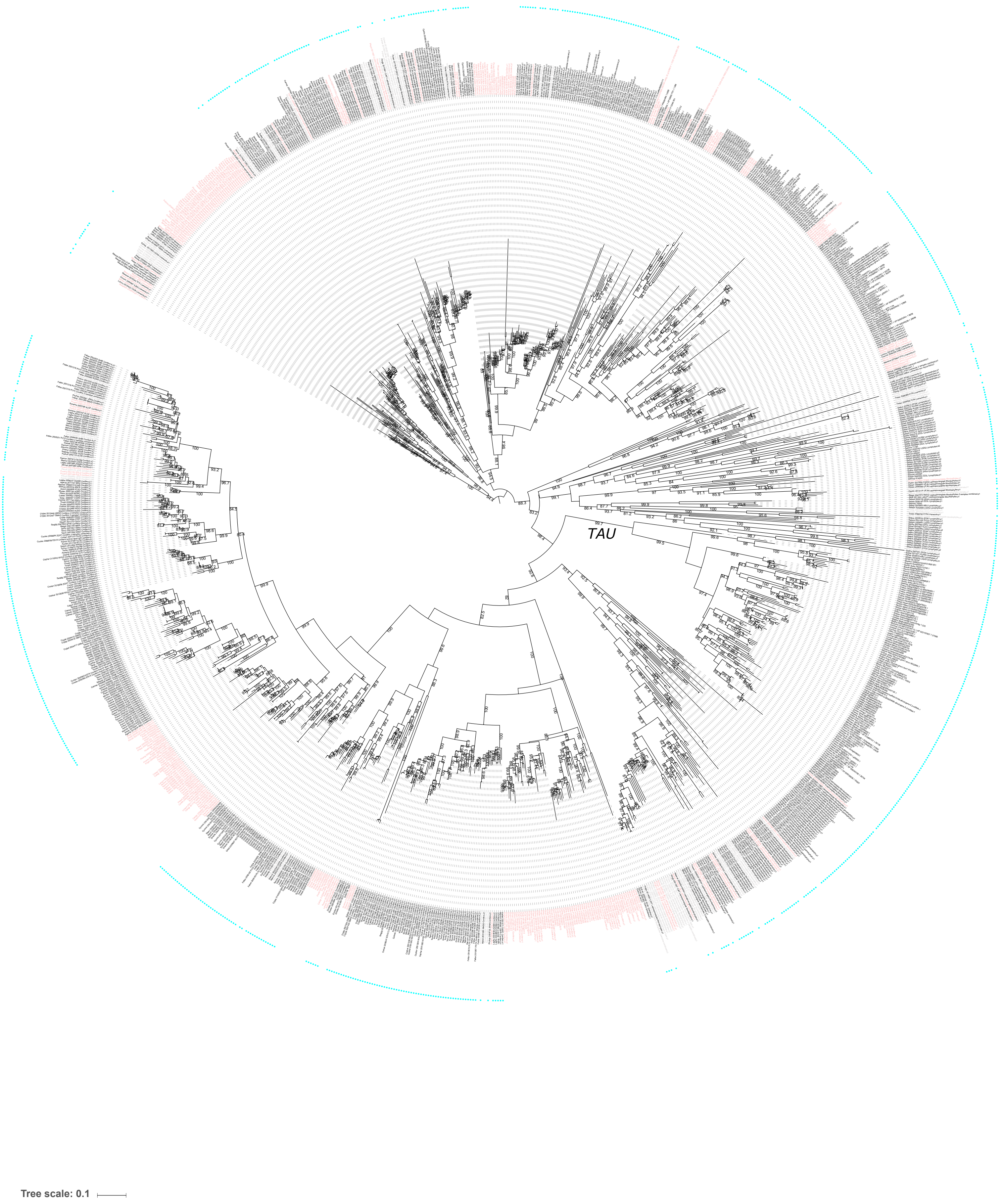

Supplementary Figure 33: Alignment of Nelumbo nucifera CCaMK from 3 assemblies with reference sequences showing an indel in the kinase domain and an amino acid switch on the essential for phosphorylation Serine 343

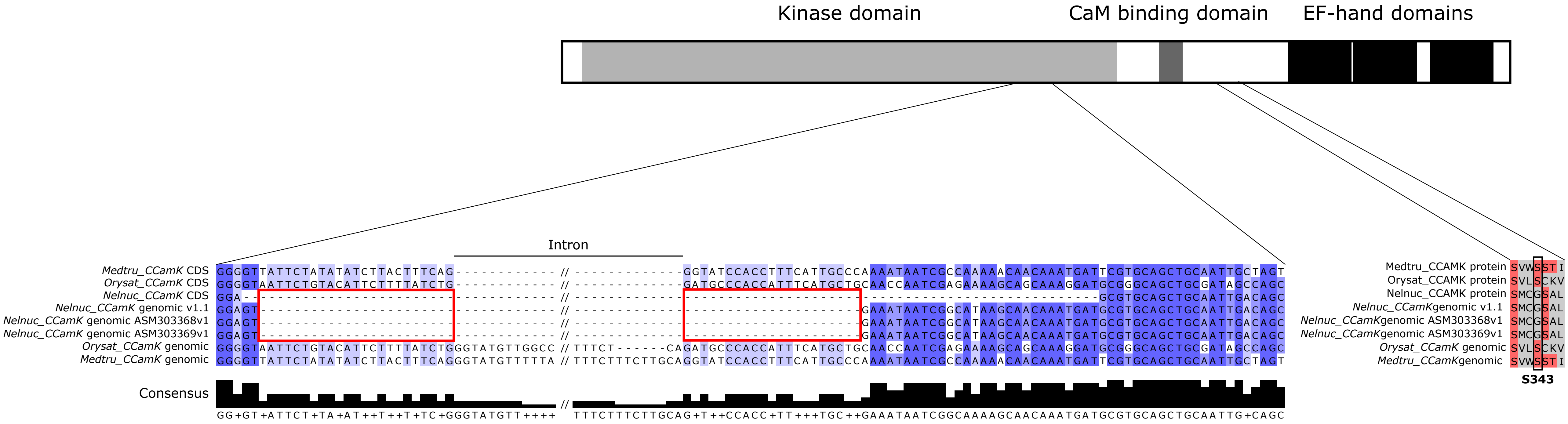

Supplementary Figure 34: Sites under positive selection in *CYCLOPS* in non-mycorrhizal mosses

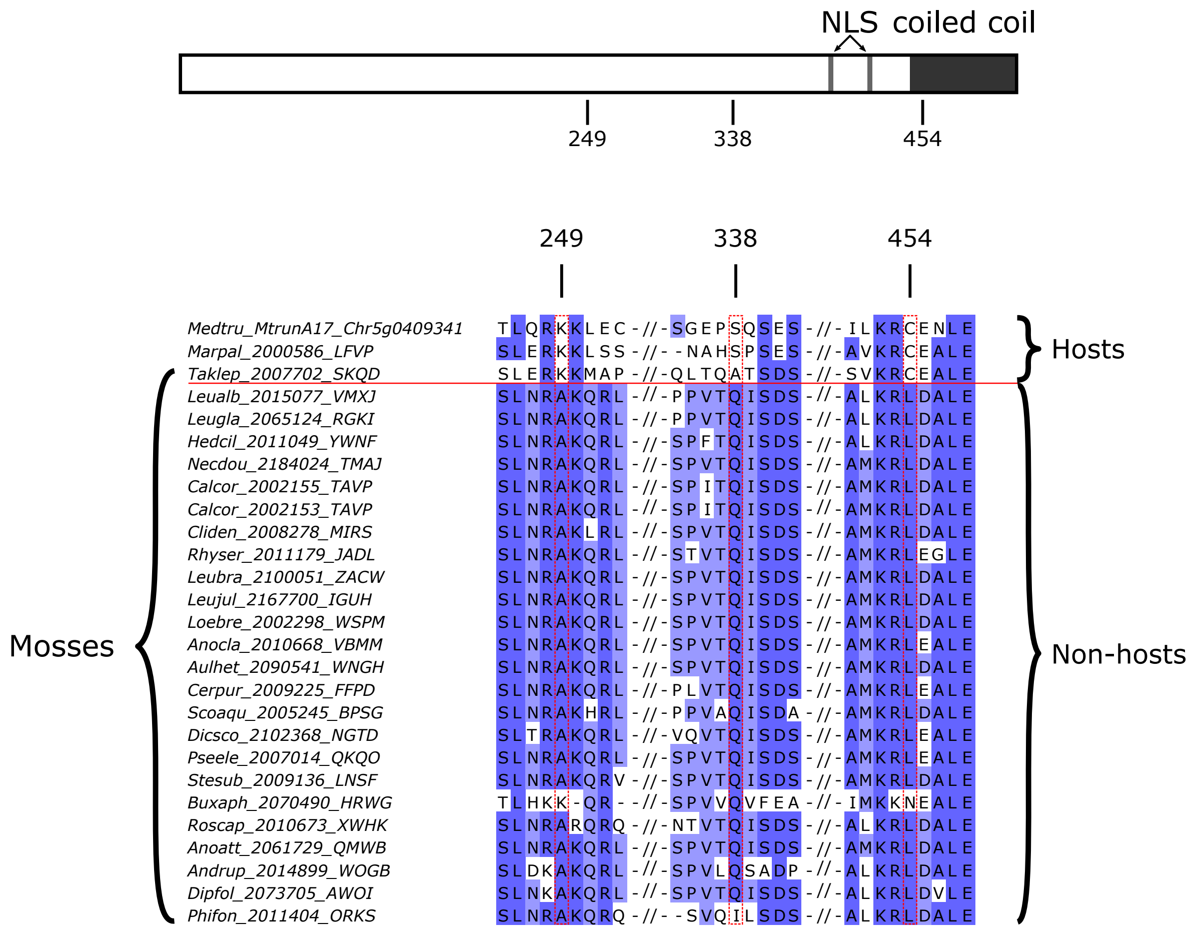

Supplementary Figure 35: Sites under positive selection in *CCaMK* in non-mycorrhizal mosses

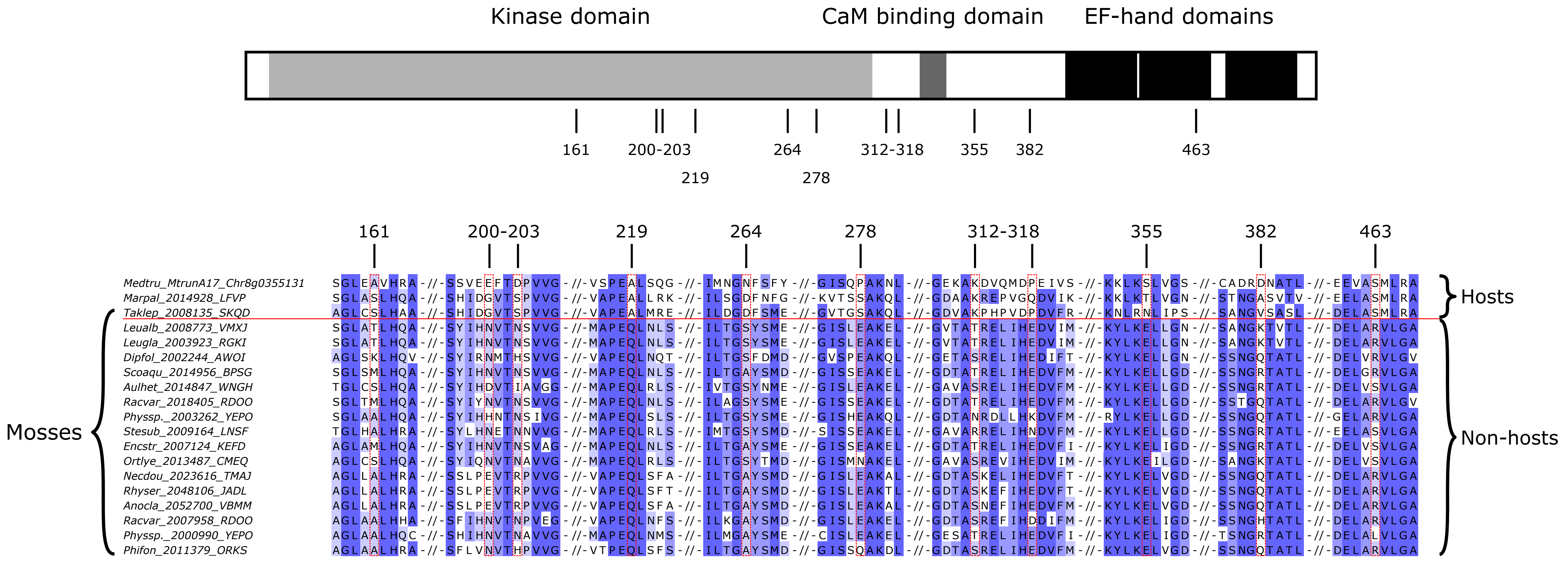

Supplementary Figure 36: Comparative alignment of 1kb from *CYCLOPS* among the three subspecies genomes and 35 accessions of *Marchantia polymorpha*.

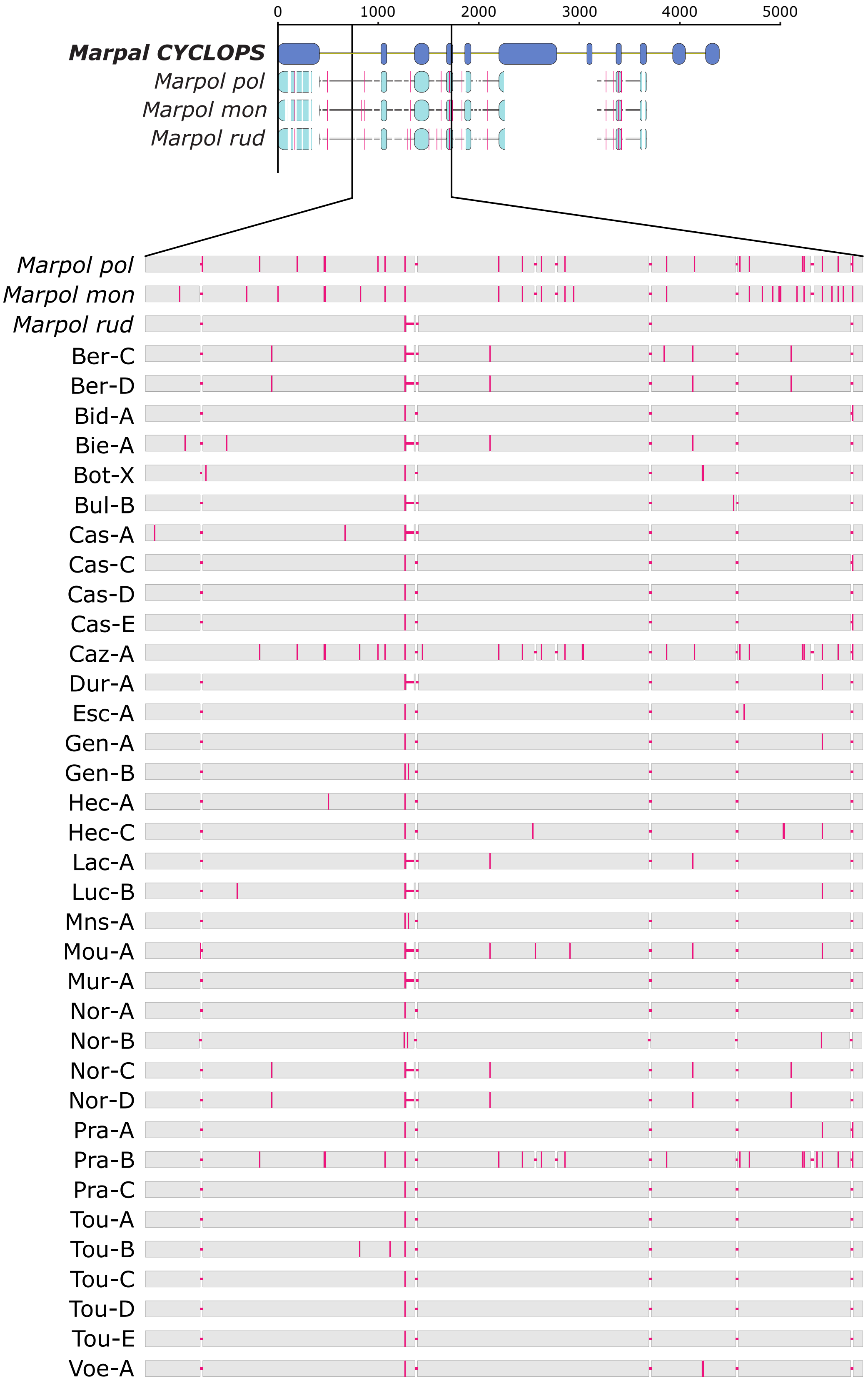

Supplementary figure 37: Presence/absence pattern of 11 genes in *Ericaceae* compared to species with different symbiotic abilities

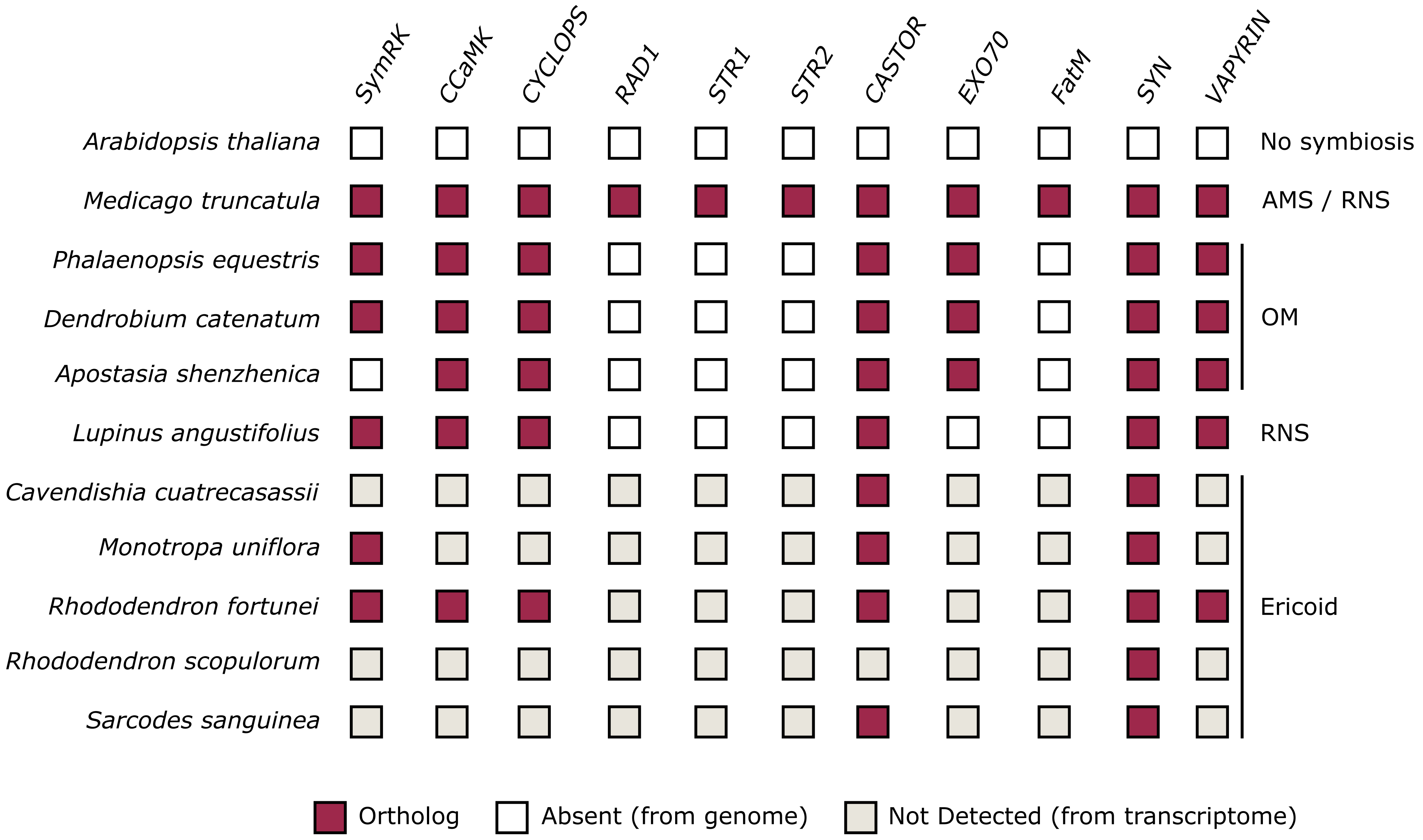

**Supplementary Table 1:** List of species investigated in this study. For each species symbiotic statues are indicated by 0 (not established), 1 (established) or ND (not determined) according to litterature.
AMS: Arbuscular Mycorrhizal Symbiosis; RNS: Root Nodule Symbiosis; OM: Orchid Mycorrhiza; EcM: Ectomycorrhizal.

| Species name | Group | Order | Family | abbreviation | Source | AMS | RNS | OM | Ericoid-like | Cyanobacteria | EcM | Endosymbiosis | Extracellular symbiosis | Code 1KP | Source AMS | Source extracellular symbiosis | Comments |
|---|---|---|---|---|---|---|---|---|---|---|---|---|---|---|---|---|---|
| *Abies lasiocarpa* | Gymnosperms | Pinales | Pinaceae | Abilas | 1KP; 10.1186/2047-217X-3-17 | 0 | 0 | 0 | 0 | 0 | 1 | 0 | 1 | VSRH | . | 10.1139/b97-896 | . |
| *Acmopyle pancheri* | Gymnosperms | Pinales | Podocarpaceae | Acmpan | 1KP; 10.1186/2047-217X-3-17 | ND | 0 | 0 | 0 | 0 | 0 | ND | 0 | HILW | . | . | . |
| *Adiantum aleuticum* | Leptosporangiate | Polypodiales | Pteridaceae | Adiale | 1KP; 10.1186/2047-217X-3-17 | 1 | 0 | 0 | 0 | 0 | 0 | 1 | 0 | WCLG | . | . | . |
| *Adiantum raddianum* | Leptosporangiate | Polypodiales | Pteridaceae | Adirad | 1KP; 10.1186/2047-217X-3-17 | 1 | 0 | 0 | 0 | 0 | 0 | 1 | 0 | BMJR | 10.1007/s00572-004-0306-5 | . | . |
| *Agathis macrophylla* | Gymnosperms | Pinales | Araucariaceae | Agamac | 1KP; 10.1186/2047-217X-3-17 | 1 | 0 | 0 | 0 | 0 | 0 | 1 | 0 | ACWS | 10.1016/j.funbio.2016.01.015 | . | . |
| *Agathis robusta* | Gymnosperms | Pinales | Araucariaceae | Agarob | 1KP; 10.1186/2047-217X-3-17 | 1 | 0 | 0 | 0 | 0 | 0 | 1 | 0 | MIXZ | 10.1016/j.funbio.2016.01.015 | . | . |
| *Alnus glutinosa* | Angiosperms | Fagales | Betulaceae | Alnglu | 10.1126/science.aat1743 | 1 | 1 | 0 | 0 | 0 | 1 | 1 | 1 | NA | Fraga-Beddiar, A., & Tacon, F. L. (1990). | 10.1007/s11104-008-9877-9 | . |
| *Amborella trichopoda* | Angiosperms | Amborellales | Amborellaceae | Ambtri | 10.1126/science.1241089 | 1 | 0 | 0 | 0 | 0 | 0 | 1 | 0 | NA | . | . | . |
| *Amentotaxus argotaenia* | Gymnosperms | Pinales | Taxaceae | Amearg | 1KP; 10.1186/2047-217X-3-17 | 1 | 0 | 0 | 0 | 0 | 0 | 1 | 0 | IAJW | www.mycorrhizas.info/ | . | . |
| *Ananas comosus* | Angiosperms | Bromeliales | Bromeliaceae | Anacom | 10.1038/ng.3435 | 1 | 0 | 0 | 0 | 0 | 0 | 1 | 0 | NA | 10.1080/00103629009368382 | . | . |
| *Andreaea rupestris* | Mosses | Andreaeales | Andreaeaceae | Andrup | 1KP; 10.1186/2047-217X-3-17 | 0 | 0 | 0 | 0 | 0 | 0 | 0 | 0 | WOGB | 10.1098/rstb.2000.0617 | . | . |
| *Anemia tomentosa* | Leptosporangiate | Schizaeales | Anemiaceae | Anetom | 1KP; 10.1186/2047-217X-3-17 | 1 | 0 | 0 | 0 | 0 | 0 | 1 | 0 | CQPW | 10.1007/s00572-009-0231-8 | . | . |
| *Angiopteris evecta* | Eusporangiate | Marattiales | Marattiaceae | Angeve | 1KP; 10.1186/2047-217X-3-17 | 1 | 0 | 0 | 0 | 0 | 0 | 1 | 0 | NHCM | 10.1007/s13199-012-0185-z; www.myc | . | . |
| *Anomodon attenuatus* | Mosses | Hypnales | Anomodontaceae | Anoatt | 1KP; 10.1186/2047-217X-3-17 | 0 | 0 | 0 | 0 | 0 | 0 | 0 | 0 | QMWB | 10.1098/rstb.2000.0617 | . | . |
| *Anthoceros angustus* | Hornworts | Anthocerotales | Anthocerotaceae | Antang | 1KP; 10.1186/2047-217X-3-17 | 1 | 0 | 0 | 0 | 0 | 0 | 1 | 0 | IQJU | 10.1098/rspb.2013.0207 | . | . |
| *Apostasia shenzhenica* | Angiosperms | Asparagales | Orchidaceae | Aposhe | 10.1038/nature23897 | 0 | 0 | 1 | 0 | 0 | 0 | 1 | 0 | NA | . | . | . |
| *Arabidopsis halleri* | Angiosperms | Brassicales | Brassicaceae | Arahal | Unpublished - Phytozome | 0 | 0 | 0 | 0 | 0 | 0 | 0 | 0 | NA | . | . | . |
| *Arabidopsis lyrata* | Angiosperms | Brassicales | Brassicaceae | Aralyr | 10.1038/ng.807 | 0 | 0 | 0 | 0 | 0 | 0 | 0 | 0 | NA | . | . | . |
| *Arabidopsis thaliana* | Angiosperms | Brassicales | Brassicaceae | Aratha | 10.1093/nar/gkr1090 | 0 | 0 | 0 | 0 | 0 | 0 | 0 | 0 | NA | . | . | . |
| *Arachis duranensis* | Angiosperms | Fabales | Fabaceae | Aradur | 10.1038/ng.3517 | 1 | 1 | 0 | 0 | 0 | 0 | 1 | 0 | NA | . | . | . |
| *Arachis hypogaea* | Angiosperms | Fabales | Fabaceae | Arahyp | Unpublished - peanutbase.org | 1 | 1 | 0 | 0 | 0 | 0 | 1 | 0 | NA | Carling, D. E., Roncadori, R. W., & Huss | . | . |
| *Arachis ipaensis* | Angiosperms | Fabales | Fabaceae | Araipa | 10.1038/ng.3517 | 1 | 1 | 0 | 0 | 0 | 0 | 1 | 0 | NA | . | . | . |
| *Araucaria rulei* | Gymnosperms | Pinales | Araucariaceae | Ararul | 1KP; 10.1186/2047-217X-3-17 | 1 | 0 | 0 | 0 | 0 | 0 | 1 | 0 | XTZO | www.mycorrhizas.info/ | . | . |
| *Argyrochosma nivea* | Leptosporangiate | Polypodiales | Pteridaceae | Argniv | 1KP; 10.1186/2047-217X-3-17 | ND | 0 | 0 | 0 | 0 | 0 | ND | 0 | XDDT | . | . | . |
| *Asplenium nidus* | Leptosporangiate | Polypodiales | Aspleniaceae | Aspnid | 1KP; 10.1186/2047-217X-3-17 | 0 | 0 | 0 | 0 | 0 | 0 | 0 | 0 | PSKY | 10.1002/j.1537-2197.1992.tb13665.x | . | . |
| *Asplenium platyneuron* | Leptosporangiate | Polypodiales | Aspleniaceae | Asppla | 1KP; 10.1186/2047-217X-3-17 | 1 | 0 | 0 | 0 | 0 | 0 | 1 | 0 | KJZG | 10.1007/s00572-009-0234-5 | . | . |
| *Athrotaxis cupressoides* | Gymnosperms | Pinales | Cupressaceae | Athcup | 1KP; 10.1186/2047-217X-3-17 | 1 | 0 | 0 | 0 | 0 | 0 | 1 | 0 | XIRK | www.mycorrhizas.info/ | . | . |
| *Athyrium filix-femina* | Leptosporangiate | Polypodiales | Dryopteridaceae | Athfil | 1KP; 10.1186/2047-217X-3-17 | 1 | 0 | 0 | 0 | 0 | 0 | 1 | 0 | URCP | 10.2307/3792863 | . | . |
| *Atrichum angustatum* | Mosses | Polytrichales | Polytrichaceae | Atrang | 1KP; 10.1186/2047-217X-3-17 | 0 | 0 | 0 | 0 | 0 | 0 | 0 | 0 | ZTHV | 10.1098/rstb.2000.0617 | . | . |
| *Aulacomnium heterostichum* | Mosses | Aulacomniales | Aulacomniaceae | Aulhet | 1KP; 10.1186/2047-217X-3-17 | 0 | 0 | 0 | 0 | 0 | 0 | 0 | 0 | WNGH | 10.1098/rstb.2000.0617 | . | . |
| *Austrocedrus chilensis* | Gymnosperms | Pinales | Cupressaceae | Auschi | 1KP; 10.1186/2047-217X-3-17 | 1 | 0 | 0 | 0 | 0 | 0 | 1 | 0 | YYPE | 10.1007/s005720050207 | . | . |
| *Austrotaxus spicata* | Gymnosperms | Pinales | Taxaceae | Ausspi | 1KP; 10.1186/2047-217X-3-17 | 1 | 0 | 0 | 0 | 0 | 0 | 1 | 0 | BTTS | www.mycorrhizas.info/ | . | . |
| *Azolla cf. caroliniana* | Leptosporangiate | Salviniales | Salviniaceae | Azocar | 1KP; 10.1186/2047-217X-3-17 | 0 | 0 | 0 | 0 | 1 | 0 | 0 | 1 | CVEG | www.mycorrhizas.info/ | . | . |
| *Azolla filiculoides* | Leptosporangiate | Salviniales | Salviniaceae | Azofil | 10.1038/s41477-018-0188-8 | 0 | 0 | 0 | 0 | 1 | 0 | 0 | 1 | NA | 10.2307/2444993 | . | . |
| *Barbilophozia barbata* | Liverworts | Jungermanniales | Anastrophyllaceae | Barbar | 1KP; 10.1186/2047-217X-3-17 | 0 | 0 | 0 | 1 | 0 | 0 | 1 | 0 | OFTV | 10.1017/S0953756203008141 | . | . |
| *Bazzania trilobata* | Liverworts | Jungermanniales | Lepidoziaceae | Baztri | 1KP; 10.1186/2047-217X-3-17 | 0 | 0 | 0 | 1 | 0 | 0 | 1 | 0 | WZYK | . | . | Assumes Jungermanniales are not AM |
| *Begonia fuchsioides* | Angiosperms | Cucurbitales | Begoniaceae | Begfuc | 10.1126/science.aat1743 | 1 | 0 | 0 | 0 | 0 | 0 | 1 | 0 | NA | . | . | . |
| *Beta vulgaris ssp. vulgaris KWS2320* | Angiosperms | Caryophyllales | Amaranthaceae | Betvul | 10.1038/nature12817 | 0 | 0 | 0 | 0 | 0 | 0 | 0 | 0 | NA | . | . | . |
| *Blasia sp.* | Liverworts | Blasiales | Blasiaceae | Blasp. | 1KP; 10.1186/2047-217X-3-17 | 0 | 0 | 0 | 0 | 1 | 0 | 0 | 1 | AEXY | 10.1017/S0953756203008141 | . | . |
| *Blechnum spicant* | Leptosporangiate | Polypodiales | Blechnaceae | Blespi | 1KP; 10.1186/2047-217X-3-17 | 1 | 0 | 0 | 0 | 0 | 0 | 1 | 0 | RWYZ & VITX | . | . | . |
| *Boechera stricta* | Angiosperms | Brassicales | Brassicaceae | Boestr | Unpublished - Phytozome | 0 | 0 | 0 | 0 | 0 | 0 | 0 | 0 | NA | . | . | . |
| *Bolbitis repanda* | Leptosporangiate | Polypodiales | Dryopteridaceae | Bolrep | 1KP; 10.1186/2047-217X-3-17 | 0 | 0 | 0 | 0 | 0 | 0 | 0 | 0 | JBLI | 10.1007/s005720000076 | . | Two close-relative Bolbitis sp. species are not AMS |
| *Botrypus virginianus* | Eusporangiate | Ophioglossales | Ophioglossaceae | Botvir | 1KP; 10.1186/2047-217X-3-17 | 1 | 0 | 0 | 0 | 0 | 0 | 1 | 0 | BEGM | 10.1007/s00572-007-0137-2 | . | . |
| *Brachypodium distachyon* | Angiosperms | Poales | Poaceae | Bradis | 10.1038/nature08747 | 1 | 0 | 0 | 0 | 0 | 0 | 1 | 0 | NA | 10.1007/s00425-012-1677-z | . | . |
| *Brassica oleraceae capitata* | Angiosperms | Brassicales | Brassicaceae | Braolecap | 10.1038/ncomms4930 | 0 | 0 | 0 | 0 | 0 | 0 | 0 | 0 | NA | . | . | . |
| *Brassica rapa FPsc* | Angiosperms | Brassicales | Brassicaceae | Brarap | Unpublished - Phytozome | 0 | 0 | 0 | 0 | 0 | 0 | 0 | 0 | NA | . | . | . |
| *Bryum argenteum* | Mosses | Bryales | Bryaceae | Bryarg | 1KP; 10.1186/2047-217X-3-17 | 0 | 0 | 0 | 0 | 0 | 0 | 0 | 0 | JMXW | 10.1098/rstb.2000.0617 | . | . |
| *Buxbaumia aphylla* | Mosses | Buxbaumiales | Buxbaumiaceae | Buxaph | 1KP; 10.1186/2047-217X-3-17 | 0 | 0 | 0 | 0 | 0 | 0 | 0 | 0 | HRWG | 10.1098/rstb.2000.0617 | . | . |
| *Cajanus cajan* | Angiosperms | Fabales | Fabaceae | Cajcaj | 10.1038/nbt.2022. | 1 | 1 | 0 | 0 | 0 | 0 | 1 | 0 | NA | . | . | . |
| *Calliergon cordifolium* | Mosses | Hypnales | Calliergonaceae | Calcor | 1KP; 10.1186/2047-217X-3-17 | 0 | 0 | 0 | 0 | 0 | 0 | 0 | 0 | TAVP | 10.1098/rstb.2000.0617 | . | . |
| *Callitris gracilis* | Gymnosperms | Pinales | Cupressaceae | Calgra | 1KP; 10.1186/2047-217X-3-17 | 1 | 0 | 0 | 0 | 0 | 0 | 1 | 0 | IFLI | www.mycorrhizas.info/ | . | . |
| *Callitris macleayana* | Gymnosperms | Pinales | Cupressaceae | Calmac | 1KP; 10.1186/2047-217X-3-17 | 1 | 0 | 0 | 0 | 0 | 0 | 1 | 0 | RMMV | www.mycorrhizas.info/ | . | . |
| *Calocedrus decurrens* | Gymnosperms | Pinales | Cupressaceae | Caldec | 1KP; 10.1186/2047-217X-3-17 | 1 | 0 | 0 | 0 | 0 | 0 | 1 | 0 | FRPM | www.mycorrhizas.info/ | . | . |
| *Calypogeia fissa* | Liverworts | Jungermanniales | Calypogeiaceae | Calfis | 1KP; 10.1186/2047-217X-3-17 | 0 | 0 | 0 | 1 | 0 | 0 | 1 | 0 | RTMU | 10.1017/S0953756203008141 | . | . |
| *Capsella grandiflora* | Angiosperms | Brassicales | Brassicaceae | Capgra | 10.1038/ng.2669 | 0 | 0 | 0 | 0 | 0 | 0 | 0 | 0 | NA | . | . | . |
| *Capsella rubella* | Angiosperms | Brassicales | Brassicaceae | Caprub | 10.1038/ng.2669 | 0 | 0 | 0 | 0 | 0 | 0 | 0 | 0 | NA | . | . | . |
| *Capsicum annuum cvCM334* | Angiosperms | Solanales | Solanaceae | Capann | 10.1038/ng.2877 | 1 | 0 | 0 | 0 | 0 | 0 | 1 | 0 | NA | 10.1080/14620316.1988.11515864 | . | . |
| *Carica papaya* | Angiosperms | Brassicales | Caricaceae | Carpap | 10.1038/nature06856 | 1 | 0 | 0 | 0 | 0 | 0 | 1 | 0 | NA | 10.1078/0176-1617-00731 | . | . |
| *Castanea mollissima* | Angiosperms | Fagales | Fagaceae | Casmol | 10.1186/s12864-015-1942-1 | 1 | 0 | 0 | 0 | 0 | 1 | 1 | 1 | NA | . | 10.1007/s11104-008-9877-9 | . |
| *Castanospermum australe* | Angiosperms | Fabales | Fabaceae | Casaus | 10.1126/science.aat1743 | 1 | 0 | 0 | 0 | 0 | 0 | 1 | 0 | NA | 10.1007/s005720050008 | . | . |
| *Casuarina glauca* | Angiosperms | Fagales | Casuarinaceae | Casgla | 10.1126/science.aat1743 | 1 | 1 | 0 | 0 | 0 | 1 | 1 | 1 | NA | . | 10.1007/s11104-008-9877-10 | . |
| *Cathaya argyrophylla* | Gymnosperms | Pinales | Pinaceae | Catagr | 1KP; 10.1186/2047-217X-3-17 | 0 | 0 | 0 | 0 | 0 | 1 | 0 | 1 | NPRL | www.mycorrhizas.info/ | . | . |
| *Cavendishia cuatrecasasii* | Angiosperms | Ericales | Ericaceae | Cavcua | 1KP; 10.1186/2047-217X-3-17 | 0 | 0 | 0 | 1 | 0 | 0 | 1 | 0 | AVJK | . | . | . |
| *Cedrus libani* | Gymnosperms | Pinales | Pinaceae | Cedlib | 1KP; 10.1186/2047-217X-3-17 | 0 | 0 | 0 | 0 | 0 | 1 | 0 | 1 | GGEA | www.mycorrhizas.info/ | . | . |
| *Cephalotaxus harringtonia* | Gymnosperms | Pinales | Cephalotaxaceae | Cephar | 1KP; 10.1186/2047-217X-3-17 | 1 | 0 | 0 | 0 | 0 | 0 | 1 | 0 | GJTI,NVGZ,WYAJ | . | . | . |
| *Cephalotus follicularis* | Angiosperms | Oxalidales | Cephalotaceae | Cepfol | 10.1038/s41559-016-0059 | 0 | 0 | 0 | 0 | 0 | 0 | 0 | 0 | NA | . | . | . |
| *Ceratodon purpureus* | Mosses | Dicranales | Dicranaceae | Cerpur | 1KP; 10.1186/2047-217X-3-17 | 0 | 0 | 0 | 0 | 0 | 0 | 0 | 0 | FFPD | 10.1098/rstb.2000.0617 | . | . |
| *Cercis canadensis* | Angiosperms | Fabales | Caesalpiniaceae | Cercan | 10.1126/science.aat1743 | 1 | 0 | 0 | 0 | 0 | 0 | 1 | 0 | NA | . | . | . |
| *Chamaecrista fasciculata* | Angiosperms | Fabales | Fabaceae | Chafas | 10.1126/science.aat1743 | 1 | 1 | 0 | 0 | 0 | 0 | 1 | 0 | NA | . | . | . |
| *Chamaecyparis lawsoniana* | Gymnosperms | Pinales | Cupressaceae | Chalaw | 1KP; 10.1186/2047-217X-3-17 | 1 | 0 | 0 | 0 | 0 | 0 | 1 | 0 | AIGO | . | . | . |
| *Chenopodium quinoa* | Angiosperms | Caryophyllales | Amaranthaceae | Chequi | 10.1038/nature21370 | 0 | 0 | 0 | 0 | 0 | 0 | 0 | 0 | NA | . | . | . |
| *Cicer arietinum ICC4958* | Angiosperms | Fabales | Fabaceae | Cicari | 10.1038/srep12806 | 1 | 1 | 0 | 0 | 0 | 0 | 1 | 0 | NA | Chaturvedi, C., & Kumar, A. (1991). Noc | . | . |
| *Citrullus lanatus subsp. vulgaris 97103* | Angiosperms | Cucurbitales | Cucurbitaceae | Citlan | 10.1038/ng.2470 | 1 | 0 | 0 | 0 | 0 | 0 | 1 | 0 | NA | 10.1081/PLN-120022379 | . | . |
| *Citrus clementina* | Angiosperms | Sapindales | Rutaceae | Citcle | 10.1038/nbt.2906 | 1 | 0 | 0 | 0 | 0 | 0 | 1 | 0 | NA | . | . | . |
| *Citrus sinensis* | Angiosperms | Sapindales | Rutaceae | Citsin | 10.1038/nbt.2906 | 1 | 0 | 0 | 0 | 0 | 0 | 1 | 0 | NA | . | . | . |
| *Claopodium rostratum* | Mosses | Hypnales | Hypnaceae | Anocla | 1KP; 10.1186/2047-217X-3-17 | 0 | 0 | 0 | 0 | 0 | 0 | 0 | 0 | VBMM | 10.1098/rstb.2000.0617 | . | . |
| *Climacium dendroides* | Mosses | Hypnales | Climaciaceae | Cliden | 1KP; 10.1186/2047-217X-3-17 | 0 | 0 | 0 | 0 | 0 | 0 | 0 | 0 | MIRS | 10.1098/rstb.2000.0617 | . | . |
| *Conocephalum conicum* | Liverworts | Marchantiales | Conocephalaceae | Concon | 1KP; 10.1186/2047-217X-3-17 | 1 | 0 | 0 | 0 | 0 | 0 | 1 | 0 | ILBQ | 10.5586/asbp.1999.011 | . | . |
| *Crepidomanes venosum* | Leptosporangiate | Hymenophyllales | Hymenophyllaceae | Creven | 1KP; 10.1186/2047-217X-3-17 | 0 | 0 | 0 | 0 | 0 | 0 | 0 | 0 | TWFZ | 10.1007/s005720000076 | . | . |
| *Cryptogramma acrostichoides* | Leptosporangiate | Polypodiales | Pteridaceae | Cryacr | 1KP; 10.1186/2047-217X-3-17 | 0 | 0 | 0 | 0 | 0 | 0 | 0 | 0 | WQML | 10.1080/00275514.1982.12021584 | . | Another genus described as non AMS |
| *Cryptomeria japonica* | Gymnosperms | Pinales | Cupressaceae | Cryjap | 1KP; 10.1186/2047-217X-3-17 | 1 | 0 | 0 | 0 | 0 | 0 | 1 | 0 | DSXO,GMHZ,HOUF | 10.4489/KJM.2014.42.4.255 | . | . |
| *Cucumis melo* | Angiosperms | Cucurbitales | Cucurbitaceae | Cucmel | 10.1073/pnas.1205415109 | 1 | 0 | 0 | 0 | 0 | 0 | 1 | 0 | NA | . | . | . |
| *Cucumis sativus PI183967* | Angiosperms | Cucurbitales | Cucurbitaceae | Cucsat | 10.1038/ng.2801 | 1 | 0 | 0 | 0 | 0 | 0 | 1 | 0 | NA | 10.1016/0098-8472(91)90055-S | . | . |
| *Cucurbita maxima* | Angiosperms | Cucurbitales | Cucurbitaceae | Cucmax | 10.1016/j.molp.2017.09.003 | 1 | 0 | 0 | 0 | 0 | 0 | 1 | 0 | NA | . | . | . |
| *Cucurbita moschata* | Angiosperms | Cucurbitales | Cucurbitaceae | Cucmos | 10.1016/j.molp.2017.09.004 | 1 | 0 | 0 | 0 | 0 | 0 | 1 | 0 | NA | . | . | . |
| *Cucurbita pepo* | Angiosperms | Cucurbitales | Cucurbitaceae | Cucpep | Unpublished - cucurbitgenomics.org | 1 | 0 | 0 | 0 | 0 | 0 | 1 | 0 | NA | . | . | . |
| *Culcita macrocarpa* | Leptosporangiate | Cyatheales | Culcitaceae | Culmac | 1KP; 10.1186/2047-217X-3-17 | 1 | 0 | 0 | 0 | 0 | 0 | 1 | 0 | PNZO | 10.1111/jse.12227 | . | At the family level |
| *Cunninghamia lanceolata* | Gymnosperms | Pinales | Cupressaceae | Cunlan | 1KP; 10.1186/2047-217X-3-17 | 1 | 0 | 0 | 0 | 0 | 0 | 1 | 0 | ESYX,OUOI,ZQVF | . | . | . |
| *Cupressus dupreziana* | Gymnosperms | Pinales | Cupressaceae | Cupdup | 1KP; 10.1186/2047-217X-3-17 | 1 | 0 | 0 | 0 | 0 | 0 | 1 | 0 | QNGJ | www.mycorrhizas.info/ | . | . |
| *Cyathea (Alsophila) spinulosa* | Leptosporangiate | Cyatheales | Cyatheaceae | Cyaspi | 1KP; 10.1186/2047-217X-3-17 | 1 | 0 | 0 | 0 | 0 | 0 | 1 | 0 | GANB | 10.1007/s005720000076 | . | . |
| *Cycas micholitzii* | Gymnosperms | Cycadales | Cycadaceae | Cycmic | Unpublished - gymno-plaza / 1KP; 10.1186/2047-2 | 1 | 0 | 0 | 0 | ? | 0 | 1 | 0 | XZUY | www.mycorrhizas.info/ | . | . |
| *Cystopteris fragilis* | Leptosporangiate | Polypodiales | Cystopteridaceae | Cysfra | 1KP; 10.1186/2047-217X-3-17 | 1 | 0 | 0 | 0 | 0 | 0 | 1 | 0 | LHLE,XWDM,XXHP | . | . | . |
| *Cystopteris protrusa* | Leptosporangiate | Polypodiales | Cystopteridaceae | Cyspro | 1KP; 10.1186/2047-217X-3-17 | 1 | 0 | 0 | 0 | 0 | 0 | 1 | 0 | YOWV | Harley, J. L., & Harley, E. L. (1987). A ch | . | Assumes AM+ |
| *Cystopteris reevesiana* | Leptosporangiate | Polypodiales | Cystopteridaceae | Cysree | 1KP; 10.1186/2047-217X-3-17 | 1 | 0 | 0 | 0 | 0 | 0 | 1 | 0 | RICC | 10.1111/j.1469-8137.1987.tb00674.x | . | . |
| *Cystopteris utahensis* | Leptosporangiate | Polypodiales | Cystopteridaceae | Cysuta | 1KP; 10.1186/2047-217X-3-17 | 1 | 0 | 0 | 0 | 0 | 0 | 1 | 0 | HNDZ | . | . | . |
| *Dacrycarpus compactus* | Gymnosperms | Pinales | Podocarpaceae | Daccom | 1KP; 10.1186/2047-217X-3-17 | 1 | 0 | 0 | 0 | 0 | 0 | 1 | 0 | FMWZ | www.mycorrhizas.info/ | . | . |
| *Dacrydium balansae* | Gymnosperms | Pinales | Podocarpaceae | Dacbal | 1KP; 10.1186/2047-217X-3-17 | 1 | 0 | 0 | 0 | 0 | 0 | 1 | 0 | IZGN | www.mycorrhizas.info/ | . | . |
| *Datisca glomerata* | Angiosperms | Cucurbitales | Dasticaceae | Datglo | 10.1126/science.aat1743 | 1 | 1 | 0 | 0 | 0 | 0 | 1 | 0 | NA | . | . | . |
| *Daucus carota* | Angiosperms | Apiales | Apiaceae | Daucar | 10.1038/ng.3565 | 1 | 0 | 0 | 0 | 0 | 0 | 1 | 0 | NA | 10.1111/j.1469-8137.2004.01095.x | . | . |
| *Davallia fejeensis* | Leptosporangiate | Polypodiales | Davalliaceae | Davfej | 1KP; 10.1186/2047-217X-3-17 | ND | 0 | 0 | 0 | 0 | 0 | ND | 0 | OQWW | . | . | . |

| | | | | | | | | | | | | | | | | | |
|---|---|---|---|---|---|---|---|---|---|---|---|---|---|---|---|---|---|
| *Dendrobium catenatum* | Angiosperms | Asparagales | Orchidaceae | Dencat | 10.1038/nature23897 | 0 | 0 | 1 | 0 | 0 | 0 | 1 | 0 | NA | . | . | . |
| *Dendrolycopodium obscurum* | Lycophytes | Lycopodiales | Lycopodiaceae | Denobs | 1KP; 10.1186/2047-217X-3-17 | 0 | 0 | 0 | 0 | 0 | 0 | 0 | 0 | XNXF | 10.1111/nph.13221 | . | . |
| *Dennstaedtia davallioides* | Leptosporangiate | Polypodiales | Dennstaedtiaceae | Dendav | 1KP; 10.1186/2047-217X-3-17 | 1 | 0 | 0 | 0 | 0 | 0 | 1 | 0 | MTGC | 10.1007/s005720000076 | . | Based on 2 other species |
| *Deparia lobato-crenata* | Leptosporangiate | Polypodiales | Athyriaceae | Deplob | 1KP; 10.1186/2047-217X-3-17 | ND | 0 | 0 | 0 | 0 | 0 | ND | ND | FCHS | . | . | . |
| *Dianthus caryophyllus* | Angiosperms | Caryophyllales | Caryophyllaceae | Diacar | 10.1093/dnares/dst053 | 0 | 0 | 0 | 0 | 0 | 0 | 0 | 0 | NA | 10.1139/b97-110 | . | . |
| *Dicranum scoparium* | Mosses | Dicranales | Dicranaceae | Dicsco | 1KP; 10.1186/2047-217X-3-17 | 0 | 0 | 0 | 0 | 0 | 0 | 0 | 0 | NGTD | 10.1098/rstb.2000.0617 | . | . |
| *Didymochlaena truncatula* | Leptosporangiate | Polypodiales | Hypodematiaceae | Didtru | 1KP; 10.1186/2047-217X-3-17 | ND | 0 | 0 | 0 | 0 | 0 | ND | ND | RFRB | . | . | . |
| *Dioon edule* | Gymnosperms | Cycadales | Zamiaceae | Dioedu | 1KP; 10.1186/2047-217X-3-17 | 1 | 0 | 0 | 0 | 0 | 0 | 1 | 0 | WLIC | Fisher, J. B., & Vovides, A. P. (2004). My | . | . |
| *Diphasiastrum digitatum* | Lycophytes | Lycopodiales | Lycopodiaceae | Dipdig | 1KP; 10.1186/2047-217X-3-17 | 1 | 0 | 0 | 0 | 0 | 0 | 1 | 0 | WAFT | Harley, J. L., & Harley, E. L. (1987). A ch | . | Assumed for the genus |
| *Diphyscium foliosum* | Mosses | Diphysciales | Diphysciaceae | Dipfol | 1KP; 10.1186/2047-217X-3-17 | 0 | 0 | 0 | 0 | 0 | 0 | 0 | 0 | AWOI | 10.1098/rstb.2000.0617 | . | . |
| *Diplazium wichurae* | Leptosporangiate | Polypodiales | Athyriaceae | Dipwic | 1KP; 10.1186/2047-217X-3-17 | 1 | 0 | 0 | 0 | 0 | 0 | 1 | 0 | UFJN | 10.1007/s005720000076 | . | . |
| *Dipteris conjugata* | Leptosporangiate | Gleicheniales | Dipteridaceae | Dipcon | 1KP; 10.1186/2047-217X-3-17 | ND | 0 | 0 | 0 | 0 | 0 | ND | ND | MEKP | . | . | . |
| *Discaria trinervis* | Angiosperms | Rosales | Rhamnaceae | Distri | 10.1126/science.aat1743 | 1 | 1 | 0 | 0 | 0 | 0 | 1 | 0 | NA | . | . | . |
| *Diselma archeri* | Gymnosperms | Pinales | Cupressaceae | Disarc | 1KP; 10.1186/2047-217X-3-17 | 1 | 0 | 0 | 0 | 0 | 0 | 1 | 0 | GKCZ | www.mycorrhizas.info/ | . | . |
| *Dryas drummondii* | Angiosperms | Rosales | Rosaceae | Drydru | 10.1126/science.aat1743 | 1 | 1 | 0 | 0 | 0 | 0 | 1 | 0 | NA | . | . | . |
| *Encalypta streptocarpa* | Mosses | Funariales | Encalyptaceae | Encstr | 1KP; 10.1186/2047-217X-3-17 | 0 | 0 | 0 | 0 | 0 | 0 | 0 | 0 | KEFD | 10.1098/rstb.2000.0617 | . | . |
| *Encephalartos barteri* | Gymnosperms | Cycadales | Zamiaceae | Encbar | 1KP; 10.1186/2047-217X-3-17 | 1 | 0 | 0 | 0 | 0 | 0 | 1 | 0 | GNQG | www.mycorrhizas.info/ | . | Several species of the familly formed AMS |
| *Ephedra sinica* | Gymnosperms | Ephedrales | Ephedraceae | Ephsin | 1KP; 10.1186/2047-217X-3-17 | 1 | 0 | 0 | 0 | 0 | 0 | 1 | 0 | VDAO | www.mycorrhizas.info/ | . | . |
| *Equisetum diffusum* | Eusporangiate | Equisetales | Equisetaceae | Equdif | 1KP; 10.1186/2047-217X-3-17 | 1 | 0 | 0 | 0 | 0 | 0 | 1 | 0 | CAPN | 10.1016/S0007-1536(85)80202-3 | . | . |
| *Equisetum hyemale* | Eusporangiate | Equisetales | Equisetaceae | Equhye | 1KP; 10.1186/2047-217X-3-17 | 1 | 0 | 0 | 0 | 0 | 0 | 1 | 0 | JVSZ | 10.1016/S0007-1536(85)80202-3 | . | . |
| *Eutrema salsugineum* | Angiosperms | Brassicales | Brassicaceae | Eutsal | 10.3389/fpls.2013.00046 | 0 | 0 | 0 | 0 | 0 | 0 | 0 | 0 | NA | . | . | . |
| *Falcatifolium taxoides* | Gymnosperms | Pinales | Podocarpaceae | Faltax | 1KP; 10.1186/2047-217X-3-17 | 1 | 0 | 0 | 0 | 0 | 0 | 1 | 0 | PLYX,QHBI,ROWR | . | . | . |
| *Fokienia hodginsii* | Gymnosperms | Pinales | Cupressaceae | Fokhod | 1KP; 10.1186/2047-217X-3-17 | 1 | 0 | 0 | 0 | 0 | 0 | 1 | 0 | UEVI | www.mycorrhizas.info/ | . | . |
| *Fontinalis antipyretica* | Mosses | Isobryales | Fontinalaceae | Fonant | 1KP; 10.1186/2047-217X-3-17 | 0 | 0 | 0 | 0 | 0 | 0 | 0 | 0 | DHWX | 10.1098/rstb.2000.0617 | . | . |
| *Fragaria vesca* | Angiosperms | Rosales | Rosaceae | Fraves | 10.1093/gigascience/gix124 | 1 | 0 | 0 | 0 | 0 | 0 | 1 | 0 | NA | Harley, J. L., & Harley, E. L. (1987). A ch | . | . |
| *Fragaria x ananassa* | Angiosperms | Rosales | Rosaceae | Fraana | Unpublished - rosaceae.org | 1 | 0 | 0 | 0 | 0 | 0 | 1 | 0 | NA | 10.1139/b04-007 | . | . |
| *Fraxinus excelsior* | Angiosperms | Lamiales | Oleaceae | Fraexc | doi:10.1038/nature20786 | 1 | 0 | 0 | 0 | 0 | 0 | 1 | 0 | NA | 10.1080/014904599270767 | . | . |
| *Frullania sp.* | Liverworts | Porellales | Frullaniaceae | Frusp. | 1KP; 10.1186/2047-217X-3-17 | ND | 0 | 0 | 0 | 0 | 0 | ND | 0 | TGKW | . | . | . |
| *Frullania spp.* | Liverworts | Porellales | Frullaniaceae | Fruspp | 1KP; 10.1186/2047-217X-3-17 | ND | 0 | 0 | 0 | 0 | 0 | ND | 0 | CHJJ | . | . | . |
| *Gaga arizonica* | Leptosporangiate | Polypodiales | Pteridaceae | Gagari | 1KP; 10.1186/2047-217X-3-17 | ND | 0 | 0 | 0 | 0 | 0 | ND | ND | DCDT | . | . | . |
| *Ginkgo biloba* | Gymnosperms | Ginkgoales | Ginkgoaceae | Ginbil | 10.1186/s13742-016-0154-1 / 1KP; 10.1186/2047- | 1 | 0 | 0 | 0 | 0 | 0 | 1 | 0 | SGTW | . | . | . |
| *Glycine max* | Angiosperms | Fabales | Fabaceae | Glymax | 10.1038/nature08670 | 1 | 1 | 0 | 0 | 0 | 0 | 1 | 0 | NA | 10.1046/j.0028-646x.2001.00187.x | . | . |
| *Glyptostrobus pensilis* | Gymnosperms | Pinales | Cupressaceae | Glypen | 1KP; 10.1186/2047-217X-3-17 | 1 | 0 | 0 | 0 | 0 | 0 | 1 | 0 | OXGJ | www.mycorrhizas.info/ | . | . |
| *Gnetum montanum* | Gymnosperms | Gnetales | Gnetaceae | Gnemon | 10.5061/dryad.0vm37 / 1KP; 10.1186/2047-217X- | 0 | 0 | 0 | 0 | 0 | 1 | 0 | 1 | GTHK | . | . | . |
| *Gossypium raimondii* | Angiosperms | Malvales | Malvaceae | Gosrai | 10.1038/nature11798 | 1 | 0 | 0 | 0 | 0 | 0 | 1 | 0 | NA | . | . | . |
| *Gymnocarpium dryopteris* | Leptosporangiate | Polypodiales | Cystopteridaceae | Gymdry | 1KP; 10.1186/2047-217X-3-17 | 1 | 0 | 0 | 0 | 0 | 0 | 1 | 0 | HEGQ | 10.2307/3792863 | . | . |
| *Halocarpus bidwillii* | Gymnosperms | Pinales | Podocarpaceae | Halbid | 1KP; 10.1186/2047-217X-3-17 | 1 | 0 | 0 | 0 | 0 | 0 | 1 | 0 | OWFC | www.mycorrhizas.info/ | . | . |
| *Hedwigia ciliata* | Mosses | Hedwigiales | Hedwigiaceae | Hedcil | 1KP; 10.1186/2047-217X-3-17 | 0 | 0 | 0 | 0 | 0 | 0 | 0 | 0 | YWNF | 10.1098/rstb.2000.0617 | . | . |
| *Helianthus annuus* | Angiosperms | Asterales | Asteraceae | Helann | 10.1038/nature22380 | 1 | 0 | 0 | 0 | 0 | 0 | 1 | 0 | NA | 10.1007/BF00011797 | . | . |
| *Homalosorus pycnocarpos* | Leptosporangiate | Polypodiales | Aspleniaceae | Hompyc | 1KP; 10.1186/2047-217X-3-17 | ND | 0 | 0 | 0 | 0 | 0 | ND | ND | OCZL | . | . | . |
| *Hordeum vulgare* | Angiosperms | Poales | Poaceae | Horvul | 10.1038/nature22043 / 10.1038/sdata.2017.44 | 1 | 0 | 0 | 0 | 0 | 0 | 1 | 0 | NA | Champawat, R. S., Pathak, V. N., & Krish | . | . |
| *Humulus lupulus* | Angiosperms | Rosales | Cannabaceae | Humlup | 10.1093/pcp/pcu169 | 1 | 0 | 0 | 0 | 0 | 0 | 1 | 0 | NA | Harley, J. L., & Harley, E. L. (1987). A ch | . | . |
| *Huperzia lucidula* | Lycophytes | Lycopodiales | Lycopodiaceae | Hupluc | 1KP; 10.1186/2047-217X-3-17 | 1 | 0 | 0 | 0 | 0 | 0 | 1 | 0 | GKAG | 10.2307/2444993; Harley, J. L., & Harle | . | At the genus level |
| *Huperzia myrsinites* | Lycophytes | Lycopodiales | Lycopodiaceae | Hupmyr | 1KP; 10.1186/2047-217X-3-17 | 1 | 0 | 0 | 0 | 0 | 0 | 1 | 0 | CBAE | 10.2307/2444993; Harley, J. L., & Harle | . | At the genus level |
| *Huperzia selago* | Lycophytes | Lycopodiales | Lycopodiaceae | Hupsel | 1KP; 10.1186/2047-217X-3-17 | 1 | 0 | 0 | 0 | 0 | 0 | 1 | 0 | GTUO,NYBX,YHZW | Harley, J. L., & Harley, E. L. (1987). A ch | . | . |
| *Huperzia squarrosa* | Lycophytes | Lycopodiales | Lycopodiaceae | Hupsqu | 1KP; 10.1186/2047-217X-3-17 | 1 | 0 | 0 | 0 | 0 | 0 | 1 | 0 | GAON | 10.2307/2444993; Harley, J. L., & Harle | . | At the order level |
| *Hymenophyllum bivalve* | Leptosporangiate | Hymenophyllales | Hymenophyllaceae | Hymbiv | 1KP; 10.1186/2047-217X-3-17 | 0 | 0 | 0 | 0 | 0 | 0 | 0 | 0 | QIAD | 10.1111/jse.12227 | . | At the order level |
| *Hymenophyllum cupressiforme* | Leptosporangiate | Hymenophyllales | Hymenophyllaceae | Hymcup | 1KP; 10.1186/2047-217X-3-17 | 0 | 0 | 0 | 0 | 0 | 0 | 0 | 0 | TRPJ | 10.1111/jse.12227 | . | At the order level |
| *Isoetes sp.* | Lycophytes | Isoëtales | Isoëtaceae | Isoesp. | 1KP; 10.1186/2047-217X-3-17 | ND | 0 | 0 | 0 | 0 | 0 | ND | 0 | PYHZ | 10.1016/0304-3770(85)90052-X | . | Genus seems AMS but Isoetes lacustri is probably non-AMS: 10.1016/j.aquabot.2011.02.003; Nature 268, 232–233 (1977) |
| *Isoetes tegetiformans* | Lycophytes | Isoëtales | Isoëtaceae | Isoteg | 1KP; 10.1186/2047-217X-3-17 | ND | 0 | 0 | 0 | 0 | 0 | ND | 0 | PKOX | 10.1016/0304-3770(85)90052-X | . | Genus seems AMS but Isoetes lacustri is probably non-AMS: 10.1016/j.aquabot.2011.02.003; Nature 268, 232–233 (1977) |
| *Juglans regia* | Angiosperms | Fagales | Juglandaceae | Jugreg | 10.1111/tpj.13207 | 1 | 0 | 0 | 0 | 0 | 0 | 1 | 0 | NA | 10.1007/BF02913007 | . | . |
| *Juniperus scopulorum* | Gymnosperms | Pinales | Cupressaceae | Junsco | 1KP; 10.1186/2047-217X-3-17 | 1 | 0 | 0 | 0 | 0 | 0 | 1 | 0 | XMGP | www.mycorrhizas.info/ | . | . |
| *Keteleeria evelyniana* | Gymnosperms | Pinales | Pinaceae | Keteve | 1KP; 10.1186/2047-217X-3-17 | 0 | 0 | 0 | 0 | 0 | 1 | 0 | 1 | JUWL | www.mycorrhizas.info/ | . | . |
| *Lagarostrobos franklinii* | Gymnosperms | Pinales | Podocarpaceae | Lagfra | 1KP; 10.1186/2047-217X-3-17 | 1 | 0 | 0 | 0 | 0 | 0 | 1 | 0 | ZQWM | www.mycorrhizas.info/ | . | . |
| *Lagenaria siceraria* | Angiosperms | Cucurbitales | Cucurbitaceae | Lagsic | 10.1111/tpj.13722 | 1 | 0 | 0 | 0 | 0 | 0 | 1 | 0 | NA | . | . | . |
| *Larix speciosa* | Gymnosperms | Pinales | Pinaceae | Larspe | 1KP; 10.1186/2047-217X-3-17 | 0 | 0 | 0 | 0 | 0 | 1 | 0 | 1 | WVWN | www.mycorrhizas.info/ | . | . |
| *Ledum palustre* | Angiosperms | Ericales | Ericaceae | Ledpal | 1KP; 10.1186/2047-217X-3-17 | 0 | 0 | 0 | 1 | 0 | 0 | 1 | 0 | WXVX | . | . | . |
| *Leiosporoceros dussii* | Hornworts | Leiosporocerotales | Leiosporocerotaceae | Leidus | 1KP; 10.1186/2047-217X-3-17 | 0 | 0 | 0 | 0 | 0 | 0 | 0 | 0 | FANS | 10.1098/rspb.2013.0207 | . | . |
| *Lepidothamnus sp.* | Gymnosperms | Pinales | Podocarpaceae | Lepsp. | 1KP; 10.1186/2047-217X-3-17 | 1 | 0 | 0 | 0 | 0 | 0 | 1 | 0 | BBDD | 10.1007/s11104-013-1609-0 | . | . |
| *Leucobryum albidum* | Mosses | Dicranales | Leucobryaceae | Leualb | 1KP; 10.1186/2047-217X-3-17 | 0 | 0 | 0 | 0 | 0 | 0 | 0 | 0 | VMXJ | 10.1098/rstb.2000.0617 | . | . |
| *Leucobryum glaucum* | Mosses | Dicranales | Leucobryaceae | Leugla | 1KP; 10.1186/2047-217X-3-17 | 0 | 0 | 0 | 0 | 0 | 0 | 0 | 0 | RGKI | 10.1098/rstb.2000.0617 | . | . |
| *Leucodon brachypus* | Mosses | Hypnales | Leucodontaceae | Leubra | 1KP; 10.1186/2047-217X-3-17 | 0 | 0 | 0 | 0 | 0 | 0 | 0 | 0 | ZACW | 10.1098/rstb.2000.0617 | . | . |
| *Leucodon julaceus* | Mosses | Hypnales | Leucodontaceae | Leujul | 1KP; 10.1186/2047-217X-3-17 | 0 | 0 | 0 | 0 | 0 | 0 | 0 | 0 | IGUH | 10.1098/rstb.2000.0617 | . | . |
| *Leucostegia immersa* | Leptosporangiate | Polypodiales | Hypodematiaceae | Leuimm | 1KP; 10.1186/2047-217X-3-17 | ND | 0 | 0 | 0 | 0 | 0 | ND | ND | WGTU | . | . | . |
| *Lindsaea linearis* | Leptosporangiate | Polypodiales | Lindsaeaceae | Linlin | 1KP; 10.1186/2047-217X-3-17 | 1 | 0 | 0 | 0 | 0 | 0 | 1 | 0 | NOKI | 10.1080/0028825X.1976.10428891 | . | . |
| *Lindsaea microphylla* | Leptosporangiate | Polypodiales | Lindsaeaceae | Linmic | 1KP; 10.1186/2047-217X-3-17 | 1 | 0 | 0 | 0 | 0 | 0 | 1 | 0 | YIXP | 10.1080/0028825X.1976.10428891 | . | At the genus level |
| *Loeskeobryum brevirostre* | Mosses | Hypnales | Hylocomiaceae | Loebre | 1KP; 10.1186/2047-217X-3-17 | 0 | 0 | 0 | 0 | 0 | 0 | 0 | 0 | WSPM | 10.1098/rstb.2000.0617 | . | . |
| *Lotus japonicus* | Angiosperms | Fabales | Fabaceae | Lotjap | 10.1093/dnares/dsn008 | 1 | 1 | 0 | 0 | 0 | 0 | 1 | 0 | NA | . | . | . |
| *Lunularia cruciata* | Liverworts | Marchantiales | Lunulariaceae | Luncru | 1KP; 10.1186/2047-217X-3-17 | 1 | 0 | 0 | 0 | 0 | 0 | 1 | 0 | TXVB | 10.3732/ajb.94.11.1756 | . | . |
| *Lupinus angustifolius* | Angiosperms | Fabales | Fabaceae | Lupang | 10.1111/pbi.12615 | 0 | 1 | 0 | 0 | 0 | 0 | 1 | 0 | NA | . | . | . |
| *Lycopodiella appressa* | Lycophytes | Lycopodiales | Lycopodiaceae | Lycapp | 1KP; 10.1186/2047-217X-3-17 | 1 | 0 | 0 | 0 | 0 | 0 | 1 | 0 | ULKT | 10.1007/s00572-004-0314-5 | . | At the genus level |
| *Lycopodium annotinum* | Lycophytes | Lycopodiales | Lycopodiaceae | Lycann | 1KP; 10.1186/2047-217X-3-17 | 1 | 0 | 0 | 0 | 0 | 0 | 1 | 0 | ENQF | Harley, J. L., & Harley, E. L. (1987). A ch | . | . |
| *Lycopodium deuterodensum* | Lycophytes | Lycopodiales | Lycopodiaceae | Lycdeu | 1KP; 10.1186/2047-217X-3-17 | 1 | 0 | 0 | 0 | 0 | 0 | 1 | 0 | PQTO | Harley, J. L., & Harley, E. L. (1987). A ch | . | . |
| *Lygodium japonicum* | Leptosporangiate | Schizaeales | Lygodiaceae | Lygjap | 1KP; 10.1186/2047-217X-3-17 | 1 | 0 | 0 | 0 | 0 | 0 | 1 | 0 | PBUU | 10.1007/s13199-012-0185-z | . | . |
| *Malus domestica* | Angiosperms | Rosales | Rosaceae | Maldom | Unpublished - rosaceae.org | 1 | 0 | 0 | 0 | 0 | 0 | 1 | 0 | NA | 10.1007/s00572-002-0194-5 | . | . |
| *Manihot esculenta* | Angiosperms | Malpighiales | Euphorbiaceae | Manesc | 10.1038/nbt.3535 | 1 | 0 | 0 | 0 | 0 | 0 | 1 | 0 | NA | 10.1007/BF00206771 | . | . |
| *Manoao colensoi* | Gymnosperms | Pinales | Podocarpaceae | Mancol | 1KP; 10.1186/2047-217X-3-17 | 1 | 0 | 0 | 0 | 0 | 0 | 1 | 0 | CDFR | www.mycorrhizas.info/ | . | . |
| *Marattia sp.* | Eusporangiate | Marattiales | Marattiaceae | Marsp. | 1KP; 10.1186/2047-217X-3-17 | 1 | 0 | 0 | 0 | 0 | 0 | 1 | 0 | UXCS | www.jstor.org/stable/2428708 | . | . |
| *Marchantia emarginata* | Liverworts | Marchantiales | Marchantiaceae | Marema | 1KP; 10.1186/2047-217X-3-17 | 1 | 0 | 0 | 0 | 0 | 0 | 1 | 0 | TFYI | this study | . | . |
| *Marchantia paleacea* | Liverworts | Marchantiales | Marchantiaceae | Marpal | This study / 1KP; 10.1186/2047-217X-3-17 | 1 | 0 | 0 | 0 | 0 | 0 | 1 | 0 | HMHL,IHWO,LFVP | 10.1038/ncomms1105; this study | . | . |
| *Marchantia polymorpha spp ruderalis* | Liverworts | Marchantiales | Marchantiaceae | Marpol | 10.1016/j.cell.2017.09.030 / 1KP; 10.1186/2047-2 | 0 | 0 | 0 | 0 | 0 | 0 | 0 | 0 | JPYU | this study | . | . |
| *Medicago truncatula* | Angiosperms | Fabales | Fabaceae | Medtru | 10.1038/s41477-018-0286-7 | 1 | 1 | 0 | 0 | 0 | 0 | 1 | 0 | NA | . | . | . |
| *Megaceros flagellaris* | Hornworts | Dendrocerotales | Dendrocerotaceae | Megfla | 1KP; 10.1186/2047-217X-3-17 | 0 | 0 | 0 | 0 | 0 | 0 | 0 | 0 | UCRN | . | . | . |
| *Metasequoia glyptostroboides* | Gymnosperms | Pinales | Cupressaceae | Metgly | 1KP; 10.1186/2047-217X-3-17 | 1 | 0 | 0 | 0 | 0 | 0 | 1 | 0 | NRXL | 10.1017/S0953756200003221; Stockey | . | . |
| *Metzgeria crassipilis* | Liverworts | Metzgeriales | Metzgeriaceae | Metcra | 1KP; 10.1186/2047-217X-3-17 | 0 | 0 | 0 | 0 | 0 | 0 | 0 | 0 | NRWZ | . | . | Assumes Metzgeriales are not AM |
| *Microbiota decussata* | Gymnosperms | Pinales | Cupressaceae | Micdec | 1KP; 10.1186/2047-217X-3-17 | 1 | 0 | 0 | 0 | 0 | 0 | 1 | 0 | XQSG | www.mycorrhizas.info/ | . | . |
| *Microcachrys tetragona* | Gymnosperms | Pinales | Podocarpaceae | Mictet | 1KP; 10.1186/2047-217X-3-17 | 1 | 0 | 0 | 0 | 0 | 0 | 1 | 0 | MHGD | www.mycorrhizas.info/ | . | . |
| *Mimosa pudica* | Angiosperms | Fabales | Fabaceae | Mimpud | 10.1126/science.aat1743 | 1 | 1 | 0 | 0 | 0 | 0 | 1 | 0 | NA | 10.2307/2444993 | . | . |
| *mixed species* | Liverworts | NA | NA | Mixspe | 1KP; 10.1186/2047-217X-3-17 | ND | 0 | 0 | 0 | 0 | 0 | ND | ND | NWQC | . | . | . |
| *Momordica charantia* | Angiosperms | Cucurbitales | Cucurbitaceae | Momcha | Unpublished - NCBI | 1 | 0 | 0 | 0 | 0 | 0 | 1 | 0 | NA | . | . | . |
| *Monoclea gottschei* | Liverworts | Marchantiales | Monocleaceae | Mongot | 1KP; 10.1186/2047-217X-3-17 | 1 | 0 | 0 | 0 | 0 | 0 | 1 | 0 | TFDQ | 10.3732/ajb.94.11.1756 | . | . |
| *Monotropa uniflora* | Angiosperms | Ericales | Ericaceae | Monuni | 1KP; 10.1186/2047-217X-3-17 | 0 | 0 | 0 | 1 | 0 | 0 | 1 | 0 | LRTN | . | . | . |
| *Morus notabilis* | Angiosperms | Rosales | Moraceae | Mornot | 10.1038/ncomms3445 | 1 | 0 | 0 | 0 | 0 | 0 | 1 | 0 | NA | . | . | . |
| *Musa acuminata* | Angiosperms | Zingiberales | Zingiberaceae | Musacu | 10.1093/database/bat035 | 1 | 0 | 0 | 0 | 0 | 0 | 1 | 0 | NA | 10.1007/BF00017688 | . | . |
| *Myriopteris rufa* | Leptosporangiate | Polypodiales | Pteridaceae | Myrruf | 1KP; 10.1186/2047-217X-3-17 | 1 | 0 | 0 | 0 | 0 | 0 | 1 | 0 | GSXD | www.ciidirsinaloa.com.mx/Simbiosis/4 | . | At the genus level |
| *Nageia nagi* | Gymnosperms | Pinales | Podocarpaceae | Nagnag | 1KP; 10.1186/2047-217X-3-17 | 1 | 0 | 0 | 0 | 0 | 0 | 1 | 0 | UUJS | www.mycorrhizas.info/ | . | . |
| *Neckera douglasii* | Mosses | Hypnales | Neckeraceae | Necdou | 1KP; 10.1186/2047-217X-3-17 | 0 | 0 | 0 | 0 | 0 | 0 | 0 | 0 | TMAJ | 10.1098/rstb.2000.0617 | . | . |
| *Nelumbo nucifera* | Angiosperms | Proteales | Nelumbonaceae | Nelnuc | 10.1186/gb-2013-14-5-r41 | 0 | 0 | 0 | 0 | 0 | 0 | 0 | 0 | NA | . | . | . |
| *Neocallitropsis pancheri* | Gymnosperms | Pinales | Cupressaceae | Neopan | 1KP; 10.1186/2047-217X-3-17 | 1 | 0 | 0 | 0 | 0 | 0 | 1 | 0 | JDQB | www.mycorrhizas.info/ | . | . |
| *Nephrolepis exaltata* | Leptosporangiate | Polypodiales | Nephrolepidaceae | Nepexa | 1KP; 10.1186/2047-217X-3-17 | 1 | 0 | 0 | 0 | 0 | 0 | 1 | 0 | NWWI | 10.1007/s13199-012-0185-z | . | . |
| *Nicotiana benthamiana* | Angiosperms | Solanales | Solanaceae | Nicben | 10.1094/MPMI-06-12-0148-TA | 1 | 0 | 0 | 0 | 0 | 0 | 1 | 0 | NA | . | . | . |
| *Nissolia schottii* | Angiosperms | Fabales | Fabaceae | Nissch | 10.1126/science.aat1743 | 1 | 0 | 0 | 0 | 0 | 0 | 1 | 0 | NA | . | . | . |
| *Nothoceros aenigmaticus* | Hornworts | Dendrocerotales | Dendrocerotaceae | Notaen | 1KP; 10.1186/2047-217X-3-17 | ND | 0 | 0 | 0 | 0 | 0 | ND | ND | DXOU | . | . | . |
| *Nothoceros vincentianus* | Hornworts | Dendrocerotales | Dendrocerotaceae | Notvin | 1KP; 10.1186/2047-217X-3-17 | 1 | 0 | 0 | 0 | 0 | 0 | 1 | 0 | TCBC | 10.1098/rspb.2013.0207 | . | . |

| | | | | | | | | | | | | | | | | | |
|---|---|---|---|---|---|---|---|---|---|---|---|---|---|---|---|---|---|
| *Notholaena montieliae* | Leptosporangiate | Polypodiales | Pteridaceae | Notmon | 1KP; 10.1186/2047-217X-3-17 | ND | 0 | 0 | 0 | 0 | 0 | ND | ND | YCKE | . | . | . |
| *Nothotsuga longibracteata* | Gymnosperms | Pinales | Pinaceae | Notlon | 1KP; 10.1186/2047-217X-3-17 | 0 | 0 | 0 | 0 | 0 | 1 | 0 | 1 | AREG | www.mycorrhizas.info/ | . | . |
| *Odontoschisma prostratum* | Liverworts | Jungermanniales | Cephaloziaceae | Odopro | 1KP; 10.1186/2047-217X-3-17 | 0 | 0 | 0 | 1 | 0 | 0 | 1 | 0 | YBQN | . | . | Assumes Jungermanniales are not AM |
| *Onoclea sensibilis* | Leptosporangiate | Polypodiales | Onocleaceae | Onosen | 1KP; 10.1186/2047-217X-3-17 | 1 | 0 | 0 | 0 | 0 | 0 | 1 | 0 | HTFH | 10.2307/3792863 | . | . |
| *Ophioglossum petiolatum* | Eusporangiate | Ophioglossales | Ophioglossaceae | Ophpet | 1KP; 10.1186/2047-217X-3-17 | 1 | 0 | 0 | 0 | 0 | 0 | 1 | 0 | WTJG | www.mycorrhizas.info/ | . | . |
| *Ophioglossum vulgatum* | Eusporangiate | Ophioglossales | Ophioglossaceae | Ophvul | 1KP; 10.1186/2047-217X-3-17 | 1 | 0 | 0 | 0 | 0 | 0 | 1 | 0 | QHVS | 10.1111/nph.13263 | . | . |
| *Orthotrichum lyellii* | Mosses | Orthotrichales | Orthotrichaceae | Ortlye | 1KP; 10.1186/2047-217X-3-17 | 0 | 0 | 0 | 0 | 0 | 0 | 0 | 0 | CMEQ | 10.1098/rstb.2000.0617 | . | . |
| *Oryza sativa* | Angiosperms | Poales | Poaceae | Orysat | 10.1093/nar/gkl976 | 1 | 0 | 0 | 0 | 0 | 0 | 1 | 0 | NA | 10.1016/S2095-3119(15)61241-2 | . | . |
| *Osmunda javanica* | Leptosporangiate | Osmundales | Osmundaceae | Osmjav | 1KP; 10.1186/2047-217X-3-17 | 1 | 0 | 0 | 0 | 0 | 0 | 1 | 0 | VIBO | www.mycorrhizas.info/ | . | . |
| *Osmunda regalis* | Leptosporangiate | Osmundales | Osmundaceae | Osmreg | 1KP; 10.1186/2047-217X-3-17 | 1 | 0 | 0 | 0 | 0 | 0 | 1 | 0 | YKSS | 10.1038/ncomms1831 | . | . |
| *Osmunda sp.* | Leptosporangiate | Osmundales | Osmundaceae | Osmsp. | 1KP; 10.1186/2047-217X-3-17 | 1 | 0 | 0 | 0 | 0 | 0 | 1 | 0 | UOMY | . | . | Based on close-relative species |
| *Osmundastrum cinnamomeum* | Leptosporangiate | Osmundales | Osmundaceae | Osmcin | 1KP; 10.1186/2047-217X-3-17 | 1 | 0 | 0 | 0 | 0 | 0 | 1 | 0 | BIVQ | 10.2307/3792863 | . | . |
| *Pallavicinia lyellii* | Liverworts | Pallaviciniales | Pallaviciniaceae | Pallye | 1KP; 10.1186/2047-217X-3-17 | 1 | 0 | 0 | 0 | 0 | 0 | 1 | 0 | YFGP | Nebel, M., et al. Frontiers in Basidiomy | . | . |
| *Papuacedrus papuana* | Gymnosperms | Pinales | Cupressaceae | Pappap | 1KP; 10.1186/2047-217X-3-17 | 1 | 0 | 0 | 0 | 0 | 0 | 1 | 0 | OVIJ | journals.rbge.org.uk/index.php/rbgesit | . | . |
| *Parahemionitis cordata* | Leptosporangiate | Polypodiales | Pteridaceae | Parcor | 1KP; 10.1186/2047-217X-3-17 | 1 | 0 | 0 | 0 | 0 | 0 | 1 | 0 | ZXJO | 10.1590/0102-33062016abb0074 | . | . |
| *Paraphymatoceros hallii* | Hornworts | Notothyladales | Notothyladaceae | Parhal | 1KP; 10.1186/2047-217X-3-17 | 0 | 0 | 0 | 0 | 0 | 0 | 0 | 0 | FAJB | 10.1098/rspb.2013.0207 | . | . |
| *Parasitaxus usta* | Gymnosperms | Pinales | Podocarpaceae | Parust | 1KP; 10.1186/2047-217X-3-17 | ND | 0 | 0 | 0 | 0 | 0 | ND | 0 | JZVE | 10.1111/j.1365-3040.2005.01378.x | . | Probably mycoparasite |
| *Parasponia andersonii* | Angiosperms | Rosales | Cannabaceae | Parand | 10.1073/pnas.1721395115 | 1 | 1 | 0 | 0 | 0 | 0 | 1 | 0 | NA | . | . | . |
| *Pellia cf. epiphylla* | Liverworts | Pelliales | Pelliaceae | Pelepi | 1KP; 10.1186/2047-217X-3-17 | 1 | 0 | 0 | 0 | 0 | 0 | 1 | 0 | PIUF | 10.3732/ajb.94.11.1756 | . | . |
| *Pellia neesiana* | Liverworts | Pelliales | Pelliaceae | Pelnee | 1KP; 10.1186/2047-217X-3-17 | 1 | 0 | 0 | 0 | 0 | 0 | 1 | 0 | JHFI | 10.3732/ajb.94.11.1756 | . | . |
| *Petunia axillaris* | Angiosperms | Solanales | Solanaceae | Petaxi | 10.1038/nplants.2016.74 | 1 | 0 | 0 | 0 | 0 | 0 | 1 | 0 | NA | . | . | . |
| *Phaeoceros carolinianus* | Hornworts | Notothyladales | Notothyladaceae | Phacar | 1KP; 10.1186/2047-217X-3-17 | 1 | 0 | 0 | 0 | 0 | 0 | 1 | 0 | RXRQ,WCZB | . | . | . |
| *Phalaenopsis equestris* | Angiosperms | Asparagales | Orchidaceae | Phaequ | 10.1038/nature23897 | 0 | 0 | 1 | 0 | 0 | 0 | 1 | 0 | NA | . | . | . |
| *Phaseolus vulgaris* | Angiosperms | Fabales | Fabaceae | Phavul | 10.1038/ng.3008 | 1 | 1 | 0 | 0 | 0 | 0 | 1 | 0 | NA | 10.1046/j.0028-646x.2001.00187.x | . | . |
| *Philonotis fontana* | Mosses | Bartramiales | Bartramiaceae | Phifon | 1KP; 10.1186/2047-217X-3-17 | 0 | 0 | 0 | 0 | 0 | 0 | 0 | 0 | ORKS | 10.1098/rstb.2000.0617 | . | . |
| *Phlebodium pseudoaureum* | Leptosporangiate | Polypodiales | Polypodiaceae | Phlpse | 1KP; 10.1186/2047-217X-3-17 | ND | 0 | 0 | 0 | 0 | 0 | ND | 0 | ZQYU | . | . | . |
| *Phyllocladus hypophyllus* | Gymnosperms | Pinales | Podocarpaceae | Phyhyp | 1KP; 10.1186/2047-217X-3-17 | 1 | 0 | 0 | 0 | 0 | 0 | 1 | 0 | JRNA | www.mycorrhizas.info/ | . | . |
| *Phylloglossum drummondii* | Lycophytes | Lycopodiales | Lycopodiaceae | Phydru | 1KP; 10.1186/2047-217X-3-17 | 1 | 0 | 0 | 0 | 0 | 0 | 1 | 0 | ZZEI | 10.1111/j.1469-8137.2007.02276.x | . | . |
| *Phymatosorus grossus* | Leptosporangiate | Polypodiales | Polypodiaceae | Phygro | 1KP; 10.1186/2047-217X-3-17 | 0 | 0 | 0 | 0 | 0 | 0 | 0 | 0 | ORJE | Jayaprakash, S. B., & Nagarajan, N. Stuc | . | . |
| *Physcomitrella patens* | Mosses | Funariales | Funariaceae | Phypat | 10.1111/tpj.13801 | 0 | 0 | 0 | 0 | 0 | 0 | 0 | 0 | NA | 10.1098/rstb.2000.0617 | . | . |
| *Physcomitrium sp.* | Mosses | Funariales | Physcomitrium | Physsp. | 1KP; 10.1186/2047-217X-3-17 | 0 | 0 | 0 | 0 | 0 | 0 | 0 | 0 | YEPO | 10.1098/rstb.2000.0617 | . | . |
| *Picea abies* | Gymnosperms | Pinales | Pinaceae | Picabi | 10.1038/nature12211 | 0 | 0 | 0 | 0 | 0 | 1 | 0 | 1 | NA | 10.1080/00275514.1993.12026249 | . | . |
| *Picea engelmannii* | Gymnosperms | Pinales | Pinaceae | Piceng | 1KP; 10.1186/2047-217X-3-17 | 0 | 0 | 0 | 0 | 0 | 1 | 0 | 1 | AWQB | 10.1139/b97-896 | . | . |
| *Picea glauca* | Gymnosperms | Pinales | Pinaceae | Picgla | 10.1093/bioinformatics/btt178 | 0 | 0 | 0 | 0 | 0 | 1 | 0 | 1 | NA | 10.1023/A:1014435407735 | . | . |
| *Picea sitchensis* | Gymnosperms | Pinales | Pinaceae | Picsit | Unpublished - gymno-plaza | 0 | 0 | 0 | 0 | 0 | 1 | 0 | 1 | NA | Harley, J. L., & Harley, E. L. (1987). A ch | . | . |
| *Pilgerodendron uviferum* | Gymnosperms | Pinales | Cupressaceae | Piluvi | 1KP; 10.1186/2047-217X-3-17 | 1 | 0 | 0 | 0 | 0 | 0 | 1 | 0 | ETCJ | www.mycorrhizas.info/ | . | . |
| *Pilularia globulifera* | Leptosporangiate | Salviniales | Marsileaceae | Pilglo | 1KP; 10.1186/2047-217X-3-17 | 0 | 0 | 0 | 0 | 0 | 0 | 0 | 0 | KIIX | 10.1016/0304-3770(85)90052-X | . | . |
| *Pinus jeffreyi* | Gymnosperms | Pinales | Pinaceae | Pinjef | 1KP; 10.1186/2047-217X-3-17 | 0 | 0 | 0 | 0 | 0 | 1 | 0 | 1 | MFTM | www.mycorrhizas.info/ | . | . |
| *Pinus parviflora* | Gymnosperms | Pinales | Pinaceae | Pinpar | 1KP; 10.1186/2047-217X-3-17 | 0 | 0 | 0 | 0 | 0 | 1 | 0 | 1 | IIOL | www.mycorrhizas.info/ | . | . |
| *Pinus pinaster* | Gymnosperms | Pinales | Pinaceae | Pinpin | Unpublished - gymno-plaza | 0 | 0 | 0 | 0 | 0 | 1 | 0 | 1 | NA | Harley, J. L., & Harley, E. L. (1987). A ch | . | . |
| *Pinus ponderosa* | Gymnosperms | Pinales | Pinaceae | Pinpon | 1KP; 10.1186/2047-217X-3-17 | 0 | 0 | 0 | 0 | 0 | 1 | 0 | 1 | JBND | www.mycorrhizas.info/ | . | . |
| *Pinus radiata* | Gymnosperms | Pinales | Pinaceae | Pinrad | 1KP; 10.1186/2047-217X-3-17 | 0 | 0 | 0 | 0 | 0 | 1 | 0 | 1 | DZQM | www.mycorrhizas.info/ | . | . |
| *Pinus sylvestris* | Gymnosperms | Pinales | Pinaceae | Pinsyl | Unpublished - gymno-plaza | 0 | 0 | 0 | 0 | 0 | 1 | 0 | 1 | NA | Harley, J. L., & Harley, E. L. (1987). A ch | . | . |
| *Pinus taeda* | Gymnosperms | Pinales | Pinaceae | Pintae | 10.1534/genetics.113.159715 | 0 | 0 | 0 | 0 | 0 | 1 | 0 | 1 | NA | 10.1007/BF00213445 | . | . |
| *Pityrogramma trifoliata* | Leptosporangiate | Polypodiales | Pteridaceae | Pittri | 1KP; 10.1186/2047-217X-3-17 | 1 | 0 | 0 | 0 | 0 | 0 | 1 | 0 | UJTT | 10.1016/j.chemosphere.2008.03.040 | . | At the genus level |
| *Plagiogyria japonica* | Leptosporangiate | Cyatheales | Plagiogyriaceae | Plajap | 1KP; 10.1186/2047-217X-3-17 | 1 | 0 | 0 | 0 | 0 | 0 | 1 | 0 | UWOD | 10.1007/s00572-015-0648-1 | . | . |
| *Plagiomnium insigne* | Mosses | Bryales | Mniaceae | Plains | 1KP; 10.1186/2047-217X-3-17 | 0 | 0 | 0 | 0 | 0 | 0 | 0 | 0 | BGXB | 10.1098/rstb.2000.0617 | . | . |
| *Platycladus orientalis* | Gymnosperms | Pinales | Cupressaceae | Plaori | 1KP; 10.1186/2047-217X-3-17 | 1 | 0 | 0 | 0 | 0 | 0 | 1 | 0 | BUWV | 10.6018/analesbio.37.8 | . | . |
| *Pleopeltis polypodioides* | Leptosporangiate | Polypodiales | Polypodiaceae | Plepol | 1KP; 10.1186/2047-217X-3-17 | 0 | 0 | 0 | 0 | 0 | 0 | 0 | 0 | UJWU | 10.15517/rbt.v65i3.29443 | . | . |
| *Podocarpus coriaceus* | Gymnosperms | Pinales | Podocarpaceae | Podcor | 1KP; 10.1186/2047-217X-3-17 | 1 | 0 | 0 | 0 | 0 | 0 | 1 | 0 | SCEB | 10.6018/analesbio.37.8 | . | . |
| *Podocarpus rubens* | Gymnosperms | Pinales | Podocarpaceae | Podrub | 1KP; 10.1186/2047-217X-3-17 | 1 | 0 | 0 | 0 | 0 | 0 | 1 | 0 | XLGK | 10.6018/analesbio.37.8 | . | . |
| *Polypodium amorphum* | Leptosporangiate | Polypodiales | Polypodiaceae | Polamo | 1KP; 10.1186/2047-217X-3-17 | ND | 0 | 0 | 0 | 0 | 0 | ND | 0 | YLJA | . | . | . |
| *Polypodium glycyrrhiza* | Leptosporangiate | Polypodiales | Polypodiaceae | Polgly | 1KP; 10.1186/2047-217X-3-17 | 1 | 0 | 0 | 0 | 0 | 0 | 1 | 0 | CJNT | 10.1139/b88-263 | . | . |
| *Polypodium hesperium* | Leptosporangiate | Polypodiales | Polypodiaceae | Polhes | 1KP; 10.1186/2047-217X-3-17 | ND | 0 | 0 | 0 | 0 | 0 | ND | 0 | GYFU,IXLH,ZRAV | . | . | . |
| *Polystichum acrostichoides* | Leptosporangiate | Polypodiales | Dryopteridaceae | Polacr | 1KP; 10.1186/2047-217X-3-17 | 1 | 0 | 0 | 0 | 0 | 0 | 1 | 0 | FQGQ | 10.2307/3792863 | . | . |
| *Polytrichum commune* | Mosses | Polytrichales | Polytrichaceae | Polcom | 1KP; 10.1186/2047-217X-3-17 | 0 | 0 | 0 | 0 | 0 | 0 | 0 | 0 | SZYG | 10.1098/rstb.2000.0617 | . | . |
| *Populus trichocarpa* | Angiosperms | Malpighiales | Salicaceae | Poptri | 10.1126/science.1128691 | 1 | 0 | 0 | 0 | 0 | 1 | 1 | 1 | NA | . | 10.1007/s11104-008-9877-9 | . |
| *Porella navicularis* | Liverworts | Porellales | Porellaceae | Pornav | 1KP; 10.1186/2047-217X-3-17 | ND | 0 | 0 | 0 | 0 | 0 | ND | 0 | KRUQ | 10.11646/phytotaxa.9.1.13 | . | . |
| *Porella pinnata* | Liverworts | Porellales | Porellaceae | Porpin | 1KP; 10.1186/2047-217X-3-17 | ND | 0 | 0 | 0 | 0 | 0 | ND | 0 | UUHD | 10.11646/phytotaxa.9.1.13 | . | . |
| *Potentilla micrantha* | Angiosperms | Rosales | Rosaceae | Potmic | 10.1093/gigascience/giy010 | 1 | 0 | 0 | 0 | 0 | 0 | 1 | 0 | NA | . | . | . |
| *Prumnopitys andina* | Gymnosperms | Pinales | Podocarpaceae | Pruand | 1KP; 10.1186/2047-217X-3-17 | 0 | 0 | 0 | 0 | 0 | 0 | 0 | 0 | EGLZ | www.dendrology.org/publications/tree | . | . |
| *Prunus avium* | Angiosperms | Rosales | Rosaceae | Pruavi | 10.1093/dnares/dsx020 | 1 | 0 | 0 | 0 | 0 | 1 | 1 | 1 | NA | 10.1016/S0007-1536(88)80146-3 | 10.1007/s00572-005-0033-6 | . |
| *Prunus dulcis* | Angiosperms | Rosales | Rosaceae | Prudul | Unpublished - rosaceae.org | 1 | 0 | 0 | 0 | 0 | 0 | 1 | 0 | NA | . | . | . |
| *Prunus mume* | Angiosperms | Rosales | Rosaceae | Prumum | Unpublished - NCBI | 1 | 0 | 0 | 0 | 0 | 0 | 1 | 0 | NA | . | . | . |
| *Prunus persica* | Angiosperms | Rosales | Rosaceae | Pruper | 10.1038/ng.2586 | 1 | 0 | 0 | 0 | 0 | 0 | 1 | 0 | NA | . | . | . |
| *Pseudolarix amabilis* | Gymnosperms | Pinales | Pinaceae | Pseama | 1KP; 10.1186/2047-217X-3-17 | 0 | 0 | 0 | 0 | 0 | 1 | 0 | 1 | AQFM | www.mycorrhizas.info/ | . | . |
| *Pseudolycopodiella caroliniana* | Lycophytes | Lycopodiales | Lycopodiaceae | Psecar | 1KP; 10.1186/2047-217X-3-17 | ND | 0 | 0 | 0 | 0 | 0 | ND | ND | UPMJ | . | . | . |
| *Pseudotaxiphyllum elegans* | Mosses | Hypnales | Hypnaceae | Pseele | 1KP; 10.1186/2047-217X-3-17 | 0 | 0 | 0 | 0 | 0 | 0 | 0 | 0 | QKQO | 10.1098/rstb.2000.0617 | . | . |
| *Pseudotaxus chienii* | Gymnosperms | Pinales | Taxaceae | Psechi | 1KP; 10.1186/2047-217X-3-17 | 1 | 0 | 0 | 0 | 0 | 0 | 1 | 0 | YLPM | www.mycorrhizas.info/ | . | . |
| *Pseudotsuga menziesii* | Gymnosperms | Pinales | Pinaceae | Psemen | 10.1534/g3.117.300078 | 0 | 0 | 0 | 0 | 0 | 1 | 0 | 1 | NA | . | . | . |
| *Pseudotsuga wilsoniana* | Gymnosperms | Pinales | Pinaceae | Psewil | 1KP; 10.1186/2047-217X-3-17 | 0 | 0 | 0 | 0 | 0 | 1 | 0 | 1 | IOVS | 10.1007/s00572-005-0033-6 | . | . |
| *Psilotum nudum* | Eusporangiate | Psilotales | Psilotaceae | Psinud | 1KP; 10.1186/2047-217X-3-17 | 1 | 0 | 0 | 0 | 0 | 0 | 1 | 0 | QVMR | 10.1007/s13199-012-0185-z | . | . |
| *Pteris ensiformis* | Leptosporangiate | Polypodiales | Pteridaceae | Pteens | 1KP; 10.1186/2047-217X-3-17 | 1 | 0 | 0 | 0 | 0 | 0 | 1 | 0 | FLTD | . | . | . |
| *Pteris vittata* | Leptosporangiate | Polypodiales | Pteridaceae | Ptevit | 1KP; 10.1186/2047-217X-3-17 | 1 | 0 | 0 | 0 | 0 | 0 | 1 | 0 | POPJ | 10.1007/s13199-012-0185-z | . | . |
| *Ptilidium pulcherrimum* | Liverworts | Ptilidiales | Ptilidiaceae | Ptipul | 1KP; 10.1186/2047-217X-3-17 | ND | 0 | 0 | 0 | 0 | 0 | ND | 0 | HPXA | . | . | . |
| *Pyrus communis* | Angiosperms | Rosales | Rosaceae | Pyrcom | 10.1371/journal.pone.0092644 | 1 | 0 | 0 | 0 | 0 | 0 | 1 | 0 | NA | 10.1080/00221589.1991.11516211 | . | . |
| *Pyrus x bretschneideri* | Angiosperms | Rosales | Rosaceae | Pyrbre | Unpublished - NCBI | 1 | 0 | 0 | 0 | 0 | 0 | 1 | 0 | NA | . | . | . |
| *Quercus robur* | Angiosperms | Fagales | Fagaceae | Querob | 10.1111/1755-0998.12425 | 1 | 0 | 0 | 0 | 0 | 1 | 1 | 1 | NA | . | 10.1111/nph.12317 | . |
| *Racomitrium elongatum* | Mosses | Grimmiales | Grimmiaceae | Racelo | 1KP; 10.1186/2047-217X-3-17 | 0 | 0 | 0 | 0 | 0 | 0 | 0 | 0 | ABCD | 10.1098/rstb.2000.0617 | . | . |
| *Racomitrium varium* | Mosses | Grimmiales | Grimmiaceae | Racvar | 1KP; 10.1186/2047-217X-3-17 | 0 | 0 | 0 | 0 | 0 | 0 | 0 | 0 | RDOO | 10.1098/rstb.2000.0617 | . | . |
| *Radula lindenbergiana* | Liverworts | Porellales | Radulaceae | Radlin | 1KP; 10.1186/2047-217X-3-17 | ND | 0 | 0 | 0 | 0 | 0 | ND | 0 | BNCU | 10.11646/phytotaxa.9.1.13 | . | . |
| *Retrophyllum minus* | Gymnosperms | Pinales | Podocarpaceae | Retmin | 1KP; 10.1186/2047-217X-3-17 | 1 | 0 | 0 | 0 | 0 | 0 | 1 | 0 | VGSX | www.mycorrhizas.info/ | . | . |
| *Rhododendron fortunei* | Angiosperms | Ericales | Ericaceae | Rhofor | 10.3389/fpls.2016.01594 | 0 | 0 | 0 | 1 | 0 | 0 | 1 | 0 | NA | . | . | . |
| *Rhododendron scopulorum* | Angiosperms | Ericales | Ericaceae | Rhosco | 1KP; 10.1186/2047-217X-3-17 | 0 | 0 | 0 | 1 | 0 | 0 | 1 | 0 | PPPZ | . | . | . |
| *Rhynchostegium serrulatum* | Mosses | Hypnales | Brachytheciaceae | Rhyser | 1KP; 10.1186/2047-217X-3-17 | 0 | 0 | 0 | 0 | 0 | 0 | 0 | 0 | JADL | 10.1098/rstb.2000.0617 | . | . |
| *Ricciocarpos natans* | Liverworts | Marchantiales | Ricciaceae | Ricnat | 1KP; 10.1186/2047-217X-3-17 | 0 | 0 | 0 | 0 | 0 | 0 | 0 | 0 | WJLO | 10.3732/ajb.94.11.1756 | . | . |
| *Ricinus communis* | Angiosperms | Malpighiales | Euphorbiaceae | Riccom | 10.1038/nbt.1674 | 1 | 0 | 0 | 0 | 0 | 0 | 1 | 0 | NA | 10.1023/A:1004951530917 | . | . |
| *Rosa chinensis* | Angiosperms | Rosales | Rosaceae | Roschi | 10.1038/s41588-018-0110-3 | 1 | 0 | 0 | 0 | 0 | 0 | 1 | 0 | NA | . | . | . |
| *Rosulabryum cf. capillare* | Mosses | Bryales | Bryaceae | Roscap | 1KP; 10.1186/2047-217X-3-17 | 0 | 0 | 0 | 0 | 0 | 0 | 0 | 0 | XWHK | 10.1098/rstb.2000.0617 | . | . |
| *Rubus occidentalis* | Angiosperms | Rosales | Rosaceae | Rubocc | 10.1111/tpj.13215 | 1 | 0 | 0 | 0 | 0 | 0 | 1 | 0 | NA | . | . | . |
| *Salvinia cucullata* | Leptosporangiate | Salviniales | Salviniaceae | Salcuc | 10.1038/s41477-018-0188-9 | 0 | 0 | 0 | 0 | 0 | 0 | 0 | 0 | NA | . | . | . |
| *Sarcodes sanguinea* | Angiosperms | Ericales | Ericaceae | Sarsan | 1KP; 10.1186/2047-217X-3-17 | 0 | 0 | 0 | 1 | 0 | 0 | 1 | 0 | SERM | . | . | . |
| *Saxegothaea conspicua* | Gymnosperms | Pinales | Podocarpaceae | Saxcon | 1KP; 10.1186/2047-217X-3-17 | 1 | 0 | 0 | 0 | 0 | 0 | 1 | 0 | QCGM | ojs.openagrar.de/index.php/JABFQ/art | . | . |
| *Scapania nemorosa* | Liverworts | Jungermanniales | Scapaniaceae | Scanem | 1KP; 10.1186/2047-217X-3-17 | 0 | 0 | 0 | 1 | 0 | 0 | 1 | 0 | IRBN | . | . | Assumes Jungermanniales are not AM |
| *Sceptridium dissectum* | Eusporangiate | Ophioglossales | Ophioglossaceae | Scedis | 1KP; 10.1186/2047-217X-3-17 | 1 | 0 | 0 | 0 | 0 | 0 | 1 | 0 | EEAQ | www.mycorrhizas.info/ | . | . |
| *Schistochila sp.* | Liverworts | Jungermanniales | Schistochilaceae | Schisp. | 1KP; 10.1186/2047-217X-3-17 | 0 | 0 | 0 | 1 | 0 | 0 | 1 | 0 | LGOW | . | . | Assumes Jungermanniales are not AM |
| *Sciadopitys verticillata* | Gymnosperms | Pinales | Sciadopityaceae | Sciver | 1KP; 10.1186/2047-217X-3-17 | 1 | 0 | 0 | 0 | 0 | 0 | 1 | 0 | YFZK | 10.21273/HORTSCI.30.1.133 | . | . |
| *Scouleria aquatica* | Mosses | Scouleriales | Scouleriaceae | Scoaqu | 1KP; 10.1186/2047-217X-3-17 | 0 | 0 | 0 | 0 | 0 | 0 | 0 | 0 | BPSG | 10.1098/rstb.2000.0617 | . | . |
| *Selaginella acanthonota* | Lycophytes | Selaginellales | Selaginellaceae | Selaca | 1KP; 10.1186/2047-217X-3-17 | 1 | 0 | 0 | 0 | 0 | 0 | 1 | 0 | ZYCD | www.mycorrhizas.info/ | . | Assumes Selaginellas are AM+ Brundrett 2008 |
| *Selaginella apoda* | Lycophytes | Selaginellales | Selaginellaceae | Selapo | 1KP; 10.1186/2047-217X-3-17 | 1 | 0 | 0 | 0 | 0 | 0 | 1 | 0 | LGDQ | www.mycorrhizas.info/ | . | Assumes Selaginellas are AM+ Brundrett 2008 |
| *Selaginella kraussiana* | Lycophytes | Selaginellales | Selaginellaceae | Selkra | 1KP; 10.1186/2047-217X-3-17 | 1 | 0 | 0 | 0 | 0 | 0 | 1 | 0 | ZFGK | Harley, J. L., & Harley, E. L. (1987). A ch | . | . |
| *Selaginella lepidophylla* | Lycophytes | Selaginellales | Selaginellaceae | Sellep | 1KP; 10.1186/2047-217X-3-17 | 1 | 0 | 0 | 0 | 0 | 0 | 1 | 0 | ABIJ | www.mycorrhizas.info/ | . | Assumes Selaginellas are AM+ Brundrett 2008 |
| *Selaginella moellendorffii* | Lycophytes | Selaginellales | Selaginellaceae | Selmoe | 10.1126/science.1203810 | 1 | 0 | 0 | 0 | 0 | 0 | 1 | 0 | NA | 10.1007/s00572-003-0272-3 | . | . |
| *Selaginella selaginoides* | Lycophytes | Selaginellales | Selaginellaceae | Selsel | 1KP; 10.1186/2047-217X-3-17 | 1 | 0 | 0 | 0 | 0 | 0 | 1 | 0 | KUXM | Harley, J. L., & Harley, E. L. (1987). A ch | . | . |

| | | | | | | | | | | | | | | | | |
|---|---|---|---|---|---|---|---|---|---|---|---|---|---|---|---|---|
| *Selaginella stauntoniana* | Lycophytes | Selaginellales | Selaginellaceae | Selsta | 1KP; 10.1186/2047-217X-3-17 | 1 | 0 | 0 | 0 | 0 | 0 | 1 | 0 | ZZOL | www.mycorrhizas.info/ | . | Assumes Selaginellas are AM+ Brundrett 2008 |
| *Selaginella wallacei* | Lycophytes | Selaginellales | Selaginellaceae | Selwal | 1KP; 10.1186/2047-217X-3-17 | 1 | 0 | 0 | 0 | 0 | 0 | 1 | 0 | JKAA | www.mycorrhizas.info/ | . | Assumes Selaginellas are AM+ Brundrett 2008 |
| *Selaginella willdenowii* | Lycophytes | Selaginellales | Selaginellaceae | Selwil | 1KP; 10.1186/2047-217X-3-17 | 1 | 0 | 0 | 0 | 0 | 0 | 1 | 0 | KJYC | www.mycorrhizas.info/ | . | Assumes Selaginellas are AM+ Brundrett 2008 |
| *Sequoia sempervirens* | Gymnosperms | Pinales | Cupressaceae | Seqsem | 1KP; 10.1186/2047-217X-3-17 | 1 | 0 | 0 | 0 | 0 | 0 | 1 | 0 | HBGV | 10.2737/PSW-GTR-151 | . | . |
| *Sequoiadendron giganteum Glaucum* | Gymnosperms | Pinales | Cupressaceae | Seqgig | 1KP; 10.1186/2047-217X-3-17 | 1 | 0 | 0 | 0 | 0 | 0 | 1 | 0 | QFAE | 10.2737/PSW-GTR-151 | . | . |
| *Setaria italica* | Angiosperms | Poales | Poaceae | Setita | 10.1038/nbt.2196 | 1 | 0 | 0 | 0 | 0 | 0 | 1 | 0 | NA | . | . | . |
| *Solanum lycopersicum* | Angiosperms | Solanales | Solanaceae | Sollyc | 10.1038/nature11119 | 1 | 0 | 0 | 0 | 0 | 0 | 1 | 0 | NA | . | . | . |
| *Solanum pennellii* | Angiosperms | Solanales | Solanaceae | Solpen | 10.1038/ng.3046 | 1 | 0 | 0 | 0 | 0 | 0 | 1 | 0 | NA | . | . | . |
| *Sorghum bicolor* | Angiosperms | Poales | Poaceae | Sorbic | 10.1111/tpj.13781 | 1 | 0 | 0 | 0 | 0 | 0 | 1 | 0 | NA | 10.1111/j.1469-8137.2004.01095.x | . | . |
| *Sphaerocarpos texanus* | Liverworts | Sphaerocarpales | Sphaerocarpaceae | Sphtex | 1KP; 10.1186/2047-217X-3-17 | 0 | 0 | 0 | 0 | 0 | 0 | 0 | 0 | HERT | 10.3732/ajb.94.11.1756 | . | . |
| *Sphagnum fallax* | Mosses | Sphagnales | Sphagnaceae | Sphfal | Unpublished - Phytozome | 0 | 0 | 0 | 0 | 0 | 0 | 0 | 0 | NA | 10.1098/rstb.2000.0617 | . | . |
| *Sphagnum lescurii* | Mosses | Sphagnales | Sphagnaceae | Sphles | 1KP; 10.1186/2047-217X-3-17 | 0 | 0 | 0 | 0 | 0 | 0 | 0 | 0 | GOWD | 10.1098/rstb.2000.0617 | . | . |
| *Sphagnum palustre* | Mosses | Sphagnales | Sphagnaceae | Sphpal | 1KP; 10.1186/2047-217X-3-17 | 0 | 0 | 0 | 0 | 0 | 0 | 0 | 0 | RCBT | 10.1098/rstb.2000.0617 | . | . |
| *Sphagnum recurvum* | Mosses | Sphagnales | Sphagnaceae | Sphrec | 1KP; 10.1186/2047-217X-3-17 | 0 | 0 | 0 | 0 | 0 | 0 | 0 | 0 | UHLI | 10.1098/rstb.2000.0617 | . | . |
| *Spinacia oleracea* | Angiosperms | Caryophyllales | Amaranthaceae | Spiole | Unpublished - bvseq.molgen.mpg.de | 0 | 0 | 0 | 0 | 0 | 0 | 0 | 0 | NA | . | . | . |
| *Spirodela polyrhiza* | Angiosperms | Alismatales | Araceae | Spipol | 10.1038/ncomms4311 | 0 | 0 | 0 | 0 | 0 | 0 | 0 | 0 | NA | . | . | . |
| *Stangeria eriopus* | Gymnosperms | Cycadales | Zamiaceae | Staeri | 1KP; 10.1186/2047-217X-3-17 | 1 | 0 | 0 | 0 | 0 | 0 | 1 | 0 | KAWQ | www.mycorrhizas.info/ | . | Several species of the familly formed AMS |
| *Stereodon subimponens* | Mosses | Hypnales | Hypnaceae | Stesub | 1KP; 10.1186/2047-217X-3-17 | 0 | 0 | 0 | 0 | 0 | 0 | 0 | 0 | LNSF | 10.1098/rstb.2000.0617 | . | . |
| *Sticherus lobatus* | Leptosporangiate | Gleicheniales | Gleicheniaceae | Stilob | 1KP; 10.1186/2047-217X-3-17 | ND | 0 | 0 | 0 | 0 | 0 | ND | ND | XDVM | . | . | . |
| *Sundacarpus amarus* | Gymnosperms | Pinales | Podocarpaceae | Sunama | 1KP; 10.1186/2047-217X-3-17 | 1 | 0 | 0 | 0 | 0 | 0 | 1 | 0 | KLGF | www.mycorrhizas.info/ | . | . |
| *Syntrichia princeps* | Mosses | Pottiales | Pottiaceae | Synpri | 1KP; 10.1186/2047-217X-3-17 | 0 | 0 | 0 | 0 | 0 | 0 | 0 | 0 | GRKU | 10.1098/rstb.2000.0617 | . | . |
| *Taiwania cryptomerioides* | Gymnosperms | Pinales | Cupressaceae | Taicry | 1KP; 10.1186/2047-217X-3-17 | 1 | 0 | 0 | 0 | 0 | 0 | 1 | 0 | QSNJ | www.mycorrhizas.info/ | . | . |
| *Takakia lepidozioides* | Mosses | Takakiales | Takakiaceae | Taklep | 1KP; 10.1186/2047-217X-3-17 | 1 | 0 | 0 | 0 | 0 | 0 | 1 | 0 | SKQD | Boullard, B. (1988). Observations on th | . | . |
| *Tarenaya hassleriana* | Angiosperms | Brassicales | Cleomaceae | Tarhas | 10.1105/tpc.113.113480 | 0 | 0 | 0 | 0 | 0 | 0 | 0 | 0 | NA | . | . | . |
| *Taxodium distichum* | Gymnosperms | Pinales | Cupressaceae | Taxdis | 1KP; 10.1186/2047-217X-3-17 | 1 | 0 | 0 | 0 | 0 | 0 | 1 | 0 | FHST | 10.1007/s13157-010-0017-y | . | . |
| *Taxus baccata* | Gymnosperms | Pinales | Taxaceae | Taxbac | 1KP; 10.1186/2047-217X-3-17 | 1 | 0 | 0 | 0 | 0 | 0 | 1 | 0 | WWSS | 10.1139/b03-020 | . | . |
| *Taxus cuspidata* | Gymnosperms | Pinales | Taxaceae | Taxcus | 1KP; 10.1186/2047-217X-3-17 | 1 | 0 | 0 | 0 | 0 | 0 | 1 | 0 | ZYAX | www.mycorrhizas.info/ | . | . |
| *Tetraclinis sp.* | Gymnosperms | Pinales | Cupressaceae | Tetsp. | 1KP; 10.1186/2047-217X-3-17 | 1 | 0 | 0 | 0 | 0 | 0 | 1 | 0 | CGDN | hal.archives-ouvertes.fr/hal-00883979 | . | . |
| *Tetraphis pellucida* | Mosses | Tetraphidales | Tetraphidaceae | Tetpel | 1KP; 10.1186/2047-217X-3-17 | 0 | 0 | 0 | 0 | 0 | 0 | 0 | 0 | HVBQ | 10.1098/rstb.2000.0617 | . | . |
| *Thelypteris acuminata* | Leptosporangiate | Polypodiales | Thelypteridaceae | Theacu | 1KP; 10.1186/2047-217X-3-17 | 1 | 0 | 0 | 0 | 0 | 0 | 1 | 0 | MROH | 10.2307/1352570 | . | At the genus level |
| *Theobroma cacao* | Angiosperms | Malvales | Malvaceae | Thecac | 10.1186/gb-2013-14-6-r53 | 1 | 0 | 0 | 0 | 0 | 0 | 1 | 0 | NA | 10.1007/BF00335843 | . | . |
| *Thuidium delicatulum* | Mosses | Hypnales | Thuidiaceae | Thudel | 1KP; 10.1186/2047-217X-3-17 | 0 | 0 | 0 | 0 | 0 | 0 | 0 | 0 | EEMJ | 10.1098/rstb.2000.0617 | . | . |
| *Thuja plicata* | Gymnosperms | Pinales | Cupressaceae | Thupli | 1KP; 10.1186/2047-217X-3-17 | 1 | 0 | 0 | 0 | 0 | 0 | 1 | 0 | VFYZ | 10.1007/s00442-004-1777-y | . | . |
| *Thujopsis dolabrata* | Gymnosperms | Pinales | Cupressaceae | Thudol | 1KP; 10.1186/2047-217X-3-17 | 1 | 0 | 0 | 0 | 0 | 0 | 1 | 0 | NKIN | www.mycorrhizas.info/ | . | . |
| *Thyrsopteris elegans* | Leptosporangiate | Cyatheales | Thyrsopteridaceae | Thyele | 1KP; 10.1186/2047-217X-3-17 | 1 | 0 | 0 | 0 | 0 | 0 | 1 | 0 | EWXK | journals.rbge.org.uk/index.php/rbgesit | . | . |
| *Timmia austriaca* | Mosses | Timmiales | Timmiaceae | Timaus | 1KP; 10.1186/2047-217X-3-17 | 0 | 0 | 0 | 0 | 0 | 0 | 0 | 0 | ZQRI | 10.1098/rstb.2000.0617 | . | . |
| *Tmesipteris parva* | Eusporangiate | Psilotales | Psilotaceae | Tmepar | 1KP; 10.1186/2047-217X-3-17 | 1 | 0 | 0 | 0 | 0 | 0 | 1 | 0 | ALVQ | 10.1139/b05-102 | . | . |
| *Torreya nucifera* | Gymnosperms | Pinales | Taxaceae | Tornuc | 1KP; 10.1186/2047-217X-3-17 | 1 | 0 | 0 | 0 | 0 | 0 | 1 | 0 | HQOM | 10.4489/KJM.2014.42.4.255 | . | . |
| *Torreya taxifolia* | Gymnosperms | Pinales | Taxaceae | Tortax | 1KP; 10.1186/2047-217X-3-17 | 1 | 0 | 0 | 0 | 0 | 0 | 1 | 0 | EFMS | www.mycorrhizas.info/ | . | . |
| *Trema orientalis* | Angiosperms | Rosales | Cannabaceae | Treori | 10.1073/pnas.1721395115 | 1 | 0 | 0 | 0 | 0 | 0 | 1 | 0 | NA | 10.1016/S0378-1127(00)00322-4 | . | . |
| *Treubia lacunosa* | Liverworts | Treubiales | Treubiaceae | Trelac | 1KP; 10.1186/2047-217X-3-17 | 1 | 0 | 0 | 0 | 0 | 0 | 1 | 0 | FITN | 10.3732/ajb.94.11.1756 | . | . |
| *Trifolium pratense* | Angiosperms | Fabales | Fabaceae | Tripra | 10.1038/srep17394 | 1 | 1 | 0 | 0 | 0 | 0 | 1 | 0 | NA | 10.1111/j.1526-100X.1998.00628.x | . | . |
| *Trifolium subterraneum* | Angiosperms | Fabales | Fabaceae | Trisub | Unpublished - NCBI | 1 | 1 | 0 | 0 | 0 | 0 | 1 | 0 | NA | 10.1111/j.1469-8137.2004.01095.x | . | . |
| *Tsuga heterophylla* | Gymnosperms | Pinales | Pinaceae | Tsuhet | 1KP; 10.1186/2047-217X-3-17 | 0 | 0 | 0 | 0 | 0 | 1 | 0 | 1 | GAMH | 10.1007/s005720050193 | . | . |
| *Utricularia gibba* | Angiosperms | Lamiales | Lentibulariaceae | Utrgib | 10.1073/pnas.1702072114 | 0 | 0 | 0 | 0 | 0 | 0 | 0 | 0 | NA | . | . | . |
| *Vigna angularis* | Angiosperms | Fabales | Fabaceae | Vigang | 10.1038/srep080669 | 1 | 1 | 0 | 0 | 0 | 0 | 1 | 0 | NA | . | . | . |
| *Vigna radiata* | Angiosperms | Fabales | Fabaceae | Vigrad | 10.1038/ncomms6443 | 1 | 1 | 0 | 0 | 0 | 0 | 1 | 0 | NA | www.jstor.org/stable/24090234 | . | . |
| *Vigna unguiculata* | Angiosperms | Fabales | Fabaceae | Vigung | Unpublished - Phytozome | 1 | 1 | 0 | 0 | 0 | 0 | 1 | 0 | NA | 10.1007/s00572-003-0277-y | . | . |
| *Vittaria appalachiana* | Leptosporangiate | Polypodiales | Pteridaceae | Vitapp | 1KP; 10.1186/2047-217X-3-17 | 1 | 0 | 0 | 0 | 0 | 0 | 1 | 0 | NDUV | www.jstor.org/stable/43185854 | . | At the genus level |
| *Vittaria lineata* | Leptosporangiate | Polypodiales | Pteridaceae | Vitlin | 1KP; 10.1186/2047-217X-3-17 | 1 | 0 | 0 | 0 | 0 | 0 | 1 | 0 | SKYV | . | . | . |
| *Welwitschia mirabilis* | Gymnosperms | Gnetales | Welwitschiaceae | Welmir | 1KP; 10.1186/2047-217X-3-17 | 1 | 0 | 0 | 0 | 0 | 0 | 1 | 0 | TOXE | 10.1007/BF00213462 | . | . |
| *Widdringtonia cedarbergensis* | Gymnosperms | Pinales | Cupressaceae | Widced | 1KP; 10.1186/2047-217X-3-17 | 1 | 0 | 0 | 0 | 0 | 0 | 1 | 0 | AUDE | www.mycorrhizas.info/ | . | . |
| *Wollemia nobilis* | Gymnosperms | Pinales | Araucariaceae | Wolnob | 1KP; 10.1186/2047-217X-3-17 | 1 | 0 | 0 | 0 | 0 | 0 | 1 | 0 | RSCE | 10.1071/BT97064 | . | . |
| *Woodsia ilvensis* | Leptosporangiate | Polypodiales | Woodsiaceae | Wooilv | 1KP; 10.1186/2047-217X-3-17 | 0 | 0 | 0 | 0 | 0 | 0 | 0 | 0 | YQEC | Harley, J. L., & Harley, E. L. (1987). A ch | . | . |
| *Woodsia scopulina* | Leptosporangiate | Polypodiales | Woodsiaceae | Woosco | 1KP; 10.1186/2047-217X-3-17 | 0 | 0 | 0 | 0 | 0 | 0 | 0 | 0 | YJJY | Harley, J. L., & Harley, E. L. (1987). A ch | . | . |
| *Zea mays PH207* | Angiosperms | Poales | Poaceae | Zeamay | 10.1105/tpc.16.00353 | 1 | 0 | 0 | 0 | 0 | 0 | 1 | 0 | NA | . | . | . |
| *Ziziphus jujuba cv. Dongzao* | Angiosperms | Rosales | Rhamnaceae | Zizjuj | 10.1038/ncomms6315 | 1 | 0 | 0 | 0 | 0 | 0 | 1 | 0 | NA | . | . | . |
| *Zostera marina* | Angiosperms | Alismatales | Zosteraceae | Zosmar | 10.1038/nature16548 | 0 | 0 | 0 | 0 | 0 | 0 | 0 | 0 | NA | . | . | . |

**Supplementary Table 2**: List of investigated genes. For each gene, ML model and tree likelihood are indicated as well as gene code in both v4.0 and v5.0 *M. truncatula* genome

| Gene | Model | Tree log-likelihood | Gene code M. truncatula v4.0 | Gene code M. truncatula v5.0 |
|---|---|---|---|---|
| ABCB12 | TVMe+R5 | -121226.7577 | Medtr8g022270 | MtrunA17_Chr8g0344651 |
| ABCB20 | GTR+F+R5 | -116544.4636 | Medtr3g093430 | MtrunA17_Chr3g0128391/MtrunA17_Chr3g0128381 |
| CASTOR | GTR+F+R10 | -89611.3568 | Medtr7g117580 | MtrunA17_Chr7g0276651 |
| CCaMK | SYM+R6 | -69448.1399 | Medtr8g043970 | MtrunA17_Chr8g0355131 |
| CCD | GTR+F+R6 | -79123.7823 | Medtr3g110195 | MtrunA17_Chr3g0140121 |
| CYCLOPS | GTR+F+R5 | -66258.0377 | Medtr1g033360 | MtrunA17_Chr1g0161921 |
| CYTB561 | TVM+F+R10 | -249829.3446 | Medtr1g017910 | MtrunA17_Chr1g0152401 |
| DHY | TVMe+I+G4 | -40091.8735 | Medtr1g109110 | MtrunA17_Chr1g0208941 |
| DnaJ | GTR+F+R7 | -57745.1097 | Medtr1g069725 | MtrunA17_Chr1g0183641 |
| EXO70 | SYM+R10 | -322063.3316 | Medtr1g062970 | MtrunA17_Chr1g0180411 |
| FatM | GTR+F+ASC+R10 | -320885.3301 | Medtr3g099200 | MtrunA17_Chr3g0132211 |
| GRAS | SYM+R9 | -190053.664 | Medtr3g467150 | MtrunA17_Chr3g0107521 |
| HEP | GTR+F+R9 | -321719.7461 | Medtr4g104750 | MtrunA17_Chr4g0058371 |
| HYP | GTR+F+R6 | -71726.063 | Medtr2g091215 | MtrunA17_Chr2g0325571 |
| HYP2 | VT+F+G4 | -27293.192 | Medtr3g104900 | MtrunA17_Chr3g0136131 |
| HYP3 | TIM3+F+R6 | -52371.6785 | Medtr6g007690/Medtr7g116650 | MtrunA17_Chr6g0451071/MtrunA17_Chr7g0275121 |
| HYP4 | TVMe+R5 | -27177.02 | Medtr1g090320 | MtrunA17_Chr1g0196911 |
| KinF | GTR+F+R5 | -48493.1809 | Medtr5g019040 | MtrunA17_Chr5g0403371 |
| KinG1/KinG2 | GTR+F+R6 | -80985.2209 | Medtr1g112940 | MtrunA17_Chr1g0211361 |
| LIN | GTR+F+R7 | -304073.5361 | Medtr7g027190 | MtrunA17_Chr7g0224271 |
| NFP | SYM+R8 | -298839.1305 | Medtr5g020810/Medtr3g118160 | MtrunA17_Chr5g0404521/MtrunA17_Chr3g0145801 |
| pp2a | SYM+R5 | -58526.9094 | Medtr7g087500 | MtrunA17_Chr7g0254621 |
| RAD1 | TVMe+R5 | -65962.6345 | Medtr2g088700 | MtrunA17_Chr2g0322801 |
| RAM1 | GTR+F+I+G4 | -48497.8892 | Medtr0021s0370 | MtrunA17_Chr0c05g0492121 |
| RFCA/RFCB | GTR+F+R5 | -55802.4051 | Medtr6g027840 | MtrunA17_Chr6g0461061 |
| SEC | GTR+F+R9 | -244103.4951 | Medtr5g030920 | MtrunA17_Chr5g0411631 |
| STR1/STR2 | SYM+R7 | -139653.7967 | Medtr2g008520 | MtrunA17_Chr2g0278961 |
| SymRK | GTR+F+R5 | -76240.236 | Medtr5g026850 | MtrunA17_Chr5g0409341 |
| SYN | GTR+F+R7 | -117674.5928 | Medtr4g104020 | MtrunA17_Chr4g0057891 |
| TAU | GTR+F+R10 | -256967.4813 | Medtr8g107450/Medtr5g030910 | MtrunA17_Chr8g0393191/MtrunA17_Chr5g0411621 |
| VAPYRIN | GTR+F+R7 | -176119.4627 | Medtr4g097510 | MtrunA17_Chr4g0054341 |

**Supplementary Table 3:** Results of the branch and branch-site models analysis for signature of selection acting on *CCaMK* and *CYCLOPS* in the non-mycorrhizals mosses.

| | | | Branch models | | | | | Branch-site models | | |
|---|---|---|---|---|---|---|---|---|---|---|
| **Gene** | **Number of sequences** | **Number of positions** | $\omega_{BG}$ | $\omega_{FG}$ | **b_free *vs* M0** | **b_free *vs* b_neut** | **RELAX on mosses clade[1]** | **bsA *vs* M0** | **bsA *vs* bsA1** | **number of sites under positive selection** |
| *CCaMK (DMI3)* | 124 | 1092 | 0.097 | 0.297 | $\omega_{FG}$ and $\omega_{BG}$ are different *p* =2.95E-04*** | $\omega_{FG}$ is not neutral *p* =7.55E-03** | selective relaxation on mosses clade K=0.11*** | sites on FG are under relaxed selection *p* =5.55E-10*** | sites on FG are under positive selection *p* =2.07E-02* | 11 |
| *CYCLOPS (IPD3)* | 121 | 522 | 0.130 | 0.538 | $\omega_{FG}$ and $\omega_{BG}$ are different *p* =3.45E-02* | $\omega_{FG}$ is neutral *p* =0.59 | selective relaxation on mosses clade K=0.87* | sites on FG are under relaxed selection *p* =2.70E-05*** | sites on FG are under positive selection *p* =7.60E-04*** | 3 |

FG: foreground branch (branch between the ancestor of *Takakia lepidozioides* and the other mosses); BG: background branches (rest of the tree); mosses clade[1]: all branches of mosses without *T. lepidozioides*

Significance of LRT: *0.05>*p-val* >0.01, **0.01 >*p-val*> 0.001, ****p-val* <0.001, absence of symbols: not significant

**Supplementary Table 4**: List of Marchantia polymorpha accessions with GPS coordinates of the sampling sites.

| Population_id | Population_site_name | GPS_coordinates |
|---|---|---|
| Ber-C | Bernac-C | 43°10'03.3"N 0°06'31.2"E |
| Ber-D | Bernac-D | 43°10'03.3"N 0°06'31.2"E |
| Bid-A | Bida-A | 43°08'25.7"N 2°09'33.5"W |
| Bie-A | Biert-A | 42°53'53.9"N 1°18'53.9"E |
| Bot-X | Botanic-X | collected in a pot in the Toulouse botanical garden |
| Bul-B | Bulan-B | 43°02'23.3"N 0°16'38.5"E |
| Cas-A | Castillon-A | 42°55'14.8"N 1°01'57.5"E |
| Cas-C | Castillon-C | 42°55'14.8"N 1°01'57.5"E |
| Cas-D | Castillon-D | 42°55'15.3"N 1°02'05.4"E |
| Cas-E | Castillon-E | 42°55'14.9"N 1°02'01.9"E |
| Caz-A | Cazavet-A | Cazavet (Ariège, France) |
| Dur-A | Durban-sur-Arize-A | 43°01'18.5"N 1°21'03.3"E |
| Esc-A | Esconnets-A | 43°04'09.0"N 0°13'45.6"E |
| Gen-A | Genos-A | 42°48'40.9"N 0°24'13.7"E |
| Gen-B | Genos-B | 42°48'40.9"N 0°24'13.7"E |
| Hec-A | Heches-A | 43°00'58.8"N 0°22'19.2"E |
| Hec-C | Heches-C | 43°00'58.8"N 0°22'19.2"E |
| Lac-A | Lacourt-A | 42°56'37.5"N 1°10'28.3"E |
| Luc-B | Luchon-B | 42°47'08.6"N 0°35'37.2"E |
| Mns-A | Monségur-A | 42°52'13.0"N 1°49'59.6"E |
| Mou-A | Moulis-A | 42°57'36.4"N 1°05'18.8"E |
| Mur-A | Murianettes-A | 45°11'23.9"N 5°48'44.2"E |
| Nor-A | Norwich-A | 52°38'02.3"N 1°17'48.6"E |
| Nor-B | Norwich-B | 52°37'54.5"N 1°18'00.1"E |
| Nor-C | Norwich-C | 52°37'50.7"N 1°18'10.1"E |
| Nor-D | Norwich-D | 52°37'50.7"N 1°18'10.1"E |
| Pra-A | Prat-A | 43°01'44.2"N 1°01'05.5"E |
| Pra-B | Prat-B | 43°01'43.2"N 1°01'05.1"E |
| Pra-C | Prat-C | 43°01'43.1"N 1°01'03.3"E |
| Tou-A | Tou-A | 46°39'01.5"N 4°06'49.8"E |
| Tou-B | Tou-B | 46°39'01.5"N 4°06'49.8"E |
| Tou-C | Tou-C | 46°39'01.5"N 4°06'49.8"E |
| Tou-D | Tou-D | 46°39'01.5"N 4°06'49.8"E |
| Tou-E | Tou-E | 46°39'01.5"N 4°06'49.8"E |
| Voe-A | Voewood-A | 52°54'53.5"N 1°07'09.0"E |

**Supplementary Table 5:** CCaMK orthologs complement the *ccamK* root nodule organogenesis phenotype in *M. truncatula* roots. *ccamk M. truncatula* mutant roots were transformed with *pUb:CCAMK* from *M. truncatula* (Medtru), *M. pudica* (Mimpud), D. trinervis (Distri), *F. vesca* (Fraves), *H. vulgarum* (Horvul), *Z. mays* (Zeamay), *M. paleacea* (Marpal) or an empty vector (control) and scored 42 days post-inoculation (dpi) with *S. meliloti* . Numbers of plants showing at least one nodule are indicated out of the total number of plants transformed.

| **Constructs** | **Number of plants with nodule/ Total number of plants** | **% of plants showing nodules** |
|---|---|---|
| Control | 0/7 | 0 |
| *Medtru* | 10/12 | 83 |
| *Mimpud* | 15/18 | 83 |
| *Distri* | 12/17 | 86 |
| *Fraves* | 3/3 | 100 |
| *Horvul* | 17/18 | 94 |
| *Zeamay* | 12/12 | 100 |
| *Marpal* | 8/9 | 89 |

**Supplemental Table 6:** Sequences collected by tBLASTn against genome assemblies

| Medtru v5 | Gene name | Species | Scaffold name/Accession number | Predicted Protein sequences |
|---|---|---|---|---|
| Medtru_MtrunA17_Chr5g0409341 | ***CYCLOPS*** | *Mimosa pudica* | scaffold786_cov186 | MEGRGFSGLYKNTSEELFLKAVMETPIGMPAPTMEMLGFKAVSQSFRADSEELFKRWLTNGEASQAYDSILYHHNFFAFFKGYNSTGIGHTSRLSQRISTELANLSNPQHVGVISEEKINEKLYTQNNLLVNDASGDFNEPPIRDAADRDLQTSNLFLAKAWFLSDQRMTRSRSSELRRRYVEMQNAQTTQGIEPIHLVPHNGGADTLKQEVTNVNGFSYLSPCELPNHKATFMSPSNSSSSTFNAPQIMADADKISSCVSMLKGTLERKRQSNSNNIEKEAAEDSSNGIFVAQGDFAKTVFMEGQENWIYQKPINFQGTSNNGQVEDHRVRQTLEGSMNLEMDGFTNQTNPIYIGTASQEPSQSESSAAAPVVSSGLDAIEGPSISSQTPHENSWKQVGVNKGSENKVKGVREQIMDNLKDDRKKRSLERYGSVNSVVSEGKGDATKKRRVERSRKMAEAKERNSTPPVPSDMQAVLKRCEALEKEVRSLKLNLSFMNRKDSEQTKQIEELQKQNEDLADEKERLLEEIERIVSETGKI |
| Medtru_MtrunA17_Chr5g0409341 | ***CYCLOPS*** | *Casuarina glauca* | scaffold210 | MEDLYRNSSEEMFLKYFMESSNAMAAPSMEMLGFKNLSPNFRHTDSEELFRSWLTNGEASGAILSSLIGFITLNNNNTSSSMGPRTRQGSRRLSTELASLSSQQHVGIAQKKRSNDTLHPQSNALVDNIQGDLHQQSIRGSVEKGLHTTDLYLAKAWFHSSQPMTRSRSSELRRRYAAMQSAQTTLGMEVLHNASGHGANTLKQDFSNSNCFNDLSMSEIPNQWRTFMSPSNSSSSTLNTPQMAEVDKVSSVVSMLKGTLERKKLSNRIEKEAVVDSSNGIYHGHDFIVNSSFDQGQKNHLNDMPIFFQEASNVHVTDPGILHTVDESIDLELEGFVNPTNPVQLSRGSQEPSQSESSAAAPVLSSGFDACDGPSNSSQTLSFCESSRKQVGNSKSSGNGTRAKEIRERIIDTLKEDRKKGSLVRYGSVTSAVSVDKGDSTKKRRVERSRKMAEAKERNLTPAIPSDMQAVLKRCETLEKEVRSLKLNLSFMNRKDSEQTKQIEELQKQNEDLVDQKERLVEEIERVSSESGRMC |
| Medtru_MtrunA17_Chr5g0409341 | ***CYCLOPS*** | *Selaginella moellendorffii* | scaf4-2 | MMVRNTSEEMIIRSLIDGGSTATHGAAAAIAFENMGFMPHRASSEELFNSWIANSEGGLPPHTRPSRKMSSELAALLAQQQGMDHGLYLQSDHNAKSRRRYLEMQRQDLSSGAKPTEQSFLPWPDNLPQHDHNSSVSTVSTVVNMLKGSLEKKRVSKELQHHCQAEQHELHQQQNAVQDASMEASDVSTKKRRVERKRMMVEAKGRSSVPAMPSDMQAAIKRSDALEKEVRSLKLNLSFMNRKDSEQTKRIEELEKQNEELLREKSRLTEELDQLISSRSTW |
| Medtru_MtrunA17_Chr5g0409341 | ***CYCLOPS*** | *Selaginella moellendorffii* | scaf9 | MMVRNTSEEMIIRSLIDGGSTATHGAAAAIAFENMGFMPHRASSEELFNSWIANSEALSWFQTSQPMTRSRSTELRRRYLEMQRQDLSSGAKPTEQSFLPWPDNLPQHDHNSSVSTVSTVVNMLKGSLEKKRVSKELQHHCQAEQHELHQQQNAVQDASFMFRLANPSESSGGNDNNSCPATPSQIHLFHDQQAGRAISMEASDVSTKKRRVERKRMMVEAKGRSSVPAMPSDMQAAIKRSDALEKEVRSLKLNLSFMNRKDSEQTKRIEELEKQNEELLREKSRLTEELDQLISSRSTW |
| Medtru_MtrunA17_Chr8g0355131 | ***CCaMK*** | *Prunus persica* | Pp02 | MGQETRRLADEYEISEILGRGGFSVVRKGISRKSGSSDKNNVAIKTLKRPFGPSNPPPGRSGAGDHQKSFAAAFQARKQVSISNVLLTNEILVMRRIVENVSPHPNVIDLYDVYEDENGVHLVLELCSGGELFDRIVKQERYSEAGAAAVVRQIAQGLAALHKSNIVHRDLKPENCLFLNNTDDSSLKIMDFGLSSVEEFTDPVVGLFGSIDYVSPEALSQGQVTTKSDMWALGVILYILLSGYPPFIAQSNRQKQQMIMAGEFSFYEKTWKGISLSAKQLISDLLKVDPDKRPSAQELLDHPWVVGLSAKEDQMDAEIVSRLQSFNARRKLRAAAIASVWTSSIFLRTKKLKSLLGSHDLKPDEIQKLSLHFKKICVKGDNATLSEFEEVLKAMNMSSLIPLAPRIFDLFDNNRDGTVDMREILCGFSSLKNSQGDDALRLCFQMYDTDRSGCISKEEVASMLRALPDDCLPADITEPGKLDEIFDRMDANSDGQVTFDEFKAAMQRDSSLQDVVLSSLRPL |
| Medtru_MtrunA17_Chr5g0411631 | ***SYMRK*** | *Chamaecrista fasciculata* | scaffold70_cov204 | MMMELNIWKLRLIKFYVFCLIAFIRSASSAREGFVSIACCADSDYTDPKTSISYTSDYSFFSDRTSCQDINNHVGNERARVFSLDTGKRCYNLSTIKDQVYLLRGTFPSVESFSSSLDSSFDVTIGVTSLGAIEKPSQDLKIEGVFRATKNYTDFCLVKDQGSPYISQIELRPLDEVYLQGFTSSVLKLISRENLGSQEDDIRYPQDQSDRIWNSSSNPSYTSLQANVTISGHNADSTTPPLRVLQTAFTHPERLEFIHDGSDSANSEYRLFLYFLELNASVAEGQKVFDIYVNDEIKEPRFDILASGSNYRNASLNVSANGWLNITLVKSSGSAFGPLCNAYEILQVIPWMQETVQTDTEVILKVREQLLENNRGNQELESWSGDPCTLSPWQHVGCDRSSSGLLAITKLDLSSSNLQGPIPPSVTELTNLETLNLSYNSFTGNIPSFPSSSKLITIDLSHNDLTGSLLGSIPSLPQLKFLYFGCNSRMSKEIPANLSSSLSTDNGRCKAGDPGISQKQIIIGTVTCGSLLIILAVGVLLVYRYKEKLIPWRTPRVKGFPTTTNLIFSLPSSKDDFFTKSLSFRTESVQTFSLEYIEAATENYKILIGEGGFGPVYRGTLGDGQEVAVKVRSATSTQGTREFDNEVNLLSAIRHENLVPLLGYCCEDDQQILVYPFMSNGSLQDRLYGEPAKRKVLDWPTRLSIALGAARGLSYLHTFSGRSVIHRDVKSSNILLDHSMCAKVADFGFSKYAPQEGDSGASLELRGTAGYMDPEYYTTQQLSAKTDVFSFGVVLLEMVSGREPLNIHRPRNEWSLVEWAKPYIRASKIDEIVDPGIKGGYHAEAMWRVVETALSCIEQLSANRPSMAAIVRELEDALIIENNASEYMKSIDSLGGSNRYSIVIDKRVLPPTPSTTESTISTKPLSLPQPR |
| Medtru_MtrunA17_Chr5g0411631 | ***SYMRK*** | *Morus notabilis* | ENA\|ATGF01019099\|ATGF01019099.1 | MMEGLLVNNWVSRTCRCLILCLFILAQSTCAQKDFVSIACCRESNYTDPNTTISWDTDYRWFPDRKNCRKISKASTNHTGDDEARIFDIDYSWKRCYSLDTTRGEEYLIRGTFLYYENILLYETSFSVSVGVTNISQVNSAEDFEVEGIFRATKESIDFCLVKTEGDPYISRLVLRPLKGLEYLQDFSSSSVLKVVRRINLGAKDGDIIKYPTDPSDRIWKPLTSSGTANTTIQGSNVKISNYTGKVRAPLQVLQTALSDQNRLEILHNDLDKGNHQYRLTLYFLELNTTVYSGQRVFDIYVNNQRMKSSFDIKGGGSNYKEVRHDVRANGFLNLTLVKASGSSLGPICNAYEILQVFPWAQETSPSDVNGILEVKRELLACNEENKVLESWTGDPCFPFSWEGLSCESINGSTVIAKLSLPSMNLKGPIPPSITKLTNLITLYVDRRLFFLFGLSMLSLLDLRNNDLRGPLPGSLAMMQHLEILYVGCNPVAKEDPSSFNSSKLDTSSGTCNPGPASSKRRIIIGSVACGSLLLTLAAGAFVLCFYRQKLMPQGIFNGKGFKGPPMSKKNGGLIFSIPIVEEFVVKSITIQTFTLANIEAATQNYNTLIGEGGFGSVYRGTLPDGQEVAVKARSATSTQGTREFENELNLLSTIRHENLVPLLGYCSENEQQILVYPFMSNGSLQDRLYGEAAKRKRLDWPTRLSIALGAARGLTHLHTYAGRCVIHRDVKSSNILLDQSMTAKVADFGFSKYAPQEGDSGASLEVRGTAGYLDPEYYTTHHLSDKSDVFSFGVVLLEIISGREPLNIRRPRNEWSLVEWAKQHARETNIDEIVDPSIKGGYHAEAMWRVVEMALLCIEPFSVYRPSMADIVRELEDSLIIENNASEYMRSIESSGSIVLDRKIVIPSSTPTEASPVLTQPLPPPEPR |
| Medtru_MtrunA17_Chr8g0356441 | ***RAD1*** | *Carica papaya* | supercontig_87:828041- | MSSPVVQPEVPDLVCELDNVQGIVDAFSSVRWKRHQDALVELSEHGIVLIVEESGCLQAKVYLQRELFIRYEYGAQGRPRFGVSLGLFVDCLNTFSVPGHVSTIEIKYPGPDMQLLLKSVDSLDACIYAEIRTRIPDTISWDYNFEPAGTTPLTFTVKSAALKEAIDDLEWPGSSILIILQPVPPCVTFKGEGHGDLHIDFMYYANTDLLVAFHCDHQVSYRYKYKYLKATTSNIPSSVIKENRGSKLTIGRGGMLKVQHLVSVARPFSHTHIDSSGYNQPSRIAYIEFYVKPEEREDAADGS |
| Medtru_MtrunA17_Chr8g0356441 | ***RAD1*** | *Arachis ipaensis* | araip.K30076.gnm1.Araip.B08 | MSASGTEEERPELVCQLEHIQGLVDALSAVRWKRHQDAVLELSEHGIVLVVQESGCLQAKLYLKRELFVRYDYNAQVRPRFGLSLGIFVECLNAFSVPAHSTPIQIQYPGPDMQLLLKSVDSMDASICAEIRTRIPETVAWDYNFEPAGTNPLTFTVKSAALKEAIEDLEWPGSSIQVILEPDPPSVTLRAEGHGDLQIDFLYYANSDLLVAFQCDNRVSVKYKYKFLKATTSNMPSSVIKENRGSKLSIGRGGMLKVQHLVSIAKPLASHSYVDSIGYQQPGRVAHIEFFVKPEESDGSTND |
| Medtru_MtrunA17_Chr8g0356441 | ***RAD1*** | *Arachis duranensis* | aradu.V14167.gnm1.Aradu.A07 | MSASGTEEERPELVCQLEHVQGLVDALSAVRWKRHQDAVLELSEHGIVLVVQESGCLQAKLYLKRELFVRYDYNAQVRPRFGLSLGIFVECLNAFSVPAHSTPIQIQYPGPDMQLLLKSVDSMDASICAEIRTRIPETIAWDYNFEPAGTNPLTFTVKSAALKEAIEDLEWPGSSIHVILEPDPPSVTLRAEGHGDLQIDFLYYANSDLLVAFQCDHRVSVKYKYKFLKATTSNMPSSVIKENRGSKLSIGRGGMLKVQHLVSIAKPLASHSYVDSIGYQQPSRVAHIEFFVKPEESDDSTND |
| Medtru_MtrunA17_Chr8g0356441 | ***RAD1*** | *Arachis hypogaea* | arahy.Tifrunner.gnm1.Arahy.07 | MSASGTEEERPELVCQLEHVQGLVDALSAVRWKRHQDAVLELSEHGIVLVVQESGCLQAKLYLKRELFVRYDYNAQVRPRFGLSLGIFVECLNAFSVPAHSTPIQIQYPGPDMQLLLKSVDSMDASICAEIRTRIPETIAWDYNFEPAGTNPLTFTVKSAALKEAIEDLEWPGSSIQVILEPDPPSVTLRAEGHGDLQIDFLYYANSDLLVAFQCDHRVSVKYKYKFLKATTSNMPSSVIKENRGSKLSIGRGGMLKVQHLVSIAKPLASHSYVDSIGYQQPSRVAHIEFFVKPEESDDSTND |
| Medtru_MtrunA17_Chr8g0356441 | ***RAD1*** | *Castanea mollissima* | scaffold03205 | MSSEGPDLVCEVDNVQGIVDALTAVRWKRQQDAVVELSEHGVVLIVEESSCLQAKVYLQRELFIKYEYGGAQGRPRFGVSLGLLVNCLNTFSVPGSGRSSSTIEIRYPGPDMQLLLKSVDSLDACIYAEIRTRIPETISWDYNFEPAGTTPISFTVKSAALKEAIDDLEWPGSSVQIILQPDPPSVTFKGEGHGDLQIDFMYYVNTDLLIAFSCDHQVSYRYKYKFLQATTSNIPSSVIRDNRGSKLTIGRGGMLKVQHLVSVARPSIPHPHVDSAGYQQPSRIAYIEFFVKPEEDESTIND |
| Medtru_MtrunA17_Chr8g0393191 | ***STR1*** | *Quercus robur* | gnl\|Qrob_v2_Genes_v2.2_CDS_prot_20151202\|Qrob_P0386870.2 808 | MSHTNSDRRDTVIDIGKLVNFTGGLEFSSLTYTVTKKKKDVEGEWFKKEVDLLNRITGYAPKGCITAVMGPSGAGKSTLLDGLAGRIASGSLKGRVSLDGKEMSPSLIKRTSAYIMQDDRLFPMLTVYETLMFAADFRLGPISRAEKQERVDKLIEQLGLSSSRNTYIGDQGIRGVSGGERRRVTIGVDIIHGPSLLFLDEPTSGLDSTRALSVIEKVHHIARSGSTVILTIHQPSSRIQLLLDHLIILARGQLMFQGSPKDVTHHVSRMGRRVENSIEHLIDVIQEYDQSEHGVEAVAEFARTRLKAPPLAGMDMSVSSTLPPTPTPPRRSRQAEGQESADHHKGQGKRLQLQNNSLVFDSFDHSVRSPYNNTSRSWSASDRGVMQALRLTPSQANKKGPNSIRDMSVSSTLPPTPTPPRRSRQAEGQESADHHKGQGKRLHVRSPYNNTSRSWSASDRGVMQALRLKPSQANKKGPNSISLSPGHYAYSSEILPSTPTPHSSDYPVNEDDYRTPGGADVAMNSTANNNHLGPKFANSFLSEIMILMLRNFKNIRRTPELFLSRLMVLTIMGFMMATMFVKTPKSLQGITDLTSFFIFTVCLFFFSSNDAVPAFIQERFIFIRETSHNAYRASSYTIAGVVTYLPFLAIQAGVYAGIVYFALKLKGLFIYFLIVLYVSLLTTNSFVVFVSSVVPNYILGYAAVIAFTALFFLFCGYFLSGHDIPSYWQWMNKISTMTYPYEGLLINEFQRTDIYAKDPLGNNVSGIDVLDSLKIHGSGGKKWEKVFIMLGWAVIYRILFYIILRFGG |
| Medtru_MtrunA17_Chr8g0393191 | ***STR1*** | *Cucurbita moschata* | CmoCh07G005930.1 | MAHGGGGQRRDTTIDIGRTAKFNGGLEFYDLTYTVIKDKEHEGKLVKQEVDLLHRISGYSPKGSVTAVLGPSGAGKSTFLDGLAGRIASGSLKGRVSLDGMEMSPGLIKRTSAYIMQDDRLFPKLTVYETLMFAADFRLGSIPSSEKIHRVENLIEQLGLSSARNTYIGDEGTRGVSGGERRRVSIGVDIIHGPSLLFLDEPTSGLDSTSAYSVIEKVHNIARTGSTVILTIHQPSSRILSFFDHLIVLARGQLMFQGPPKDVNHHLSQMKRKVPQGESPIEYLMDVIRAYDQSEFGVEALAEFARTGTKPPHLTDEEISLSTIQASLTSSFQSGVDKTSNIITGKRLHLQTNSRALNDYGHSLRSPYNTSRSWSANNSVVMQALRLPQRQQGAKLHNQMSSSASYAYSSDVLHGTPTPHSSDYTVNENDYLTSNIGSKSAPTLHNNLGKKISNSFFSETWILMRRNFNIISRTPELFLSRMMVLTVMGFMMATMFMKPKENTQGITDRLSFFIFTVCLFFFSSNDAVPAFIQERFIFIRETSHNAYRASSYTIAGLLTHLPFLALQALVYAGIVWFALKLRGSFIYFLIVLYMSLLTSNSFVVFISSVVPNYILGYAAVIAFTALFFLFCGYFLNSHDLPPYWKWMNKISTMTYPYEGLLMNEYQTSIFFGNQSNGTVITGINILESLHIKTDSTKKWENVLVMLGWAVLYRILFYLILRFASKNQRT |
| Medtru_MtrunA17_Chr8g0393191 | ***STR1*** | *Discaria trinervis* | scaffold896\|size107627 | MAKLGRRETNKSLESLLDMDKSAAAKNALPLQRRTLVPGQGLEFHNLSYSVIKKQKKDGVWIKKQAYLLNDISGQAARGEVLAIMGPSGAGKSTFLDALAGRIAQGSLEGSVRINGKPVSASYMKMVSSYVMQDDQLFPMLTVFETFMFAAEVRLPPSISRAEKKKRVYELLDQLGLQSAMHTYIGDEGRRGVSGGERRRVSIGIDIIHKPSLLFLDEPTSGLDSTSAYSVVEKVKDIAQGGSMVLMTIHQPSYRIQMLLDRIIVLGRGRLVYAGSPLGLSSHLDGFERPVPEGENSLEYLLDVIKEYDESTVGLDPLVLYQRDGIKPDQVARTPIPKPPKTPRTPHQKTPTAKHMITLHSQGFSSGNMTPVPESGHLDYRYEDEDDDDDFDNSLERKTVQTPINIQSGVYNPRLASHFYKDFSMWVYNGVRGTPRRQPSWTPARTPGKTPNFSGGRSIVSSQFPTPQINPSVTKTPIVFSASMDSYTASYEEFDLEEEEILDEPEHHGPKFANPWLREVMVLSWRTTLNVIRTPELFLSREVVLTVMALILSSLFKDLSHETFKDINRLLNFYIFAVCLVFFSSNDAVPTFIQERFIFIRETSHNAYRASSYVISSLIVYLPFFAIQGFTFSGITKLLLHLKSNLLNFWLILYASLITTNAYVMLVSALVPSYITGYAVVIATTALFFLTCGFFLKQFQIPKYWRWLHYISAIKYPFEALLVNEFKGNKCYNGRPEDLSPGPLGDVKYSKLHNNTIGFRPDCLLIGDDVLFSMDVTKPDIWYDIGILLAWGVLYRLLFYVVLRFYSKNVRK |
| Medtru_MtrunA17_Chr8g0393191 | ***STR1*** | *Ginkgo biloba* | scaffold816 | MGTVNRYNAEETNGLEFDDLSYTVRKKQKVHDSRISKEVDLLHQISGQALRGEITAVMGPSGAGKSTFLDGLAGRIAQGSLRGSVSVDGRQVSTSFMKMISAYVLQDDQLFPMLTVYETLLFAAEVRLPPFISKMEKLKRVDDLLEQLGLKSVAHTYIGDEGRRGVSGGERRRVSIGVDIIHGPPLLFLDEPTSGLDSSSAYSVVEKLKDIAKNGSIIIMTIHQPSFRIQMLFNRLIVLGRGQLIYMGSPTYLPEHLSGFGKQVPEGENSIEYLLDVIKEYDESTVGLEPLVAYQRDGKRPSKAAQTPQPRTPVARSMSATPQRDLKTIEEDNEDFDNSLERKTPMNNFGYPRLASHFYKDFSLWIYNEITGTPSRPPSWTVARTPGRTPGLTPGRTPGRTPGRTPGRTPGRMPDQSYYPYYWRTPGRTPNFQNPHQSPVSHIAPPGVEMFSSVDRVVRFASQSEFEGDEDELELDEEFYGGSKFANPWIREVIVLSWRNFLNVIRTPELFLSREVVLIVMGLTLATLFQNLNKDGMTNVNQILSFYIFAICLLFFSSNDAVPTFIQERFIFIRETSHNAYRASSYVMSSLIVSLPFFALQAITFAAITKFIWLHLRGNLFRYWFILFASLVSTNAYVMLVSATVPSYVTGYAIVIATTALFFLTCGFFLKRGAIPWYWIWLHYISAIKYPFEALLVNEFKGNHCYNNDSVIPPAPGPMGNIKISSLHKSTNTTCGLIGQDVLSTMDIPDNSNVWIDVLILLGWGVLYRFLFYFVLRFYSKNERK |
| Medtru_MtrunA17_Chr5g0411621 | ***STR2*** | *Oryza sativa* | JN608806 | MAHRHVGDLEAAAYGGGRFGFTGGLEFTGLTYTVTKKQRGAGGEWEKKEVDLLHEVTGYAPKGCVTAVMGPSGAGKSTLLDALAGRIAARLGGRVALDGVEVSPGLVKRCSAYVMQDDRLFPMLTVRETLMFAADLRLGASVPAADKRRRVDALIDQLGLAASGNTYIGDEGTRGVSGGERRRVSIGVDIIHGPALLFLDEPTSGLDSTSAHSVVEKVRDIACAGSTVVLTIHQPSSRILQLLDHLVILARGQLMYSGAPREVAAHLGRMGRRVPKGESSIEHLLDVIQEYDQSEFGVAALAEFCLTGLKPPKLAADGISTVSSIPPTPLLAGEEDFDHSLRSQHSRSPWSAAAAAAATQFTPSRRPKKDHPEIVMGTPTPLSMSAYTLSEGDYRTPPPPRHAAAVTTLGGHRGKFANTYGGEVWVLMRRNFTNIWRTPELFLSRLMVLVAMGVLMATMFTKPRDDDQGVTERLSFFVFTVCVLFFSSNDAVPAFIQERFIFIRETSHNAYRASAYVVAGVVTYLPFLLLQSAAYAAIVWFALRLHGQFLYFLVMVYASLLSTNSFVVFISSVVPNFILGYAAVIAFTALFFLFCGYFLDSHSIPVGWKWMNTISTMKYPYEGLLMNEFQGGRVFSSHPPPAPPLTGDVILEHLKISTAEDRKWWMVLYLMGWAVFYRVLFYLVLRFASKNKRK |
| Medtru_MtrunA17_Chr5g0411621 | ***STR2*** | *Ginkgo biloba* | scaffold10016 | MLSGGLEFTDLTYTVTKKQKDLDGKWTSEEVDLLHFITGHAPKGSITAVMGPSGAGKSTFLDGLAGRIASGSLQGTVSLDGQEISPSLVKRTSAYVMQDDRLFPMLTVWETLMFAAEFRLGGSVSYSDKVMRVEKLIDQLGLTNARNTYIGNEETRGISGGERRRVSIGVDIIHGPALLFLDEPTSGLDSTSAHSVIEKVHDIAKDGSTVILTIHQPSYRVQMLLDHLIVLARGQLIYQGTPKDVAFHLGRMGRKVPKGENSIEYLLDVIQEYDRSPLTVTALAEYYRTGVKPPPITPLHQGLVSPNRFRSPRVKRAPGEENSQEAPIVPLWETRWVNNRIANDFDHSVRTPRSARGRGGSGVVESWRFTPSLQTPKRNKDRQTAPRSNSTSFSPGYYAYSADVISNTPTPHSSDYTVRESDYQTRWEDNSAKTDTDFGPKFSNSFHRELMVLIKRNFLNIRRTPELFLSRQMVLTIMGVMMATMFMNPSKDMQGITNLLSFFIFTVCIFFFSSNDAVPAFIQERFIFIRESSHNAYRASSYAISGLITYLPFLAIQALTYALLVWFALKLDGPFFYFVLILYVSLIATNSFVVFVSSVVPNYILGYAAVIAFTALFFLFCGYFLPRKSIPPYWIWMHWISTIKYPYEGLLMNEFSKDICYSRISQVCALGRKDILKHLSIDTNSSSKWASLSFLLGWAVVYRLLFYIVLRFASKNQRT |

**Supplemental Table 7:** Sequences and plasmids used for transcomplementation assays in *M. truncatula*
CCaMK-K kinases sequences were determined according to clustal Omega alignment with MtCCaMK-K (1 to 311 amino Acids).
All plasmids are available at https://www.ensa.ac.uk/

| ENSA L0 | Gene name | Species | Domesticated sequence |
|---|---|---|---|
| EC10384 | MedtruCCaMK | *Medicago truncatula* | ATGGGATATGGAACAAGAAAACTCTCAGATGAATATGAAGTTTCAGAAATTCTAGGTAGAGGTGGATTTTCTGTTGTTAGAAAAGGTACAAAAAAATCAAGCATTGAAGAAGAAAATCACAATCACAAGTAGCAATCAAAACCCTAAGAAGGTTAGGTGCTTCAAATAACCCTAGTGGATTACCAAGAAAAAAAGATATTGGAGAAAAAAGCACAATAGGGTTCCCTACAATGAGACAAGTTTCAGTTTCAGATACATTACTAACAAATGAGATACTTGTAATGAGGCGAATAGTCGAAAACGTTTCGCCACATCCAAATGTGATTGATCTTTATGATGTATATGAGGACACAAATGGTGTTCATCTTGTTCTTGAGCTTTGTTCCGGTGGTGAACTTTTCGATAGGATTGTTGCACAAGATAAGTATAGTGAGACTGAAGCTGCAACTGTGGTTCATCAAATAGCTTCAGGGTTAGAAGCTGTTCATAGAGCTAATATAGTTCATAGAGATTTGAAACCTGAAAATTGTCTTTTTTTAGATGTTAGGAAAGATTCTCCTCTTAAGATTATGGATTTTGGGTTGAGTTCTGTTGAAGAGTTTACTGATCCTGTTGTTGGTTTGTTTGGATCTATTGATTATGTTTCACCTGAGGCTCTTTCTCAAGGAAAGATTACTACTAAGAGTGATATGTGGTCCCTTGGGGTTATTCTATATATCTTACTTTCAGGGTATCCACCTTTCATTGCCCAAAATAATCGCCAAAAACAACAAATGATAATGAATGGGAATTTTAGTTTTTATGAGAAAACTTGGAAGGGAATTTCACAACCAGCAAAGAATTTGATTTCAAGTCTTTTAACCGTTGATCCTAGCAAGAGGCCTAGTGCTCTTGAGCTTCTAAGTGATCCATGGGTCAAAGGTGAGAAAGCCAAAGATGTTCAAATGGACCCTGAGATTGTCTCAAGGCTACAAAGCTTTAATGCAAGACGTAAACTTCGTGCAGCTGCAATTGCTAGTGTTTGGAGCTCCACAATCTTCCTTAGAACAAAAAAATTGAAATCATTGGTTGGATCCTATGATCTTAAAGAAGAGGAAATTGAAAATCTCAGGATGCATTTCAAGAAGATATGTGCAGATAGAGACAATGCAACTCTGTCAGAGTTTGAGGAGGTGTTAAAAGCAATGAATATGTTATCATTGATCCCTTTTGCTTCTCGTATATTTGATTTGTTTGACAACAACCGTGATGGAACAGTTGACATGCGTGAGATACTTTGTGGATTTTCCAGTCTCAAGAATTCCAAAGGAGAGGATGCTCTTCGTTTGTGCTTCCAGATGTATGATACAGATAGATCAGGCTGCATCAGCAAAGAGGAAGTAGCATCCATGCTCAGGGCTTTGCCATATGATTGTCTACCAACTGATATCACTGAACCTGGAAAATTGGATGAGATTTTTGACTTAATGGATGCTAATAATGATGGAAAAGTTACATTTGATGAATTCAAAGCTGCTATGCAAAGAGATAGCTCTCTTCAAGATGTAGTTCTCTCTTCTATTCGTCCATAA |
| EC10384 | gDNA-MedtruCCaMK | *Medicago truncatula* | ATGGGATATGGAACAAGAAAACTCTCAGATGAATATGAAGTTTCAGAAATTCTAGGTAGAGGTGGATTTTCTGTTGTTAGAAAAGGTACAAAAAAATCAAGCATTGAAGAAGAAAATCACAATCACAAGTAGCAATCAAAACCCTAAGAAGGTTAGGTGCTTCAAATAACCCTAGTGGATTACCAAGAAAAAAAGATATTGGAGAAAAAAGCACAATAGGGTTCCCTACAATGAGACAAGTTTCAGTTTCAGATACATTACTAACAAATGAGATACTTGTAATGAGGCGAATAGTCGAAAACGTTTCGCCACATCCAAATGTGATTGATCTTTATGATGTATATGAGGACACAAATGGTGTTCATCTTGTTCTTGAGCTTTGTTCCGGTGGTGAACTTTTCGATAGGATTGTTGCACAAGATAAGTATAGTGAGACTGAAGCTGCAACTGTGGTTCATCAAATAGCTTCAGGGTTAGAAGCTGTTCATAGAGCTAATATAGTTCATAGAGATTTGAAACCTGAAAATTGTCTTTTTTTAGATGTTAGGAAAGATTCTCCTCTTAAGATTATGGATTTTGGGTTGAGTTCTGTTGAAGAGTTTACTGATCCTGTTGTTGGTTTGTTTGGATCTATTGATTATGTTTCACCTGAGGCTCTTTCTCAAGGAAAGATTACTACTAAGAGTGATATGTGGTCCCTTGGGGTTATTCTATATATCTTACTTTCAGGGTATGTTTTTACTTTTCTTTCTTTCTTATATTTAAATTGTATTAGGTTTTATGTTTTGATTCTAGCGATGATGAGTTTGTATGTATCATCGTGGTTTTAACAAAACTCCACTTTAGATCTTCTACAAAATTACGGTGCTACTGTGATTTCGTCAAACTCAACATGTCCAAACATGTACTAGTAGTATGTTTAGTTTTAAATTTGGAATTTACAAACCTAACCTACAAGAAATAGCGCCGATTTGGGAACACGGTAGCTCACATCTCTCCGCGTGGTAGCTCGACTCTCCAAATTAATAGTGCGTTTGGTATTATGTTTGGAATTTCCAACCCAAACCTACAAGAAATAGTGTCTGTTTGGAAACATGTAGCTTACCTCTCACAGCGAGGTAGCTCATTACTTCAAGATGAACTTCCTCGAAGTGACATCATAAAAAAAGAAACCCAAACAAACACATGCTTAAGCTTTTGTTAGATACCTTGTGATTTCGTTAAACTCAACGTATCCAAACATATACTAGTGGTATGTTTAGTTTTAAAATTATTTTCATATAATTGTTATGCAGGTATCCACCTTTCATTGCCCAAAATAATCGCCAAAAACAACAAATGATAATGAATGTAAGTACCATCTCTTACAGAATATAGAAGTTCTTTCCACAATTACTTTGCTTACTAAAACAAAATGGTTTGGAACAGGGGAATTTTAGTTTTTATGAGAAAACTTGGAAGGGAATTTCACAACCAGCAAAGAATTTGATTTCAAGTCTTTTAACCGTTGATCCTAGCAAGAGGCCTAGTGCTCTTGAGGTATTGTACACTACTACTATCTTGAACATACCAACTTAACATTAAATGTCCACATAAATATTTTTTAATCGAGTTTGATGAAATTAGAATAATGATGTAGATATTTTATGACTTAAAGGACCAATTTATAACAAAAAAAACTTAAATGACTAACTTGCTATAATACAACTTAAGAATCGTAGATGATATTTGACCTATTTTTCATGACATTATCTTCAAATGTAAAGAAGTTTAAGGTGAATAAAAACCACAACACACATGTTGTTGTACTTTGTGTTTTTTGGAACAGCTTCTAAGTGATCCATGGGTCAAAGGTGAGAAAGCCAAAGATGTTCAAATGGACCCTGAGATTGTCTCAAGGCTACAAAGCTTTAATGCAAGACGTAAACTTCGTGCAGCTGCAATTGCTAGTGTTTGGAGCTCCACAATCTTCCTTAGAACAAAAAAATTGAAATCATTGGTTGGATCCTATGATCTTAAAGAAGAGGAAATTGAAAATCTCAGGATGCATTTCAAGAAGATGTAAGTACCCAAAAGTCAAAAATTATCATAGTAACATTTAACTTTAGCTCTTAAATTTCACAAGGAACCAAACATTTTAAGTAAATAATCAAATTAGTTAAGAATGACAGATAATTAAAATATTCAAAAGTATTCAGAACTCCTAGCTCAACTGGCAAAAATGTCGAAATTGTTAGGCCGAATGCCATAACCGGGGTTCTACCAAAAAAAAAAATATTCAAAAGTAATTTTCATAATATTTACACAATTCGAGCTAAGGTTTTAAAAAACAGTTCAAAGTCGCGTTTGAGATGTGACATCAATGTTTTTTTATGTCTTTGTAGTAATTACAATTACATTTGCTCGAAGTGTATTTTGACACAAAAAATTGGGACTATGTGTACGATACACTTCGGACTAATGTTTTCTTTTTGAATACAAAGAAAATACTAATAGAAAATACACCATTTCTGATTTTTCAATATTAACAAGTGTGTTGCAGTTGTAAACTTTGATAGTTTTGATTAAAGTAATGTAATGAGTTGTTAACAGATGTGCAGATAGAGACAATGCAACTCTGTCAGAGTTTGAGGAGGTGTTAAAAGCAATGAATATGTTATCATTGATCCCTTTTGCTTCTCGTATATTTGATTTGTTTGACAACAACCGTGATGGAACAGTTGACATGCGTGAGATACTTTGTGGATTTTCCAGTCTCAAGAATTCCAAAGGAGAGGATGCTCTTCGTTTGTGCTTCCAGGTGAATTATACTCAACACCCAAACAAATATTAAGTTTTACAAATATGAATATGTGAAGCAAATGGATTCAAATTATAGATATTTGTGAGTTGAAATTTATGTTGTTAGTATCAGACTAGATAGTTCGACTAGTTTAAACTAAATGTTAAGGAGTTTAGACATTGAATTTAAACAACACTAACGAAATAGTAAATATTAACCTAAAATTATGTATTTAACTTAGTTGATCAACATTGGGGCCTCATAACTTGTTAGACCTGTACATTCAAACCTTAGACTTAGTACAACATGATTTATTTTATTTTTCTAAGTCTTGATAAAACGTTCACTATATATTTAGGCTTAATTTTTTTTTGTCAAGTTTTACTTACCTTCAAGCTGAATTACGGATTATTAAAGTCTCGTTCACCTAAGAATCGAAGAGTTAACACCAAAATAAAATAAAAATTACTTTTGACCTTTAAATATGTTGGACTGAATCAATAAATTAGTTTTTTTTAGAGGTTGAACCAATAAATTAGTCTTTTTTTTAAGGAATAAATTAATGTCTCTAAAGAAATAGCTTAAAAATTAATGTTATGAACATACACTCTATAACTAGTTTTCAACATGTTAACCACATTTGCCTCTTTATTTTTATTTTGTTGGCAAATAGATGTATGATACAGATAGATCAGGCTGCATCAGCAAAGAGGAAGTAGCATCCATGCTCAGGGTAATGCACACACATCTCAATTAATTAGCTTCATCTTTTCACTTTCATTATATTAACACACATTTATACATTAATTAAACAGGCTTTGCCATATGATTGTCTACCAACTGATATCACTGAACCTGGAAAATTGGATGAGATTTTTGACTTAATGGATGCTAATAATGATGGAAAAGTTACATTTGATGAATTCAAAGCTGCTATGCAAAGAGATAGCTCTCTTCAAGATGTAGTTCTCTCTTCTATTCGTCCATAA |
| EC10390 | MedtruCCaMK-K | *Medicago truncatula* | ATGGGATATGGAACAAGAAAACTCTCAGATGAATATGAAGTTTCAGAAATTCTAGGTAGAGGTGGATTTTCTGTTGTTAGAAAAGGTACAAAAAAATCAAGCATTGAAGAAGAAAATCACAATCACAAGTAGCAATCAAAACCCTAAGAAGGTTAGGTGCTTCAAATAACCCTAGTGGATTACCAAGAAAAAAAGATATTGGAGAAAAAAGCACAATAGGGTTCCCTACAATGAGACAAGTTTCAGTTTCAGATACATTACTAACAAATGAGATACTTGTAATGAGGCGAATAGTCGAAAACGTTTCGCCACATCCAAATGTGATTGATCTTTATGATGTATATGAGGACACAAATGGTGTTCATCTTGTTCTTGAGCTTTGTTCCGGTGGTGAACTTTTCGATAGGATTGTTGCACAAGATAAGTATAGTGAGACTGAAGCTGCAACTGTGGTTCATCAAATAGCTTCAGGGTTAGAAGCTGTTCATAGAGCTAATATAGTTCATAGAGATTTGAAACCTGAAAATTGTCTTTTTTTAGATGTTAGGAAAGATTCTCCTCTTAAGATTATGGATTTTGGGTTGAGTTCTGTTGAAGAGTTTACTGATCCTGTTGTTGGTTTGTTTGGATCTATTGATTATGTTTCACCTGAGGCTCTTTCTCAAGGAAAGATTACTACTAAGAGTGATATGTGGTCCCTTGGGGTTATTCTATATATCTTACTTTCAGGGTATCCACCTTTCATTGCCCAAAATAATCGCCAAAAACAACAAATGATAATGAATGGGAATTTTAGTTTTTATGAGAAAACTTGGAAGGGAATTTCACAACCAGCAAAGAATTTGATTTCAAGTCTTTTAACCGTTGATCCTAGCAAGAGGCCTAGTGCTCTTGAGCTTCTAAGTGATCCATGGGTCAAAGGTGAGAAAGCCTAA |
| EC10390 | gDNA-MedtruCCaMK-K | *Medicago truncatula* | ATGGGATATGGAACAAGAAAACTCTCAGATGAATATGAAGTTTCAGAAATTCTAGGTAGAGGTGGATTTTCTGTTGTTAGAAAAGGTACAAAAAAATCAAGCATTGAAGAAGAAAATCACAATCACAAGTAGCAATCAAAACCCTAAGAAGGTTAGGTGCTTCAAATAACCCTAGTGGATTACCAAGAAAAAAAGATATTGGAGAAAAAAGCACAATAGGGTTCCCTACAATGAGACAAGTTTCAGTTTCAGATACATTACTAACAAATGAGATACTTGTAATGAGGCGAATAGTCGAAAACGTTTCGCCACATCCAAATGTGATTGATCTTTATGATGTATATGAGGACACAAATGGTGTTCATCTTGTTCTTGAGCTTTGTTCCGGTGGTGAACTTTTCGATAGGATTGTTGCACAAGATAAGTATAGTGAGACTGAAGCTGCAACTGTGGTTCATCAAATAGCTTCAGGGTTAGAAGCTGTTCATAGAGCTAATATAGTTCATAGAGATTTGAAACCTGAAAATTGTCTTTTTTTAGATGTTAGGAAAGATTCTCCTCTTAAGATTATGGATTTTGGGTTGAGTTCTGTTGAAGAGTTTACTGATCCTGTTGTTGGTTTGTTTGGATCTATTGATTATGTTTCACCTGAGGCTCTTTCTCAAGGAAAGATTACTACTAAGAGTGATATGTGGTCCCTTGGGGTTATTCTATATATCTTACTTTCAGGGTATGTTTTTACTTTTCTTTCTTTCTTATATTTAAATTGTATTAGGTTTTATGTTTTGATTCTAGCGATGATGAGTTTGTATGTATCATCGTGGTTTTAACAAAACTCCACTTTAGATCTTCTACAAAATTACGGTGCTACTGTGATTTCGTCAAACTCAACATGTCCAAACATGTACTAGTAGTATGTTTAGTTTTAAATTTGGAATTTACAAACCTAACCTACAAGAAATAGCGCCGATTTGGGAACACGGTAGCTCACATCTCTCCGCGTGGTAGCTCGACTCTCCAAATTAATAGTGCGTTTGGTATTATGTTTGGAATTTCCAACCCAAACCTACAAGAAATAGTGTCTGTTTGGAAACATGTAGCTTACCTCTCACAGCGAGGTAGCTCATTACTTCAAGATGAACTTCCTCGAAGTGACATCATAAAAAAAGAAACCCAAACAAACACATGCTTAAGCTTTTGTTAGATACCTTGTGATTTCGTTAAACTCAACGTATCCAAACATATACTAGTGGTATGTTTAGTTTTAAAATTATTTTCATATAATTGTTATGCAGGTATCCACCTTTCATTGCCCAAAATAATCGCCAAAAACAACAAATGATAATGAATGTAAGTACCATCTCTTACAGAATATAGAAGTTCTTTCCACAATTACTTTGCTTACTAAAACAAAATGGTTTGGAACAGGGGAATTTTAGTTTTTATGAGAAAACTTGGAAGGGAATTTCACAACCAGCAAAGAATTTGATTTCAAGTCTTTTAACCGTTGATCCTAGCAAGAGGCCTAGTGCTCTTGAGGTATTGTACACTACTACTATCTTGAACATACCAACTTAACATTAAATGTCCACATAAATATTTTTTAATCGAGTTTGATGAAATTAGAATAATGATGTAGATATTTTATGACTTAAAGGACCAATTTATAACAAAAAAAACTTAAATGACTAACTTGCTATAATACAACTTAAGAATCGTAGATGATATTTGACCTATTTTTCATGACATTATCTTCAAATGTAAAGAAGTTTAAGGTGAATAAAAACCACAACACACATGTTGTTGTACTTTGTGTTTTTTGGAACAGCTTCTAAGTGATCCATGGGTCAAAGGTGAGAAAGCCTAA |
| EC15199 | MedtruCYCLOPS | *Medicago truncatula* | ATGGAAGGGAGAGGATTTTCTGGTTTATACAAGAATTCAAGTGAGGAGTTATTCTTGAAAACAGTGATGGAGAGTCCTATTGGTATGCCGGTACCTACCATGGAGATGTTAGGATTCAAGACTGTTTCTCAAAGCTTTCGCACCGATAGTGAAGAGCTTTTCAAACGCTGGCTAACAAATGATCAAGAGGGATACAATTCATCAAGCATGGGACTTAACAGTCGTTTGTCGAAGAGAATATCAACTGAAATAGCTAATATGTCTAATCAACAACACATTGGTGTGGCTTCAGAAGGAAGAAACAATGATAAATCATGCTTACAAAATAACTTCTTGGCAAATGATGTTTCAAGTGATTTCAATTTTCCAATCAGAGATCCTGTTGATAGAGAATTGCAATCTAGTAACTTGTTTCTGGCCAAGGCCTGGTTTATTACCGATCAACGAATGACAAGAAGCCGGTCATCTGAATTGCGGCGAAGGTATACTGAAATGCAAAATTCTCAAGCACCACAAGGATTGGATTCTATGTTCATGGTTCCTGAGCATGATACTAACACTATAAAAGAAGAACTTGCAAATTTTAATGGGTTTGATTACCTTTCCATGTGTGAGTTACCAAGCCAAAAGGGCACATTCATGTCTCCATCCAACTCATCTTCGTCTACCTTCAACACACATCAATTGGTTGATGTAGATAAAGTTTCATCTTGTGTAAGTATGCTAAAAGGTACATTACAGCGCAAGAAACTCGAATGCCAAGTCGAGAAAGAAGCTGCAGAAGATGGCTTGAATGAAATATTTTGCATTCGAGAACCTCTTTTCCAATCAGCTTTTAATGAAGAAGAAAGTTGGAATCAACAAAAGCTAGTAAACGTTCAAGGAGATTTTACCGATCAAGTTAACGATCCCGGAGTCATGCAAACCCTTGAAGGAACCACAAACTTTGTCTTAGATGGTTTTGCAAATCAGACGAACCAAATACAAGGCAGAACAGCTTCTGGAGAACCCTCTCAAAGTGAATCTTCTGCTGCTGCACCAGTAATCTCATCTGGCTTAGATGCATGTGAAGGTCCCAGCAATTCAAATCAAACTCTTGGTGATAGCTCATGGAAACAAGTGGGAGAAAGCACTCAAAATAAAGTCAGAGGTGTCAGAGAACAGATAATGGATAATCTGAAAGATGACAGAAAGAGGAAAAGTCTAGAAAGATATGGATCTGTAACATCAGCTGTTTCAGATGGCAAGATGGATAACACAAAAAAGCGGAGGGTGGAGCGCTCAAGAAAAATGGCTGAAGCAAAGGAAAGAAATTTGACACCAACAATTCCCTCAGATATGCAAGCTATCTTGAAGCGATGCGAAAACCTTGAGAAGGAAGTTCGATCACTAAAGCTTAATTTGTCCTTCATGAATAGGAAGGATTCTGAACAAACAAAGCAGATAGAGGACCTTCAGAAGCAGAATGAGGACTTGGCGGATGAAAAAGAGCGCCTCCTCGAAGAGATTGAAAGAATTCTATCAGAAACTGGAAAGATTTGA |
| EC89030 | MimpudCCaMK | *Mimosa pudica* | ATGAGGATGGGAACCAGAAAGCTCTCAGATGAATATGAAGTTTCAGATATCTTAGGAAGAGGAGGATTTTCTGTTGTCAGAAAAGGCATTAAAATATCAGGCAGTGAAAAAACCCAAGTAGCCATAAAAACACTCAGAAGAGTAGGTAACTCTGATCATTTTGGAGTTCCAAAAACTCAGAAAGGTAATAAGAGCATAGCTTCCATGGCGTTTCCAACATGGAGACAAGTTTCAATATCAGATGCTTTACTCACCAATGAGATATTGGTCATGAGGAAGATAGTAGAGAATGTTTCACCACATCCAAATGTGATTGAACTCTATGATGTTTTTGAGGACTCAAATGGTGTTCATCTTGTACTTGAGCTTTGTTCCGGCGGCGAGCTGTTTGATCGAATTGTGGCACAAGAGAAGTACTCAGAGGCAGAAGCTGCAGCTGTAGTTCATCAGATTGCAAGAGGACTAGAGGCTATTCATGGAGCTAATATTGTGCATAGGGACTTGAAGCCTGAGAATTGCTTATTTCTGGACAGTAGAAAGGATTCTTCTCTAAAGATCATGGATTTTGGGTTGAGTTCTGTGGAGGAGTTTACTGACCCAATTGTTGGCTTGTTTGGATCAATTGATTATGTTTCACCAGAGGCTCTTTCTCAAGGAAAGATTACTAACAAGAGTGACATGTGGTCTGTGGGAGTGATTCTTTACATCTTACTCTCTGGGTATCCGCCTTTTATCGCTCAGAATAATCGCCAAAAGCAACAAATGATAATGACTGGAAGTTTCAGTTTCTATGAAAAGACATGGAAAACTATTTCCTCATCAGCAAAGCAATTGATTTCAAGTCTCCTAGCCGTTGATCCTCATAGAAGGCCTAGTGCCCAAGAGCTTCTCAGTCATCCATGGGTCAGAGGAGAAAAGGCTAAAGATGATCAGATGGATCCTGAGATTGTAGACAGATTACAAAAATTTAATGCTCGTCGCAAACTTCGTGCTGCTGCAATTGCTAGTTTATATAGTAGCACAGTTTTCTTGAGAATTAAAAATTTGAAATCCTTGTTAGGATCCTATGATCTTAAACAAGAAGAAATTGAAAGTCTCAAGATACACTTTAAGAAAATTTGTGCAAATGGAGACAATGCCACACTTCCTGAATTTGAGGAGGTGCTTAGAGCAATGAACATGTCATCACTTGTCCCTCTGGCACCACGTATATTTGACATATTTGATAATAATCAAGATGGAACAGTTGACATGCGTGAGATTCTTTGTGGATTTTCCAGCCTTAAGAATTCCAAGGGTGATGATGCTCTCCGTCTTTGCTTTCAGatgtatgacacagaccgatctgggtgcataactaaggaagaagtagcatctatgcttacaGCATTGCCAGAAGATTGTCTGCCAGCTGACATCTTTGAACCAGGCAAATTGGATGAGATATTTGACCTGATGGATGCCAACAGTGATGGCAAAGTCACCTTTGAAGAATTCAAAGCAGCTATGCTGAGAGACAGCTCTCTGCAGGATGTAGTCCTTTCTTCTCTCCGTCCATCATAG |
| EC89031 | MimpudCCaMK-K | *Mimosa pudica* | ATGAGGATGGGAACCAGAAAGCTCTCAGATGAATATGAAGTTTCAGATATCTTAGGAAGAGGAGGATTTTCTGTTGTCAGAAAAGGCATTAAAATATCAGGCAGTGAAAAAACCCAAGTAGCCATAAAAACACTCAGAAGAGTAGGTAACTCTGATCATTTTGGAGTTCCAAAAACTCAGAAAGGTAATAAGAGCATAGCTTCCATGGCGTTTCCAACATGGAGACAAGTTTCAATATCAGATGCTTTACTCACCAATGAGATATTGGTCATGAGGAAGATAGTAGAGAATGTTTCACCACATCCAAATGTGATTGAACTCTATGATGTTTTTGAGGACTCAAATGGTGTTCATCTTGTACTTGAGCTTTGTTCCGGCGGCGAGCTGTTTGATCGAATTGTGGCACAAGAGAAGTACTCAGAGGCAGAAGCTGCAGCTGTAGTTCATCAGATTGCAAGAGGACTAGAGGCTATTCATGGAGCTAATATTGTGCATAGGGACTTGAAGCCTGAGAATTGCTTATTTCTGGACAGTAGAAAGGATTCTTCTCTAAAGATCATGGATTTTGGGTTGAGTTCTGTGGAGGAGTTTACTGACCCAATTGTTGGCTTGTTTGGATCAATTGATTATGTTTCACCAGAGGCTCTTTCTCAAGGAAAGATTACTAACAAGAGTGACATGTGGTCTGTGGGAGTGATTCTTTACATCTTACTCTCTGGGTATCCGCCTTTTATCGCTCAGAATAATCGCCAAAAGCAACAAATGATAATGACTGGAAGTTTCAGTTTCTATGAAAAGACATGGAAAACCATTTCCTCATCAGCAAAGCAATTGATTTCAAGTCTCCTAGCCGTTGATCCTCATAGAAGGCCTAGTGCCCAAGAGCTTCTCAGTCATCCATGGGTCAGAGGAGAAAAGGCTTAG |
| EC89024 | DistriCCaMK | *Discaria trinervis* | ATGGGACAAGAAACAAAAAGACTCACAGATGAATTTGAAATCTCAGAGGTTTTAGGCAGAGGAGGATTCTCAGTTGTAAGAAAAGGGATAAGAAAATCATCGTCAGTGAATAACACCGATATCAAATCCCAAGTAGCAATCAAAACCCTGAAAAGGTTAGGACCCAGCTCATCGACATTTCCTCCTCCCAATTCTCGCGGATTTTCCGAGAAAACAAACAGGGGATTTCAATTGAGGAAGCAAGTTTCGGTATCAGATGCTCTGCTAACCAATGAGATCTTGGTGATGAGGAAGATAGTGGAAAATGTTTCACCGCACGAGAATGTGATCGACCTCTATGATGTATTCGAGGATCAAAATGGGGTCCATCTGGTGTTGGAGCTCTGCTCTGGCGGCGAACTGTTCGATAGGATTGTTGCGCAAGAAAGGTACAACGAGATCGGAGCCGCCGCGGTTGTCCGGCAGATTGCCAAAGGATTGGGAGCTCTTCACAGGGCTAATATCGTCCACAGGGATTTGAAACCCGAAAACTGCCTTTTCTTGAATACCAATCAGGATTCTTCGTTGAAGATCATGGATTTTGGATTGAGCTCTGTGGAGGAATTCACTGACCCTGTTGTTGGGTTGTTTGGTTCCATTGATTATGTCTCTCCGGAGGCTCTTTCTCAGGGAAATGTTACTTCCAAAAGTGATATGTGGTCTTTGGGGGTTATCTTGTACATCCTTCTTTCTGGGTATCCACCTTTCATTGCTCAGACAAATCGGCAAAAACAACAAATGATAATGGCTGGAGACTTCAGCTTTTATGAGAAAACATGGAAGAACATTTCTTCATCAGCAAAACAATTGATCAAGAGCCTCTTGACTGTTGACCCTCAAAGGAGGCCTAGTGCTCTAGAGCTTGTAGATCATCCATGGGTGAGAGGTGATTCAGCGAAGGAGGATCAAATGGACGCTGAGATAGTTTCCAGGCTGCAGAGTTTCAATGCCCGGCGCAAACTCCGAGCCGCCGCCATTGCTAGTGTTTGGAGCTCCTCCATTTTCTTAAGGACCAAAAAGCTCAAATCTTTGCTTGGTTCTCACGACCTCTCATCAGATGAAATTGAAAATCTCAGGGATCATTTCAAGAAAATATGTGCAAAGGGGGACAATGCCACTCTATCTGAGTTCGAAAAGGTTTTGAAAGCAATGAACATGTCGTCCTTAATCCCTTTGGCTCCTCGCATCTTCGACCTTTTTGACGACAATCGAGATGGAACTGTGGACATGAGAGAGATACTCTGCGGGTTTTCAAGTCTCAGAAAATCTCAAGGTGACGATGCTCTGCGCCTCTGCTTTCAGATGTATGATACAGATAGATCTGGATGCATCACTAAGGAAGAAGTAGCATCAATGCTAAGAGCCTTACCAGATGACTGTCTACCAGTAGATTTAACAGAACCAGGGAAATTGGATGAGATATTTGATAGAATGGATGCAAACAGTGATGGAAAAGTTACCTTCGATGAGTTCAAAGCTGCCATGCAGATAGACAGCTCTCTCAAAGACCTAGTCCTCTCTTCTCTTCGCCCTTCATAG |

| | | | |
|---|---|---|---|
| EC89025 | DistriCCaMK-K | *Discaria trinervis* | ATGGGACAAGAAACAAAAAGACTCACAGATGAATTTGAAATCTCAGAGGTTTTAGGCAGAGGAGGATTCTCAGTTGTAAGAAAAGGGATAAGAAAATCATCGTCAGTGAATAACACCGATATCAAATCCCAAGTAGCAATCAAAACCCTGAAAAGGTTAGGACCCAGCTCATCGACATTTCCTCCTCCCAATTCTCGCGGATTTTCCGAGAAAACA<br>AACAGGGGATTTCAATTGAGGAAGCAAGTTTCGGTATCAGATGCTCTGCTAACCAATGAGATCTTGGTGATGAGGAAGATAGTGGAAAATGTTTCACCGCACGAGAATGTGATCGACCTCTATGATGTATTCGAGGATCAAAATGGGGTCCATCTGGTGTTGGAGCTCTGCTCTGGCGGCGAACTGTTCGATAGGATTGTTGCGCAAGAAAGGTAC<br>AACGAGATCGGAGCCGCCGCGGTTGTCCGGCAGATTGCCAAAGGATTGGGAGCTCTTCACAGGGCTAATATCGTCCACAGGGATTTGAAACCCGAAAACTGCCTTTTCTTGAATACCAATCAGGATTCTTCGTTGAAGATCATGGATTTTGGATTGAGCTCTGTGGAGGAATTCACTGACCCTGTTGTTGGGTTGTTTGGTTCCATTGATTATGTCTC<br>TCCGGAGGCTCTTTCTCAGGGAAATGTTACTTCCAAAAGTGATATGTGGTCTTTGGGGGTTATCTTGTACATCCTTCTTTCTGGGTATCCACCTTTCATTGCTCAGACAAATCGGCAAAAACAACAAATGATAATGGCTGGAGACTTCAGCTTTTATGAGAAAACATGGAAGAACATTTCTTCATCAGCAAAACAATTGATCAAGAGCCTCTTGACTGT<br>TGACCCTCAAAGGAGGCCTAGTGCTCTAGAGCTTGTAGATCATCCATGGGTGAGAGGTGATTCAGCGTAG |
| EC89107 | FravesCCaMK | *Fragaria vesca* | ATGGGACAAGAAACTAGGAGACTCGCCGACGAGTATGAAGTGGCTGAGATTTTAGGGAGAGGAGGATTCTCAGTAGTGAGGAAAGGGATAAGCAGAAAAGAAGGTAGCAGTGAAAAAAACAATGTTGCCATCAAAACACTCAAAAGACCGTTTGGGGGTTCTTCCTCTTCCACATCAAAAAATTCTTCATCCAATCCTGGTTTGAGGCAGGGGTT<br>CCCTGTCCCGACTAGGAAACAAGTCTCGATATCTAATGTTTTGCTCACAAATGAAATCTTGGTCATGAGGAAGATCGTTGAGAATGTGTCGCCGCATCCGAATGTGATTGACCTCTATGATGTGTATGAGGATGAGAATGGTGTTCACCTTGTCCTGGAGCTTTGTTCTGGTGGGGAACTCTTTGATAGGATTGTGAAGGAAGAGAGGTACTCTGAG<br>GTAGGGGCTGCAGCTGTAGTCAGGCAGATTGCACAAGGTTTAGCTGCTCTGCACAAGTCAAATATTGTTCATAGGGATTTGAAGCCTGAGAATTGTCTGTTCTTGAATAGTTGTGATGATTCTCCTTTGAAGATTATGGATTTCGGACTCAGTTCTGTCGAGGAGTTTACTGACCCTGTTGTTGGCTTGTTTGGTTCCATTGATTATGTATCACCAGAG<br>GCTCTTTCTCAGGGACAAATCACTTGTAAAAGTGATATGTGGGCTCTTGGTGTAATCTTGTATATCCTTCTCTCCGGATACCCACCTTTTATTGCTCAGTCGAATCGGCAGAAGCAGCAGATGATAATGGCTGGAGAATTCAGCTTTTACGAGAAAACTTGGAAGGGAATTTCCTTATCAGCCAAGCAATTAATATCAGACCTCCTCAAAGTTGACCCT<br>GAAAAGAGGCCTAGTGCTCAAGAGCTTTTGGACCATCCATGGGTTGTAGGTCTTTCAGCCAGAGAGGATCAAATGGATGCTGAGATTGTGTCACGACTGCAGAGTTTTAATGCTCGACGCAAACTCCGAGCTGCAGCAATAGCTAGTGTGTGGACAACTTCTATTTTCTTGAGGACAAAGAAGCTCAAATCATTACTAGGGTCCCATGACCTTAAAC<br>AGGAGGAAATCCAGAACCTGAGTTTGCATTTTAAGAAGATATGCAAAATGGGTGATAATGCGACGCTGTCTGAATTTGAAGAGGTACTGAAAGCAATGAAAATGTCATCACTGGTCCCTTTAGCACCCCGTATTTTCGACCTATTTGACAACAATCGAGATGGAACAGTAGACATGCGAGAGATACTCTGTGGTTTCTCTAGTCTCAAGAATTCACA<br>AGGAGATGATGCTCTCCGCCTCTGCTTCCAGATGTATGATACAGATCGGTCGGGATGCATCAGTAAAGAAGAAGTGGCATCTATGTTGAGGGCTTTGCCAGATGACTGCCTACCAGCTGATATTACCGAACCAGGGAAGTTGGATGAAATTTTTGATCGAATGGATGCCAACAGCGACGGAAAAGTTACCTTTGATGAGTTCAAAGCCGCCATGCA<br>GAGAGATAGCTCTCTTCAAGATGCAAAACTGACGTTTGATGCTACGTGTCATATGTTCGAAAATGATCTGTGA |
| EC89108 | FravesCCaMK-K | *Fragaria vesca* | ATGGGACAAGAAACTAGGAGACTCGCCGACGAGTATGAAGTGGCTGAGATTTTAGGGAGAGGAGGATTCTCAGTAGTGAGGAAAGGGATAAGCAGAAAAGAAGGTAGCAGTGAAAAAAACAATGTTGCCATCAAAACACTCAAAAGACCGTTTGGGGGTTCTTCCTCTTCCACATCAAAAAATTCTTCATCCAATCCTGGTTTGAGGCAGGGGTT<br>CCCTGTCCCGACTAGGAAACAAGTCTCGATATCTAATGTTTTGCTCACAAATGAAATCTTGGTCATGAGGAAGATCGTTGAGAATGTGTCGCCGCATCCGAATGTGATTGACCTCTATGATGTGTATGAGGATGAGAATGGTGTTCACCTTGTCCTGGAGCTTTGTTCTGGTGGGGAACTCTTTGATAGGATTGTGAAGGAAGAGAGGTACTCTGAG<br>GTAGGGGCTGCAGCTGTAGTCAGGCAGATTGCACAAGGTTTAGCTGCTCTGCACAAGTCAAATATTGTTCATAGGGATTTGAAGCCTGAGAATTGTCTGTTCTTGAATAGTTGTGATGATTCTCCTTTGAAGATTATGGATTTCGGACTCAGTTCTGTCGAGGAGTTTACTGACCCTGTTGTTGGCTTGTTTGGTTCCATTGATTATGTATCACCAGAG<br>GCTCTTTCTCAGGGACAAATCACTTGTAAAAGTGATATGTGGGCTCTTGGTGTAATCTTGTATATCCTTCTCTCCGGATACCCACCTTTTATTGCTCAGTCGAATCGGCAGAAGCAGCAGATGATAATGGCTGGAGAATTCAGCTTTTACGAGAAAACTTGGAAGGGAATTTCCTTATCAGCCAAGCAATTAATATCAGACCTCCTCAAAGTTGACCCT<br>GAAAAGAGGCCTAGTGCTCAAGAGCTTTTGGACCATCCATGGGTTGTAGGTCTTTCAGCCTGA |
| EC16953 | HorvulCCaMK | *Hordeum vulgarum* | ATGTCAACAACTGAAAGCAGAAGGCTGTCCGACGACTATGAAGTCACAGATGTCCTCGGCCGAGGCGGTTTCTCCATAGTGAGAAGAGGTGTGAGCAAGTCTGATGAAGGAAAGACCCAGGTCGCGATAAAGACTCTCCGAAGGCTTGGCCCTGCGATGATGGGGATGCAGCAGGGGTCAAAGGCCGCCCCGAGCTCGGGAGGCCTCCCGCTG<br>TGGAAACAGGTATCCATCTCCGATGCGCTGCTCACTAACGAGATTCTTGTCATGAGGAGGATAGTGGAGAACGTCGCGCCGCATCCGAACGTCATCTGCCTGCATGACGTGTATGAGGATGCGCATGGTGTGCACCTCATCCTCGAGCTGTGCTCGGGTGGTGAGCTGTTCGACAGGATCATAGGGCGTGAGCGGTACTCGGAGTTCGATGCTGC<br>TGCTGTTGTCAGGCAGATTGCTAGTGGGTTGAAGGCTCTTCACAAGGCGAACATCATACACAGAGATCTGAAGCCAGAGAACTGCCTCTTCTCGGACAGAAAAGAAGATTCCACGCTGAAGATCATGGATTTTGGCTTGAGTTCTGTAGAAGATTTCAGTGACCCGGTTGTGGCGCTGTTTGGGTCGGTAGATTATGTTTCGCCAGAAGCACTCTC<br>GCGGCAAGAGGTTTCGACTGCTAGCGATATGTGGTCTGTTGGGGTGATTCTGTATATTCTTTTATCCGGATGCCCACCATTTCATGCTGCGACTAATCAAGAAAAGCAGCAAAGGATACTGCAAGGTGAATTTAGTTTTCAGGAACATACATGGAAAACAATAACTTCATCAGCCAAAGATCTGATTTCCAGTCTCCTTTCCGTCGAACCTTACAAAA<br>GGCCCACCGCAAGTGATCTCTTGATGCATCCTTGGGTGATAGGAGACTGCGCCAAGCAAGATCGCATTGATGCAGAGGTCGTTTCGAAACTGCAAAGGTTCAATGCTAGAAGGAAATTACGGGCGGCAGCAATAGCCAGCGTCCTGAGCAGCAAAGTGGCATTGAGGACAAAAAGGCTGAGGAGTCTTTTGGGAACCCATGACCTTAGCTCGGA<br>AGAGCTAGATAACTTGCGGCAACATTTTGCAAGGATATGTGCAGACGGAGAGAACGCCACGCTAGCAGAATTCGAGCAGGTGCTGAAAGCAATGAAGATGGACTCACTAATCCCTCTAGCTCCCCGTGTGTTTGATCTGTTTGACAACAACCGGGATGGAACAGTGGACATGAGGGAGATCCTCTGCGGGCTCTCGAGCCTGAGGAACTCGCGG<br>GGCGACGATGCGCTGCGACTCTGCTTCCAGATGTACGACGCGGACCGGTCGGGTTGCATCAGCAAGGAGGAGCTGACGTCGATGCTCCGGGCGCTGCCTGAGGAGTGCCTCCCGGGGGACATCGCGGAGCCGGGGAAGTTGGACGAGATGTTCGACACGATGGACGCCAACGGGGACGGCGAGGTGACCTTCGACGAGTTCAAGGCGGCGA<br>TGCAGAAGGACAGCGCCCTCCAAGACGTCGTCCTCTCCTCCCTGCGACCCGGACAGCAGTAG |
| EC16954 | HorvulCCaMK-K | *Hordeum vulgarum* | ATGTCAACAACTGAAAGCAGAAGGCTGTCCGACGACTATGAAGTCACAGATGTCCTCGGCCGAGGCGGTTTCTCCATAGTGAGAAGAGGTGTGAGCAAGTCTGATGAAGGAAAGACCCAGGTCGCGATAAAGACTCTCCGAAGGCTTGGCCCTGCGATGATGGGGATGCAGCAGGGGTCAAAGGCCGCCCCGAGCTCGGGAGGCCTCCCGCTG<br>TGGAAACAGGTATCCATCTCCGATGCGCTGCTCACTAACGAGATTCTTGTCATGAGGAGGATAGTGGAGAACGTCGCGCCGCATCCGAACGTCATCTGCCTGCATGACGTGTATGAGGATGCGCATGGTGTGCACCTCATCCTCGAGCTGTGCTCGGGTGGTGAGCTGTTCGACAGGATCATAGGGCGTGAGCGGTACTCGGAGTTCGATGCTGC<br>TGCTGTTGTCAGGCAGATTGCTAGTGGGTTGAAGGCTCTTCACAAGGCGAACATCATACACAGAGATCTGAAGCCAGAGAACTGCCTCTTCTCGGACAGAAAAGAAGATTCCACGCTGAAGATCATGGATTTTGGCTTGAGTTCTGTAGAAGATTTCAGTGACCCGGTTGTGGCGCTGTTTGGGTCGGTAGATTATGTTTCGCCAGAAGCACTCTC<br>GCGGCAAGAGGTTTCGACTGCTAGCGATATGTGGTCTGTTGGGGTGATTCTGTATATTCTTTTATCCGGATGCCCACCATTTCATGCTGCGACTAATCAAGAAAAGCAGCAaAGGATACTGCAAGGTGAATTTAGTTTTCAGGAACATACATGGAAAACAATAACTTCATCAGCCAAAGATCTGATTTCCAGTCTCCTTTCCGTCGAACCTTACAAAA<br>GGCCCACCGCAAGTGATCTCTTGATGCATCCTTGGGTGATAGGAGACTGCGCCTAG |
| EC11058 | ZeamayCCaMK | *Zea mays* | ATGTCTAAGACCGAGAGCAGGAAGCTGTCGGACGACTATGAGGTCGCGGAGGTCCTCGGCCGAGGCGGATTCTCGATCGTGAGGCGGGGAGTGAGCAAGTCAGAGGGGAAGGCGGCTCAACAGGTCGCCATAAAGACCCTGCGAAGGCTGGGCCCGGCCACCACGACCGGCGGGAGCACGCTGCCGCCGCAGCAAGGCTCAAATAAGGGC<br>GGCGGCCTGGTTCCGACGTGGAAGCAGGTGTCGTCCATCTCCGACGCGCTGCTCACCAACGAGATCCTCGTGATGCGGCGGATCGTGGAGGCCGTGGCGCCGCACCCCAACGTCATCGCGCTGCACGACGTTTACGAGGACGCGCGCGGCGTGCACCTGGTCCTGGAGCTCTGCTCGGGCGGCGAGCTGTTCGACCGGATCGTGGGGCGCGAC<br>CGCCACTACTCGGAGTTCGACGCGGCGGGCGTCGTCCGCCAGATCGCGCGCGGGCTGCAGGCGCTCCACGGCGCGGGCGTCGTGCACCGGGACCTCAAGCCCGAGAACTGCCTCTTCGCCGACAGGGGCGAGGCCTCCACGCTCAAGATCATGGATTTCGGCCTCAGCTCCGTCGAGGACTTCAGCGACCCCGTCGTCACCCTGTTCGGCTCGA<br>TCGACTACGTTTCGCCGGAGGCCCTCTCGAGGCAGGATGTCTCGGCTGCCAGCGACATGTGGTCTGTTGGGGTGATTCTGTATATCCTCTTGTCCGGATGCCCGCCGTTCCATGCACCAACTAATCGAGAGAAGCAGCAGAGGATACTGCAGGGAGAGTTCAGTTTCCAGGACCATACATGGAAAACAATCTCTTCATCAGCCAAAGAACTGATCT<br>CCTCTCTTCTCTCTGTTGAGCCTTACAAGAGGCCAACAGCAAGTGACCTGCTGGGGCATCCTTGGGTGATAGGGGACTGCGCGAAGCAGGACCTCATGGACGCGGAGGTCGTGACGAAGCTGCAGAGGTTCAACGCCAGAAGGAAGCTGCGGGCGGCGGCGATCGCCAGCGTGCTCAGCAGCAAGGTGGCGCTGCGGACGAAGCGGCTGCGG<br>AACCTGCTGGGACGGCATGACCTGAGCTCCGAGGAGCTCGACAACCTGCGGGCTCACTTTGCACGGATATGCGCCGACGGCGAGAACGCGACGCTGGCGGAGTTCGAGCAGGTGCTGAGGGCGATGAGGCTGGAGGCGCTGGTCCCGCTGGCGCCGCGCGTGTTCGACCTGTTCGACAACAACCGCGACGGCACGGTGGACATGAGGGAGAT<br>CCTGTGCGGGCTGTCCAGCCTCCGCAACTCCCGCGGCGACGACGCGCTCCGGCTCTGCTTCCAGATGTACGACGCCGACCGCTCCGGGTGCATAAGCAAGGACGAGCTGGCGTCGATGCTGCGAGCGCTGCCCGAGGAGTGCCTTCCGGGCGACATCACGGAGCCCGGGAAGCTGGACGAGGTGTTCGACGAGATGGACGCCAACGGCGACG<br>GCAAGGTGAGCTTCGAGGAGTTCAAGGCGGCCATGCAGAAGGACAGCGCCCTCCAGGACGTCGTCCTCTCGTCCCTCCGGCCGCCGCCGCCGCCATAG |
| EC11077 | ZeamayCCaMK-K | *Zea mays* | ATGTCTAAGACCGAGAGCAGGAAGCTGTCGGACGACTATGAGGTCGCGGAGGTCCTCGGCCGAGGCGGATTCTCGATCGTGAGGCGGGGAGTGAGCAAGTCAGAGGGGAAGGCGGCTCAACAGGTCGCCATAAAGACCCTGCGAAGGCTGGGCCCGGCCACCACGACCGGCGGGAGCACGCTGCCGCCGCAGCAAGGCTCAAATAAGGGC<br>GGCGGCCTGGTTCCGACGTGGAAGCAGGTGTCGTCCATCTCCGACGCGCTGCTCACCAACGAGATCCTCGTGATGCGGCGGATCGTGGAGGCCGTGGCGCCGCACCCCAACGTCATCGCGCTGCACGACGTTTACGAGGACGCGCGCGGCGTGCACCTGGTCCTGGAGCTCTGCTCGGGCGGCGAGCTGTTCGACCGGATCGTGGGGCGCGAC<br>CGCCACTACTCGGAGTTCGACGCGGCGGGCGTCGTCCGCCAGATCGCGCGCGGGCTGCAGGCGCTCCACGGCGCGGGCGTCGTGCACCGGGACCTCAAGCCCGAGAACTGCCTCTTCGCCGACAGGGGCGAGGCCTCCACGCTCAAGATCATGGATTTCGGCCTCAGCTCCGTCGAGGACTTCAGCGACCCCGTCGTCACCCTGTTCGGCTCGA<br>TCGACTACGTTTCGCCGGAGGCCCTCTCGAGGCAGGATGTCTCGGCTGCCAGCGACATGTGGTCTGTTGGGGTGATTCTGTATATCCTCTTGTCCGGATGCCCGCCGTTCCATGCACCAACTAATCGAGAGAAGCAGCAGAGGATACTGCAGGGAGAGTTCAGTTTCCAGGACCATACATGGAAAACAATCTCTTCATCAGCCAAAGAACTGATCT<br>CCTCTCTTCTCTCTGTTGAGCCTTACAAGAGGCCAACAGCAAGTGACCTGCTGGGGCATCCTTGGGTGATAGGGGACTGCGCGTAG |
| EC11033 | MarpalCCaMK | *Marchantia paleacea* | ATGTTGGAAGATTCGAACCACCCGGCCAGCTCGGGAGGAGTAGAATTTTATCATGTGGGGGAGAAGAGGAAGAGTCCGATGGTGGCGCTTGATCGGCCAGGGCAGCGCAGAGTAGATGACGATTACATAGTGGGCAAGGTACTGGGCAGCGGTGGATTCTCAGTGGTGCGCCAAGGCATGAACAGAGAAGATGGAGCGCTTGTGGCCATAAA<br>AACTTTGAAGAAACAGGGCATGGGAATGGGCTTCGGACCAAATGCCATCCACCAACAGTTTGCTGGTGGTGGAGGAGGGCGGGGAATAGGAGGTGGTGGAGTGAAACGGAAACTAGGAATGACAGCCAGCACAACAGAGTTGTCCATCTCGGAGGCTCTCATTGCCAACGAGATCATAGTCATGAAACGAATTGTCGAGGATGTTTCACCTCAT<br>GCGAACGTAGTTCATCTGCTGGATGTCTACGAAGATCCGACCGGTGTCCACCTAGCGCTCGAGTTGTGCTCGGGGGGTGAGCTCTTTGATCGCATTGTGTCTCAGGAACGATACTCCGAAGCAGGAGCTGCTGAAGTTATTCGTCAAATTGCCAGTGGCCTCGCCAGTCTACACCAAGCTCAAATTGTCCACCGTGACTTGAAACCAGAAAACTGCC<br>TCTTTCTTACTCCTGCTCAGGATTCACCTCTGAAGATCATGGACTTTGGTCTTAGCCACATAGATGGAGTCACGAGTCCAGTCGTTGGGATGTTTGGATCCATCGACTATGTTGCTCCAGAAGCTCTTCTTCGGAAAGCCGTGCTACCAGCCAGCGATATGTGGTCACTGGGGGTCATCTTGTACATTCTCCTCTGTGGGTATCCTCCTTTCCACGCAA<br>GAACAAACAGAGATAAGCAGAAGCTGATCCTCTCGGGTGACTTTGGTAACTTTGGAGAGTTCACGTGGAACAAAGTCACGTCATCTGCAAAGCAACTCATCGCTAGTTTATTGGCGGTGGATCCTCACCGCCGACCTTCTGCCGCAGAGTTACTACAACACGCATGGGTGAAAGGTGATGCTGCGAAAAGGGAGCCTGTGGGTCAAGATGTCATC<br>AAGCGTATACAATCTTTAAATGCACGTCGCAAATTCCGTGCTGCAGCATTTGCAAGTATCTTCAGTAGCAAGTTCCTCCTTCGGACCAAAAAGCTGAAAACTTTGGTCGGCTCGAATAATATTTTGAGTGAAGAAGATCTTGAAAATCTTCAAACACATTTCAAACGCATTTCAACAAACGGAGCAAGTGTGACGGTTTCAGAATTTAAGGATGTGCT<br>GAAGGCTATGAACTTGGCGAGTTTGGTACCATTGGCCCAACGCATTTTTGACCTCTTTGACGGCAACAAAGATGGTTGCGTGGACATGAGAGAGATTATTGTCGGATTTTCTTCTCTGAAAAAGTCGCACGGAGATGAGGCTCTGAGACTCTGCTTCCAGCTGTATGACACCGATCGCTCGGGCTTCATCTCACGAGAGGAACTGGCGTCCATGCTA<br>CGGGCATTACCTGAGGAGTACTTGCCACCGGACGTCACAGAGCCAGGGAAGCTAGATGAGATGTTTGATCGTATGGATGCTAATAACGATGGAAGGATAAGTTTTGAAGAGTTTAAGGAGGCCATACAAGACAACAATTTCTTGCAAGATGCCGTTCTTTTTCCCCTACGGCAGTCTCCTGTTTAG |
| EC11031 | MarpalCCaMK-K | *Marchantia paleacea* | ATGTTGGAAGATTCGAACCACCCGGCCAGCTCGGGAGGAGTAGAATTTTATCATGTGGGGGAGAAGAGGAAGAGTCCGATGGTGGCGCTTGATCGGCCAGGGCAGCGCAGAGTAGATGACGATTACATAGTGGGCAAGGTACTGGGCAGCGGTGGATTCTCAGTGGTGCGCCAAGGCATGAACAGAGAAGATGGAGCGCTTGTGGCCATAAA<br>AACTTTGAAGAAACAGGGCATGGGAATGGGCTTCGGACCAAATGCCATCCACCAACAGTTTGCTGGTGGTGGAGGAGGGCGGGGAATAGGAGGTGGTGGAGTGAAACGGAAACTAGGAATGACAGCCAGCACAACAGAGTTGTCCATCTCGGAGGCTCTCATTGCCAACGAGATCATAGTCATGAAACGAATTGTCGAGGATGTTTCACCTCAT<br>GCGAACGTAGTTCATCTGCTGGATGTCTACGAAGATCCGACCGGTGTCCACCTAGCGCTCGAGTTGTGCTCGGGGGGTGAGCTCTTTGATCGCATTGTGTCTCAGGAACGATACTCCGAAGCAGGAGCTGCTGAAGTTATTCGTCAAATTGCCAGTGGCCTCGCCAGTCTACACCAAGCTCAAATTGTCCACCGTGACTTGAAACCAGAAAACTGCC<br>TCTTTCTTACTCCTGCTCAGGATTCACCTCTGAAGATCATGGACTTTGGTCTTAGCCACATAGATGGAGTCACGAGTCCAGTCGTTGGGATGTTTGGATCCATCGACTATGTTGCTCCAGAAGCTCTTCTTCGGAAAGCCGTGCTACCAGCCAGCGATATGTGGTCACTGGGGGTCATCTTGTACATTCTCCTCTGTGGGTATCCTCCTTTCCACGCAA<br>GAACAAACAGAGATAAGCAGAAGCTGATCCTCTCGGGTGACTTTGGTAACTTTGGAGAGTTCACGTGGAACAAAGTCACGTCATCTGCAAAGCAACTCATCGCTAGTTTATTGGCGGTGGATCCTCACCGCCGACCTTCTGCCGCAGAGTTACTACAACACGCATGGGTGAAAGGTGATGCTGCGTAG |
| EC11036 | MarpalCYCLOPS | *Marchantia paleacea* | ATGTTGGAAGATTCGAACCACCCGGCCAGCTCGGGAGGAGTAGAATTTTATCATGTGGGGGAGAAGAGGAAGAGTCCGATGGTGGCGCTTGATCGGCCAGGGCAGCGCAGAGTAGATGACGATTACATAGTGGGCAAGGTACTGGGCAGCGGTGGATTCTCAGTGGTGCGCCAAGGCATGAACAGAGAAGATGGAGCGCTTGTGGCCATAAA<br>AACTTTGAAGAAACAGGGCATGGGAATGGGCTTCGGACCAAATGCCATCCACCAACAGTTTGCTGGTGGTGGAGGAGGGCGGGGAATAGGAGGTGGTGGAGTGAAACGGAAACTAGGAATGACAGCCAGCACAACAGAGTTGTCCATCTCGGAGGCTCTCATTGCCAACGAGATCATAGTCATGAAACGAATTGTCGAGGATGTTTCACCTCAT<br>GCGAACGTAGTTCATCTGCTGGATGTCTACGAAGATCCGACCGGTGTCCACCTAGCGCTCGAGTTGTGCTCGGGGGGTGAGCTCTTTGATCGCATTGTGTCTCAGGAACGATACTCCGAAGCAGGAGCTGCTGAAGTTATTCGTCAAATTGCCAGTGGCCTCGCCAGTCTACACCAAGCTCAAATTGTCCACCGTGACTTGAAACCAGAAAACTGCC<br>TCTTTCTTACTCCTGCTCAGGATTCACCTCTGAAGATCATGGACTTTGGTCTTAGCCACATAGATGGAGTCACGAGTCCAGTCGTTGGGATGTTTGGATCCATCGACTATGTTGCTCCAGAAGCTCTTCTTCGGAAAGCCGTGCTACCAGCCAGCGATATGTGGTCACTGGGGGTCATCTTGTACATTCTCCTCTGTGGGTATCCTCCTTTCCACGCAA<br>GAACAAACAGAGATAAGCAGAAGCTGATCCTCTCGGGTGACTTTGGTAACTTTGGAGAGTTCACGTGGAACAAAGTCACGTCATCTGCAAAGCAACTCATCGCTAGTTTATTGGCGGTGGATCCTCACCGCCGACCTTCTGCCGCAGAGTTACTACAACACGCATGGGTGAAAGGTGATGCTGCGTAG |

**Plasmids used in GoldenGate clonings**

| Name | Description |
|---|---|
| EC15251 | pL0M-PU-LjUBI1 |
| EC41414 | pL0M-T-35S |
| EC41744 | pL1M-ELE-2 |
| A3 | L1M-R1-pAtUBI-DSred-t35S |
| EC47811 | L1M-R2 vector |
| EC50507 | pL2V vector |
